# Comparative Study of State-Dependent Cholesterol Binding Sites in Adenosine A_2A_ and A_1_ Receptors Using Coarse-Grained Molecular Dynamics Simulations in Biologically Relevant Membranes

**DOI:** 10.1101/2022.09.28.509875

**Authors:** Efpraxia Tzortzini, Robin A. Corey, Antonios Kolocouris

## Abstract

In this study we used coarse-grained molecular dynamics (CG MD) simulations to study protein-cholesterol interactions for different activation states of the A_2A_ adenosine receptor (A_2A_R) and A_1_ adenosine receptor (A_1_R). Three different membranes were used, of which a plasma mimetic membrane best matched cholesterol binding sites previously detected in X-ray structures and computationally for the inactive state of A_2A_R. Here, cholesterol binds stably between TM5 and TM6 in the intracellular leaf and between the extracellular part of TM6 and TM7. The stability of the identified binding sites to A_1_R with CG MD simulations were further investigated using potential of mean force calculations combined with umbrella sampling.

Cholesterol binding to certain cavities in active or inactive state can stabilize a conformation of active or inactive state respectively and is important to identify these cavities. For the active state the cholesterol binding sites have much longer residence times compared to the inactive state for both A_2A_R and A_1_R.

We observed for the active A_1_R two cholesterol binding sites in the extracellular membrane leaf in a cavity between TM2/TM3 and along TM3 in the intracellular leaf. We found a partially overlapped binding area between TM6 and TM7 in the extracellular membrane leaf for active state and inactive A_1_R state which can be antagonized by cholesterol molecules although the site has a much longer residence time for active A_1_R. For the active A_2A_R we predicted a high residence time cholesterol position between TM1 and TM7 in the middle region of TMD which is different from the binding sites calculated in active or inactive A_1_R and for the inactive A_2A_R. For the inactive A_2A_R cholesterol binds to the same extracellular area between TM6 and TM7 as in the inactive A_1_R.

The differences in stable, long residence time cholesterol binding sites between active and inactive states of A_1_R and for A_2A_R can be important for functional activity and orthosteric agonist or antagonist affinity and can be used for the design of allosteric modulators which can bind through lipid pathways.

## Introduction

G protein-coupled receptors constitute a superfamily of receptors which in humans counts more than 800 members. They are formed by seven transmembrane (TM) α-helices that detect and transduce diverse external stimuli across the cell membrane, the diversity of which is particularly high for class A GPCRs. ^1^ The most common activation pathway is by an agonist ligand binding in the orthosteric binding pocket located within the upper half of the TM domain (TMD) core. Given their numerous physiologic and regulatory functions, over 130 GPCRs are therapeutically targeted by more than one-third of approved drugs. ^2,3^

Adenosine receptors (ARs) are GPCRs that mediate the action of adenosine ^4^ and include four subtypes; A_1_, A_2A_, A_2B_ and A_3_. A_2A_R and A_2B_R subtypes act synergistically with the G_s_ proteins resulting in the stimulation of the adenylyl cyclase, and therefore, the increase of cAMP levels. In contrast, A_1_R and A_3_R subtypes inhibit the adenylyl cyclase and decrease cAMP levels within cell by coupling to G_i_ or G_q_ proteins. ^5,2^ A_2A_R has been extensively studied over the last few decades and its complexes with agonists, like adenosine or 5’-N-ethylcarboxamidoadenosine (NECA), ^6,7^ and several antagonists 8–12 have been solved with X-ray crystallography. The structure of A_2A_R in a complex with G_s_ protein and an agonist has been solved with cryogenic electron microscopy (cryo-EM). ^13^ Of the four AR subtypes, A_2A_R was the only one with a structure until 2017, when the structure of A_1_R in complex with antagonists was solved using X-ray crystallography ^14,15^ and the structure of the active state (as complex with G_i_ protein and agonist ^16^ or with G_i_ protein, agonist and positive allosteric modulator ^17^) using cryo-EM. ^16,17^ The comparison between the experimental structures of the inactive, ^8–12^ intermediate active ^6,7^ and active states ^13^ of A_2A_R, with Protein Data Bank (PDB) IDs, e.g. 4EIY, ^10^ 2YDO ^18^ and 5G53, ^13^ respectively, as well as the investigation of such structures with molecular dynamics (MD) simulations ^19^ have revealed the conformational changes that occur during receptor activation by bound agonists. For all the ARs these conformational changes resulted in the opening of an intracellular binding pocket that is achieved by a large outward pivotal tilt of TM6 accompanied also by movements of the TM5, TM7 helices and adjustments in extracellular loops and in the ligand binding pocket.

Cholesterol, an essential building block for membrane biogenesis, is also a signaling molecule that regulates embryonic development and numerous physiological processes by acting either as a ligand or as a precursor for oxysterols and steroid hormones. Aberrant amounts and activity of cholesterol are associated with pathological conditions such as obesity, atherosclerosis, infertility, and cancer, making its accurate sensing essential. ^20^ Being highly abundant in the cell membrane (34% of the total lipid content in mammalian plasma membranes), cholesterol is known to regulate GPCRs conformation, stability, and function 21–23 and may be a prime regulator of GPCRs, keeping their basal activity low by stabilizing their inactive or intermediate active conformation.

These effects can be due to specific molecular interaction between cholesterol and GPCR causing an allosteric modulation. ^24^ The first structural evidence for site-specific CHOL binding in GPCRs was provided by early X-ray structures of β_2_AR ^25^ and A_2A_R, ^10^ in which CHOL binds in distinct TM locations. Much progress has been made since, and now there are hundreds of GPCR structures available ^26^ solved by X-ray diffraction and several A_2A_R-antagonist complex structures have been solved with three or four cholesterol molecules bound to the extracellular part of the receptor in the regions TM2-EL1-TM3, TM5-EL3-TM6, TM6-TM7 (see relevant PDB IDs in refs 10,27–35,78). These structures gave rise early to the concept of CHOL-binding motifs and to the expectation that such motifs pervade GPCR surfaces. Specific binding sites can have high CHOL affinity for the receptor at specific positions, that may constitute a favorable environment for lower-affinity and annular CHOL molecules as shown for β_2_AR using thermostability and STD NMR experiments. ^37^ It was suggested that general binding sites existed, e.g. Cholesterol Consensus Motif (CCM), ^38,39^ comprising four amino acid side chains distant in primary sequence [4.39-4.43(R,K)]–[4.46(I,V, L)]– [4.50(W,Y)]– [2.41(F,Y)] (according to Ballesteros-Weinstein ^40^ nomenclature). Multiple sequence alignments suggested that this CCM extends far beyond β_2_AR to include as many as 44% of human GPCRs. ^21^ However, even if CCM motif is conserved in 44% of human class A GPCRs, it doesn’t always correlate with cholesterol binding sites observed in high resolution crystal structures or molecular dynamics (MD) simulations. Cholesterol Recognition Amino Acid Consensus (CRAC) motifs were also identified as contiguous residue sequences localized to single TM helices. ^23,41–45^ CRAC motifs have the sequence -L/V–(X)_1-5_–Y/F–(X)_1-5_–R/K-determined based on the sequence or in a crystal structure and suggested to stabilize GPCRs in the bilayer. It was suggested that cholesterol can also bind the reverse sequence known as “CARC” motif, i.e., -R/K–(X)_1-5_–Y/F–(X)_1-5_–L/V-. ^46^ A CRAC motif has been observed in TM7, co-localized with the highly conserved NPxxY motif, found conserved in 38% class A GPCRs. ^47^ However, the presence of the CRAC motif in the sequence is not predictive for cholesterol binding sites as illustrated by a recent work on CCK1R and CCK2R. ^47^

It has been suggested that cholesterol binding sites to GPCRs cannot be predicted based on recurring, generalizable motifs, determined based on the sequence or on the crystal structure, like the CCM and CRAC/CARC motifs and cholesterol preferential localization is due to rather specific molecular interactions between cholesterol and GPCR causing an allosteric modulation has also been evidenced, also called “cholesterol hot-spots” that can be specific for each receptor families. ^24,48–50^ It has been recently shown that cholesterol shifts the intermediate active–active equilibrium to the intermediate active by filling cavities required to be suppressed for activation realized by the agonist isoprenaline-β_1_AR complex to the active conformation. ^51^ The identified cholesterol binding sites are rather diverse across different GPCRs, ^49^ even for closely related GPCRs such as the β_1_ and β_2_ARs, 52–54 but it seems that there are two general aspects of the cholesterol action on GPCRs: (1) by filling cavities, cholesterol can reduce the total volume available for functional motions and (2) cholesterol ties together hydrophobic parts of the receptor, thereby stabilizing a particular conformation.

MD simulations have provided structural insights into GPCR-lipid interactions 55–57 and have identified multiple cholesterol binding sites on the surface of GPCRs, the occupancy of which resulted in increased conformational stability, suggested for A_2A_R 58–64 and β_2_AR. ^65^ Compared to the all-atom (AA) MD simulations, coarse-grained MD (CG MD) simulations allow for much longer simulations of proteins in membranes and more efficiently sample the diffusion of lipids, providing an unbiased picture of the interactions of lipids with membrane proteins, ^50^ e.g. with A_2A_R ^61,22^ or β_2_AR. ^22,47^ CG MD simulations have shown a good correlation between cholesterol localization found in the crystal structure of β_2_AR and the residence time calculated with the simulations. ^47^

Although many studies have focused on CHOL binding to A_2A_R, using experimental, functional or AA MD and CG MD simulations in POPC – cholesterol ^58,60,61,64^ or DPPC – DOPC – cholesterol ^60^ or a plasma mimetic membrane, ^63^ no studies have been performed for A_1_R. Different therapeutic applications have been identified in preclinical and clinical studies for A_1_R antagonists as potassium-sparing diuretic agents with kidney-protecting properties ^66^ for the treatment of chronic heart diseases, ^67^ for chronic lung diseases such as asthma, ^68^ for cognition enhancement, ^66^ and for treatment of dementia and anxiety disorders. ^69,70^ A_1_R agonists have been considered for the treatment of glaucoma and ocular hypertension, ^71^ arrhythmias, ^72^ and heart failure. ^73^

In this study, we employed CG MD simulations to characterize the interactions of cholesterol with A_1_R in the inactive state and full active state, embedded in model membranes including a plasma-mimetic membrane, in order the simulations to be more biologically realistic. ^63^ There is also an increasing understanding of the complexity of lipid membranes, the diversity of such complexity, and the impact this lipid complexity has on membrane function, particularly on protein – lipid interactions. ^77,55,78^ We explored the stability of cholesterol binding sites by its long residence time with relevant amino acid residues (we plotted only residues with residence time > 1 μs) and by potential of mean force using umbrella sampling (US/PMF) calculations. ^79^ To examine the accuracy of the applied computational models, we also performed CG MD simulations for the active and inactive state of A_2A_R embedded in the same membranes, and compare our predictions for the cholesterol binding sites from experimental structures, 26–34,78 and previous AA and CG MD simulations results. ^58–64^

We identified the previously detected experimentally or computationally binding sites of inactive A_2A_R. Thus, for the inactive A_2A_R we detected that cholesterol binds stably in the cleft between TM6 and TM7 in the extracellular leaf and in the inner leaf in TM5/TM6 area which are binding sites previously identified with AA and CG MD simulations ^61^ and have also been previously observed in crystal structures of inactive A_2A_R. ^10,27^ While it has been shown that cholesterol is a stabilizer of A_2A_R ^82^ it has been shown that depletion of cholesterol attenuates signaling of A_2A_R, coupled to G_αs_. ^82^

We calculated that in the inactive A_1_R cholesterol binds to the extracellular area between TM6 and TM7. We showed that cholesterol binds to active A_1_R to a partially overlapped position between TM6, TM7 residues with much longer residence time compared to the inactive A_1_R. We observed for the active A_1_R two additional cholesterol binding sites in the extracellular membrane leaf in a cavity between TM2/TM3 and TM3. For the active A_2A_R, we predicted a high residence time cholesterol position between TM1 and TM7 in the middle region of TMD which is different from the binding sites calculated in active or inactive A_1_R and for the inactive A_2A_R. The differences in long residence cholesterol binding sites between active and inactive states of A_1_R and for A_2A_R can be important for the predominance of the relevant conformation, effect in functional activity and can be used for the design of selective allosteric modulators. Indeed, membrane lipids have been shown to actively participate in the access and binding of ligands to transmembrane allosteric sites of GPCRs. ^83,84^

## Methods

### Protein modelling

For this study we used the experimental structures of A_1_R in the agonist-Gi-bound, active form (PDB ID 6D9H ^16^) and the antagonist-bound, inactive form (PDB ID 5UEN ^14^). We also used the experimental structures of A_2A_R in the agonist-mini-Gs, active form (PDB ID 5G53 ^13^) and antagonist-bound, inactive form (PDB ID 3EML ^85^). For each protein we added missing sequences using the SWISS-MODEL server ^86^ and embedded each protein structure in membrane with the Positioning of Proteins in Membranes (PPM) server ^87^ to obtain the Orientation of Proteins in Membrane (OPM). ^87^ We deleted antagonists or agonists and Gi or mini-Gs proteins, and then optimized the protein model using the Protein Preparation Wizard implementation in Schrödinger suite (Protein Preparation Wizard 2015-2; Schrödinger, LLC, New York, 2015) by performing a restrained energy minimization using the OPLS2005 force field ^88^ and assuming as convergence criterion for heavy atoms a Root Mean Square Deviation (RMSD) of 0.3 Å. The Define Secondary Structure of Proteins (DSSP) program, ^89,90^ which is a hydrogen bond estimation algorithm, was used to describe the secondary structure of both A_1_R and A_2A_R. AA models were converted to the CG Martini 2.2 force field ^91^–^93^ using the *martinize*.*py* script. In this procedure, secondary structure determined by DSSP was used to assign CG atom types and bonded interactions. Default position restraints were applied to the protein’s backbone (force constant of 1000 kJ mol^-1^ nm^-2^). Protein’s termini were set neutral using the *martinize*.*py* script and its cut-off distance scheme. The Martini2.2 and ElNeDyn22 force fields were applied in combination with stiff elastic network using an elastic network force constant of 500 kJ mol^-1^ nm^-2^ and upper distance cutoff of 10 Å.

### System Setup

Using the *insane*.*py* ^94^ script (available on the Martini website), for setting up bilayer systems using Martini 2.0 lipids, ^91^–^93^ we prepared three different membranes in which the CG model of active or inactive states of A_2A_R or A_1_R were embedded. Namely, one membrane consisting of 1:4 cholesterol:POPC and one membrane with POPC : cholesterol : phosphatidylinositolbisphosphate (PIP_2_), with the extracellular leaflet having 25% cholesterol and the intracellular leaflet having 25% cholesterol and 10% PIP_2_. We also used a plasma mimetic membrane (Table S1). The extracellular leaf of the plasma mimetic membrane is enriched in phosphatidylcholine (PC) lipids while the intracellular leaf is enriched in phosphatidylethanolamine (PE) lipids and phosphatidylserine (PS) lipids. Thus, the upper leaf is consisting by 4:4:1:1:3:2:5 cholesterol : 1,2-dioleoyl-sn-glycero-3-phosphocholine (DOPC) : 1,2-dioleoyl-sn-glycero-3-phosphoethanolamine (DOPE) : 1-palmitoyl-2-oleoyl-sn-glycero-3-phosphoethanolamine (POPE) : sphingomyelin (SPH) : monosialodihexosylganglioside (GM3) : cholesterol while the lower leaf is consisting by 1:1:4:4:2:1:2:5 POPC : DOPC : POPE : DOPE : 1-palmitoyl-2-oleoyl-sn-glycero-3 phosphoserine (POPS) : 1,2-dioleoyl-sn-glycero-3-phosphoserine (DOPS) : PIP_2_ : cholesterol. A description of the CG MD simulations systems is shown in Table 1.

**Table 1.** CG MD simulations performed with A_1_R and A_2A_R in model membranes for a total 300 μs simulation time. ^*a*^

| Protein<br>Membrane | A<sub>1</sub>R<br>active<br>state | A<sub>1</sub>R<br>inactive<br>state | A<sub>2A</sub>R<br>active state | A<sub>2A</sub>R<br>inactive<br>state |
| --- | --- | --- | --- | --- |
| POPC – cholesterol | 10μs × 3<br>repeats | 10μs × 3<br>repeats | 10μs × 3<br>repeats | 10μs × 3<br>repeats |
| POPC – cholesterol – PIP<sub>2</sub> | 10μs × 3<br>repeats | 10μs × 3<br>repeats | - | - |
| Plasma mimetic | 10μs × 3<br>repeats | 10μs × 3<br>repeats | 10μs × 3<br>repeats | 10μs × 3<br>repeats |

<sup>a</sup> (2 proteins x 3 membranes x 3 repeats) + (2 proteins x 2 membranes x 3 repeats) = 30 datasets with total simulation time 30 x 10 μs = 300 μs

### CG MD simulations with Gromacs

All CG MD simulations were performed using Gromacs2018.8 software. ^95,96^ Periodic boundary conditions were used in all axes. First, 5000 steps of energy minimization were applied using the steepest decent algorithm. During the energy minimization the neighbor list and long-range forces were updated in each step using the grid method. Short range forces were calculated using the cut-off scheme and long range electrostatic interactions were treated with the Particle Mesh Ewald (PME) method. ^97^ The cut-off distance for the electrostatic and Lennard-Jones interactions was set to 12 Å. After the energy minimization, two runs of 100 ns were performed in the (isothermal-isobaric) NPT ensemble to allow for system equilibration. For the non-bonded interactions, a cut-off scheme was used, with electrostatic forces shifted to zero in the range of 10–11 Å and with Lennard-Jones interactions shifted to zero in the range of 9–11 Å. Long range electrostatic interactions were treated with the PME method. ^97^ A time step of 10 fs was used with neighbor lists updated every 20 steps. The temperature was held constant at 310 K using the velocity rescale thermostat ^98^ with a coupling constant of 1.0 ps and pressure being semi-isotropically controlled using the Berendsen barostat ^99^ at 1 bar with a coupling constant of 4.0 ps and compressibility of 1 × 10^−5^. Then, a production run of 10 μs in the NPT ensemble was performed for each system in triplicate, with random velocities for each repeat with temperature held constant at 310 K using the velocity rescale thermostat ^98^ with a coupling constant of 1.0 ps while pressure was semi-isotropically controlled by the Parrinello-Rahman barostat ^100^ at 1 bar with a coupling constant of 12.0 ps and compressibility of 3 × 10^−4^. A time step of 20 fs was used with neighbor lists updated every 10 steps.

### Analysis of CG MD simulation trajectories

From the produced 10 μs CG MD simulations we discarded the first 2 μs and used the remaining 8 μs for the analysis. The visualization of the CG MD simulation trajectories and the generation of all figures were carried out using VMD 1.9.4. ^101^

Cholesterol binding sites, residence times/frequencies as defined in ref. ^102^–^105^ were calculated with the PyLipID python package (https://github.com/wlsong/PyLipID). ^106^ To avoid the rattling-cage effect in lipid binding a dual-cutoff scheme was used such that the interaction started when a cholesterol moves to a given residue closer than 5.5 Å (interaction initiation) and ended when cholesterol moves farther than 10 Å (termination of interaction). We identified the cholesterol binding sites from its contacts with A_1_R or A_2A_R using the graph theory and community analysis methods available in the PyLipID package. ^107^ Briefly, the graphs were implemented using protein residues as nodes and the frequency with which a given pair of residues interacted with cholesterol as edges. Each calculated community was considered as a candidate binding site. More details on the community method can be found at: https://pylipid.readthedocs.io/en/master/index.html. The cholesterol residence time in a binding site was calculated from the normalized survival time-correlation function *σ* (*t*) according to equation (1)

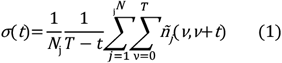

where *T* is the total simulation time, *N*_*j*_ is the total number of cholesterols, and *ñ*_*j*_ *(v, v*+*t*) is a function that takes the value 1 if cholesterol occupies the given binding site for a duration of time, *t*, after forming the contact at time *v*, and takes the value 0 if otherwise. The calculated residence times are converged if the measure of the goodness of fit of the biexponential function R^2 102,103^ is higher than 0.87 in all cases and (*k*_off_ – *k*_off,bootstrap_) is converged; *k*_off_ values were extracted from the biexponential fitting of the normalized survival function. To this end, the occupancy-based “hot” sites are polished with an implementation of network analysis previously used for protein–cholesterol interactions, ^104,105^ wherein simultaneously interacting protein residues are recorded and clustered according to defined binding sites for cholesterol.

### PMF calculations

To study the energetics of cholesterol binding sites of interest for both states of the A_1_R receptor we performed US/PMF calculations of the energetics of the free energy landscape for the interactions of cholesterol with the binding sites identified on the receptor. Steered MD (SMD) simulations ^108^ were carried out on the identified bound cholesterol sites in the complex lipid environment from the non-biased CG equilibrium simulations in order to generate the configurations for US of cholesterol-protein complexes. Thus, the bound cholesterol molecules were pulled away from the receptor in the membrane plane in a direction defined by the vector between the centre of mass (COMs) of the receptor and of the bound lipid. A rate of 1 nm/ns and a force constant of 1000 kJ/mol/nm^2^ was used. The starting configurations of the US were extracted from the SMD trajectories spacing 0.05 nm apart along the reaction coordinate. 36 US windows were generated, and each was subjected to 1μs MD simulation, in which a harmonic restrain of 1000 kJ/mol/nm^2^ was imposed on the distance between the COMs of the receptor and the bound lipid to maintain the separation of the two.

## Results

### A_2A_R in POPC - cholesterol membrane

The CG MD simulations-derived density maps of cholesterol binding showed that there are differences in cholesterol binding between the two states of A_2A_R in POPC – 20% cholesterol membrane (Figure S1). When plotting the residence time of cholesterol at each residue (as defined in the Methods Section) for A_2A_R in the active conformation (Figure 1A), we identified one key region centered at residue 95 with cholesterol residence times of ca. 1 μs, as well as three regions with residence time ca. 500 ns (centered at residues ca. 125, 190, 270). For the inactive state of the A_2A_R, we observed that, compared to the active state, cholesterol binding frequency and the number of binding regions are increased (Figure 1B and Figure S2). We observed (Figure 1B) five regions with cholesterol residence time of ca. 1-1.5 μs, (centered at residues ca. 15, 60, 190, 245, 270).

**Figure 1.**
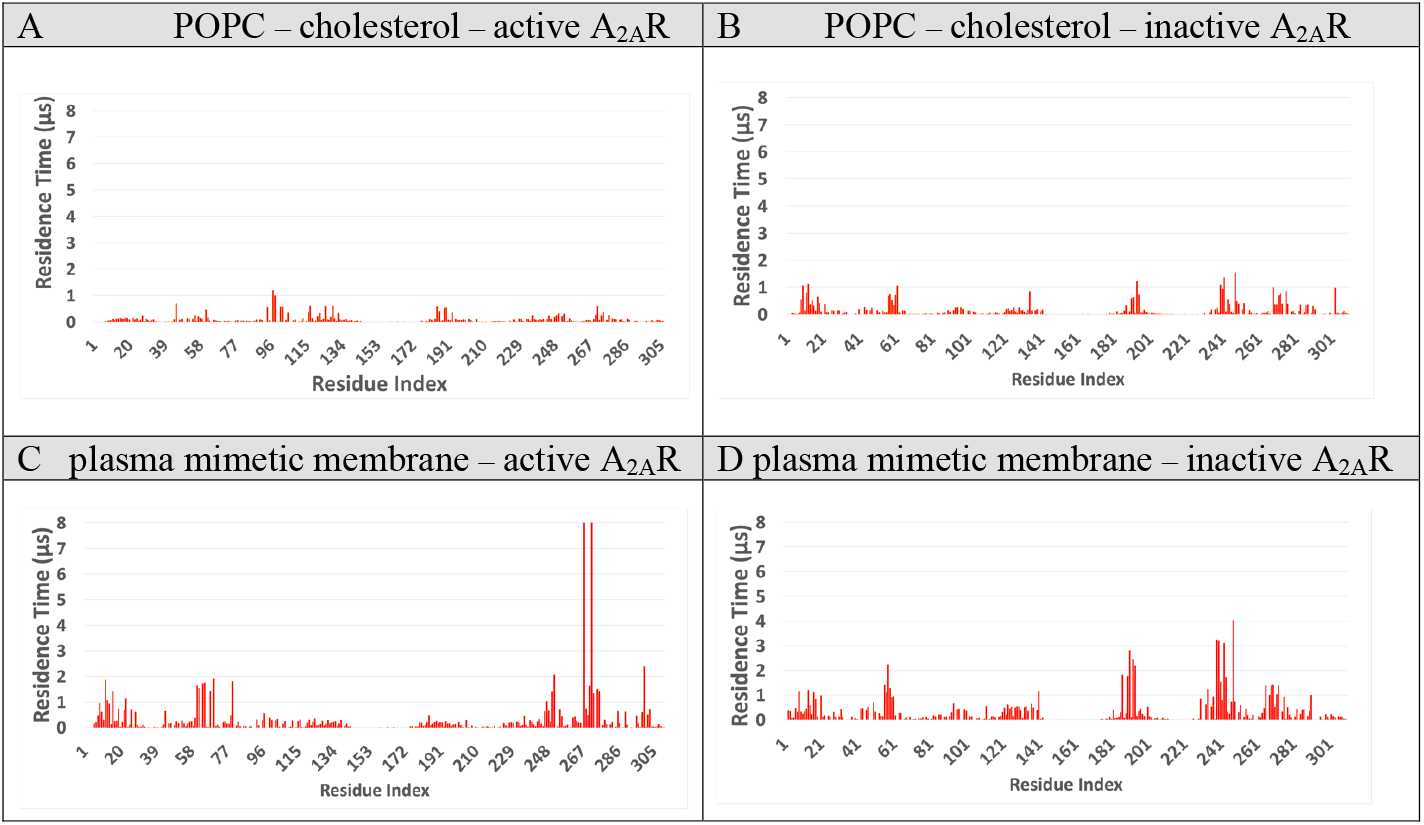
Residence time of cholesterol averaged for three repeats using for the analysis the last 8 μs of the 10 μs-CG MD simulation with Martini force field ^91–93^ of A_2A_R in two different membranes. (A) Active state of A_2A_R in membrane composed by POPC – 20% cholesterol. (B) Inactive state of A_2A_R in in membrane composed by POPC – 20% cholesterol. (C) Active state of A_2A_R in a plasma mimetic membrane. (D) Inactive state of A_2A_R in a plasma mimetic membrane.

### Binding sites analysis for active A_2A_R in POPC - cholesterol membrane

The PyLipID analysis of cholesterol contact frequencies for active A_2A_R in POPC - 20% cholesterol afforded 10 cholesterol binding sites, BS0′-BS9′ for the active form of A_2A_R (Figure S2). The residence times of cholesterol in each of these sites are shown in Table S2 and Figure S2. We grouped together some of these binding sites that were in proximity by visual inspection, leading to 8 (BS0–BS7) distinct cholesterol binding sites for the active A_2A_R, shown in Figure 2A. We focused our analyses in binding sites that have residues with more than 1 μs cholesterol residence time.

**Figure 2.**
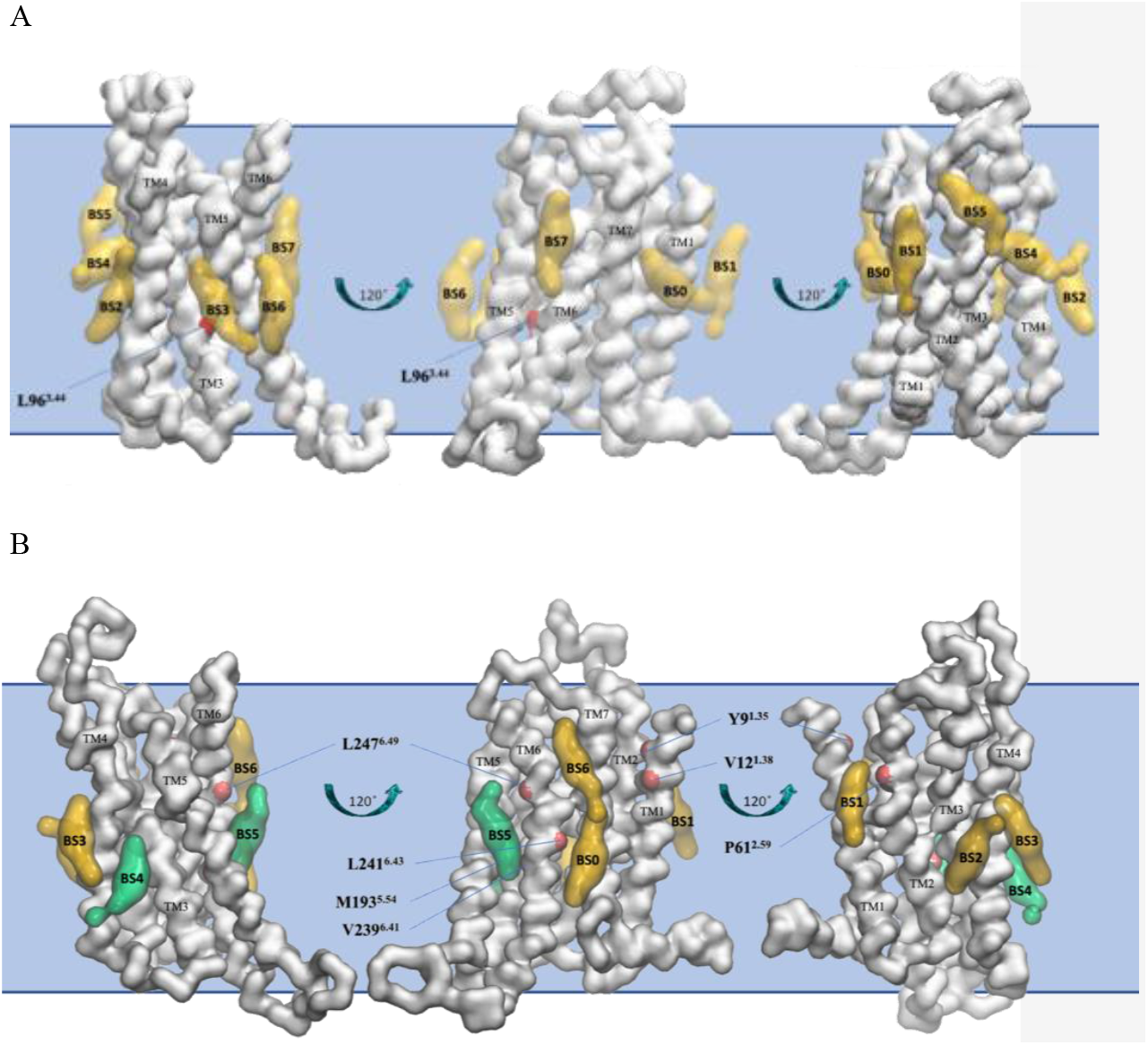
(A) Binding poses of cholesterol in the eight distinct binding sites (BS0-BS7) for the active state of A_2A_R in POPC – 20% cholesterol membrane. (B) Binding poses of cholesterol in the seven distinct binding sites (BS0-BS6) for the inactive state of A_2A_R in POPC – 20% cholesterol membrane. Binding poses were identified after the analysis of the last 8 μs of the 10 μs-CG MD simulations (three repeats) with Martini force field. ^91–93^ The receptor is shown in white surface and representative cholesterol binding poses are shown in green surface (residence time in these binding sites is more than 1 μs) or yellow surface (residence time in these binding sites is less than 1 μs). Residues that belong to the identified binding sites with more than 1 μs cholesterol residence time are shown with a red surface.

For the active state of A_2A_R, we identified ten binding sites namely BS0′-BS9′ (Figure S2 left part) that span around the middle and upper area of the receptor TMD (Figure 2A with residence times shown in Figure S2 left part and Table S2. After visual inspection and grouping together we ended up with eight binding sites (BS0-BS7) shown in Figure 2A. From these distinct eight binding sites we observed only one residue with more than 1 μs cholesterol residence time, namely L96^3.44^ with residence time 1.19 μs in which lies on TM3.

### Binding sites analysis for inactive A_2A_R in POPC - cholesterol membrane

The PyLipID analysis afforded 13 binding sites, BS0′–BS12′ for the inactive state (Figure S2) with residence times of cholesterol in each of these sites shown in Table S2 and Figure S2. After grouping these sites we ended up with 7 distinct cholesterol binding sites, BS0–BS6, for the inactive forms of A_2A_R (Figure 2B). Binding sites of cholesterol with frequent contacts include BS0, BS1, BS5 and BS6. BS0 lies along TM6 in the lower leaf and cholesterol has frequent contacts with L241^6.43^ (1.37μs) and in BS6 which lies in the upper leaf of TM6 with L247^6.49^ (1.53μs). BS1 is located in the middle area of TMD between TM1 and TM2 and has frequent interactions with Y9^1.35^ (1.05 μs), V12^1.38^ (1.13 μs) and P61^2.59^ (1.05μs) while BS5 between TM5 and TM6 and cholesterol interacts with M193^5.54^ (1.22μs) and V239^6.41^ (1.1μs).

### A_2A_R in plasma mimetic membrane

Mammalian plasma membranes are composed of approximately 65% glycerophospholipids, 10% sphingolipids, such as SPH and glycosphingolipids, and 25% of sterols such as cholesterol. ^109^ The extracellular membrane leaflet is enriched in phosphatidylcholine containing POPC, DOPC and sphingolipids. In contrast, the intracellular leaflet is enriched in the phosphatidylethanolamine lipids POPE and DOPE, the phosphatidylserine lipids POPS, DOPS, and the phosphatidylinositol lipid PIP_2_. One consequence of this composition is that the intracellular leaflet of the plasma membrane is anionic in nature. ^109,110^ Thus, to simulate the plasma membrane of the cell, we tested a mixed, asymmetric membrane that contained ten different lipid species (species and abundance are shown in Table S1 and described in Methods Section).

Plotting the residue-wise residence time of cholesterol binding sites in the inactive state of A_2A_R in plasma membrane, we observed that, while most of the cholesterol binding sites are common compared to the POPC – 20% cholesterol membrane, they were calculated with a “noise” reduction and the cholesterol residence time is 2- or 3-fold higher (Figures 1B, D). It is likely that because of the abundance of different lipid species and competition with cholesterol for receptor binding, cholesterol only binds to the most favored binding sites and less preferable positions are occupied by other lipid species. Thus, in plasma membrane (Figure 1D) we observed two regions with cholesterol residence time of ca. 1 μs, (centered at residues ca. 15, 270), two regions with cholesterol residence time, ca. 2-3 μs (centered at residues ca. 60, 190), and one region centered at residue ca. 245 with cholesterol residence time, ca. 3.5-4 μs.

Plotting the residue-wise residence time of cholesterol for the active state of A_2A_R, we observed that the residence time in the region 95-240 (Figure 1C) is low and the relevant sites seen occupied in the POPC – 20% cholesterol simulations are absent. We observed binding sites with highest cholesterol residence time in four regions around residues 10, 70, 248 and 300 with residence time ∼ 2 μs and one region around residue 270 with cholesterol residence time of ∼ 8 μs.

### Binding sites analysis for active A_2A_R in plasma mimetic membrane

In our simulations of A_2A_R in plasma mimetic membrane, we observed slightly more binding sites compared to the POPC – 20% cholesterol membrane, with 11 cholesterol binding sites, BS0′–BS10′ for the active A_2A_R and 13 cholesterol binding sites, BS0′–BS12′ for the inactive state of A_2A_R (Figure S3). The residence times of cholesterol in each of these sites are shown in Table S3 and Figure S3. We grouped together some of these binding sites by visual inspection, leading to 9 (BS0–BS8) for the active state, shown in Figure 3A spanning the whole TMD.

**Figure 3.**
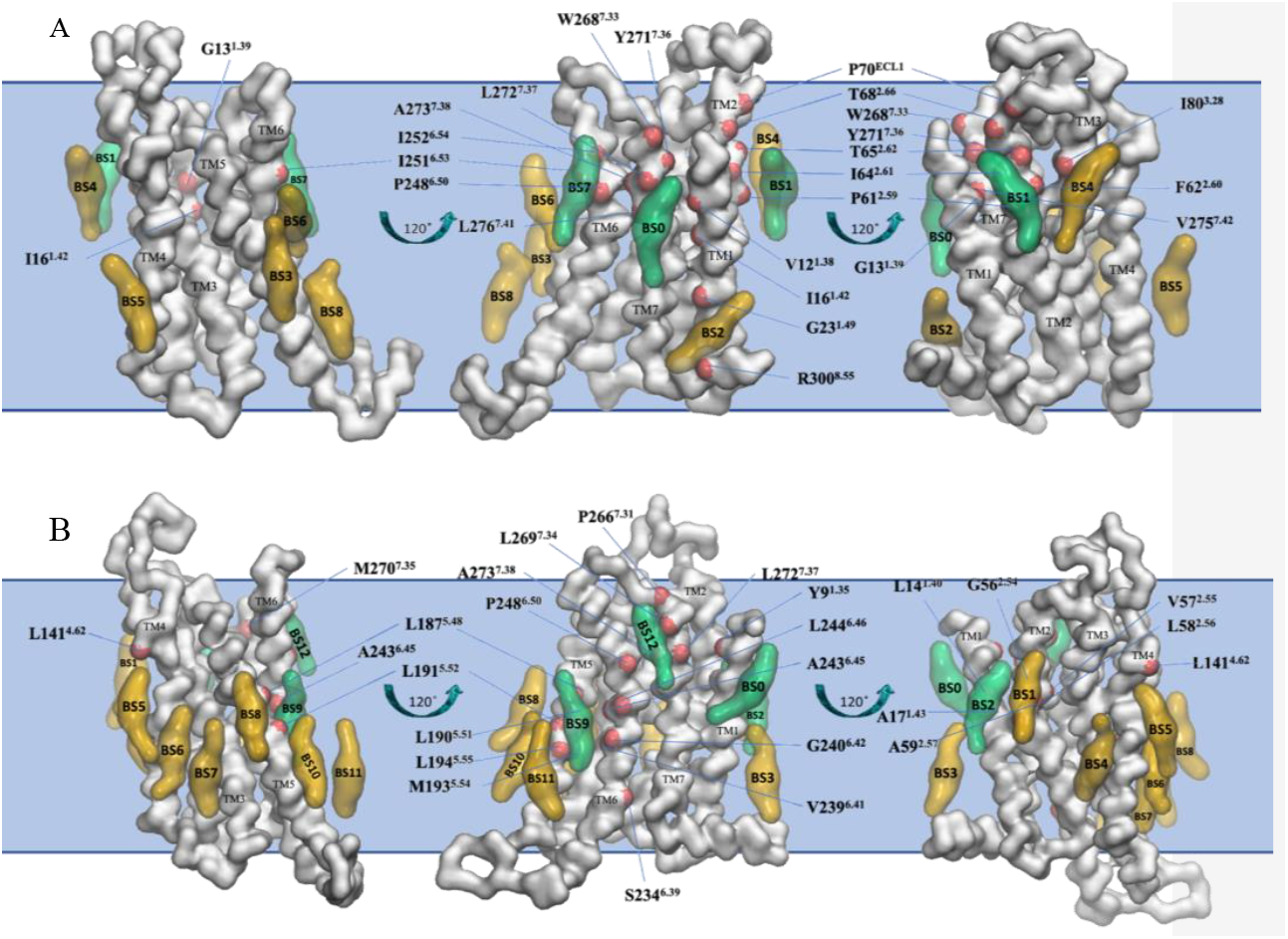
(A) Binding poses of cholesterol in the nine distinct binding sites (BS0-BS8) for the active state of A_2A_R in plasma mimetic membrane. (B) Binding poses of cholesterol in the thirteen distinct binding sites (BS0-BS12) for the inactive state of A_2A_R in plasma mimetic membrane. Binding poses were identified after the analysis of the last 8 μs of the 10 μs-CG MD simulations (three repeats) with Martini force field.^91–93^ The receptor is shown in white surface and representative cholesterol binding poses are shown in green surface (residence time in these binding sites is more than 1 μs) or yellow surface (residence time in these binding sites is less than 1 μs). Residues that belong to the identified binding sites with more than 1 μs cholesterol residence time are shown with a red surface.

The first binding site BS0 is formed in the middle region of TMD between TM1 and TM7 (Figure 5A) and include residues V12^1.38^ (1.86 μs), G13^1.39^ (1.12 μs), I16^1.42^ (1.4 μs), W268^7.33^ (8 μs), Y271^7.36^ (1.62 μs), L272^7.37^ (8 μs), A273^7.38^ (1.33 μs), L276^7.41^ (1.4 μs) and V275^7.42^ (1.50 μs). BS1 lies on TM2 and EL2 and cholesterol has frequent contacts with residues P61^2.59^ (1.62 μs), F62^2.60^ (1.53 μs), I64^2.61^ (1.73 μs), T65^2.62^ (1.75 μs), T68^2.66^ (1.4 μs), P70^ECL1^ (1.89 μs). BS2 is formed between TM8 and TM1 with residues G23^1.49^ (1.12 μs) and R300^8.55^ (2.38 μs). BS7 lies on TM6 and has residues P248^6.50^ (1.03 μs), I251^6.53^ (1.39 μs), I252^6.54^ (2.05 μs) in contact with cholesterol. BS4 lies between TM3 and TM4 and cholesterol has frequent contacts with I80^3.28^ (1.8 μs).

**Figure 4.**
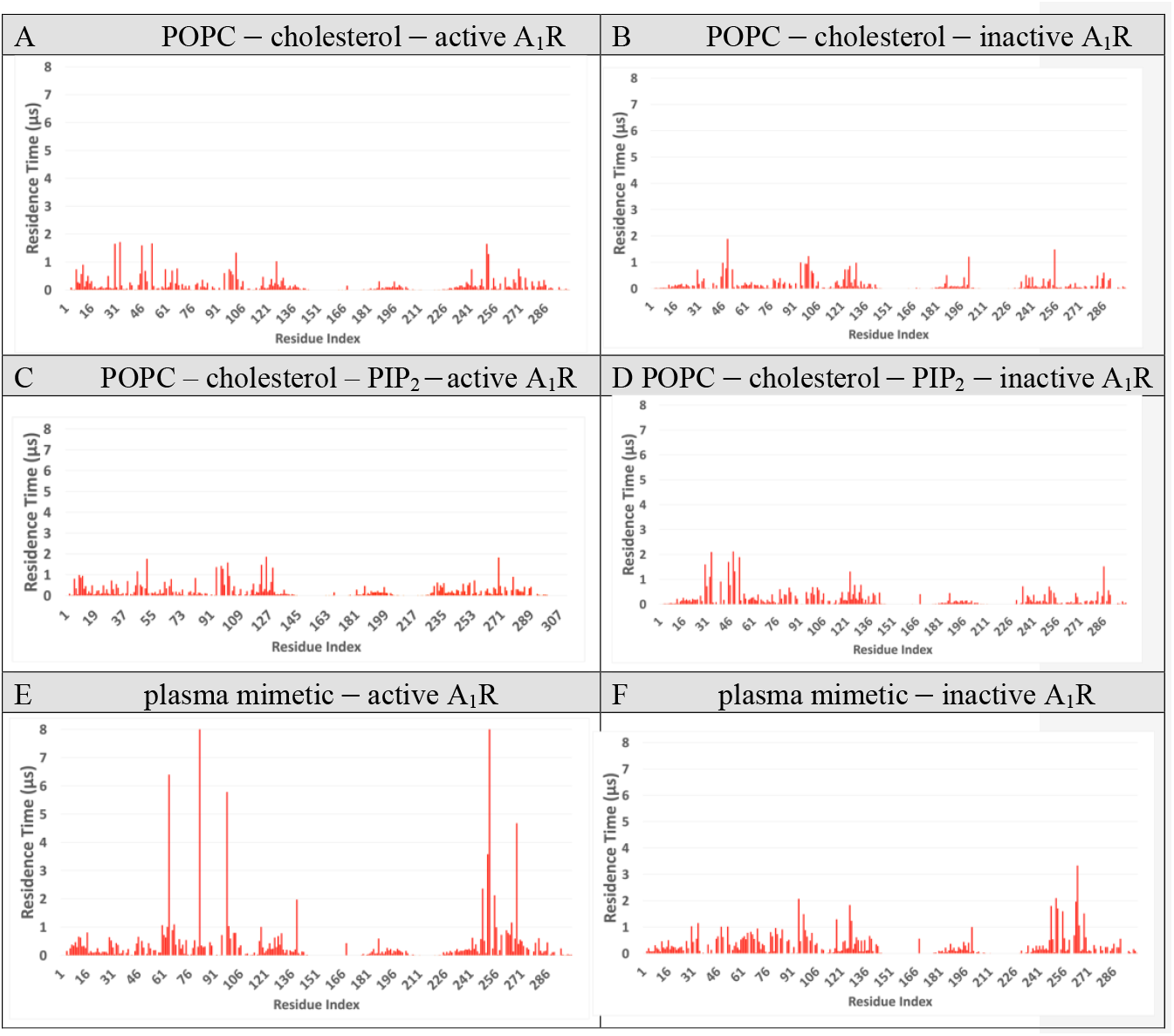
Residence time of cholesterol averaged for three repeats using for the analysis the last 8 μs of the 10 μs-CG MD simulation with Martini force field ^91–93^ of A_1_R in different membranes. (A) Active state of A_1_R in membrane composed by POPC ‒ 20% cholesterol. (B) Inactive state of A_1_R in membrane composed of POPC ‒ 20% cholesterol. (C) Active state of A_1_R in membrane composed by POPC ‒ 25% cholesterol (extracellular) / cholesterol ‒ 25% cholesterol -10% PIP2 (intracellular) (D) Inactive state of A_1_R in POPC ‒ 25% cholesterol (extracellular) / POPC ‒ 25% cholesterol ‒ 10% PIP2 (intracellular). (E). Active state of A_1_R in a plasma mimetic membrane. (F) Inactive state of A_1_R in a plasma mimetic membrane.

**Figure 5.**
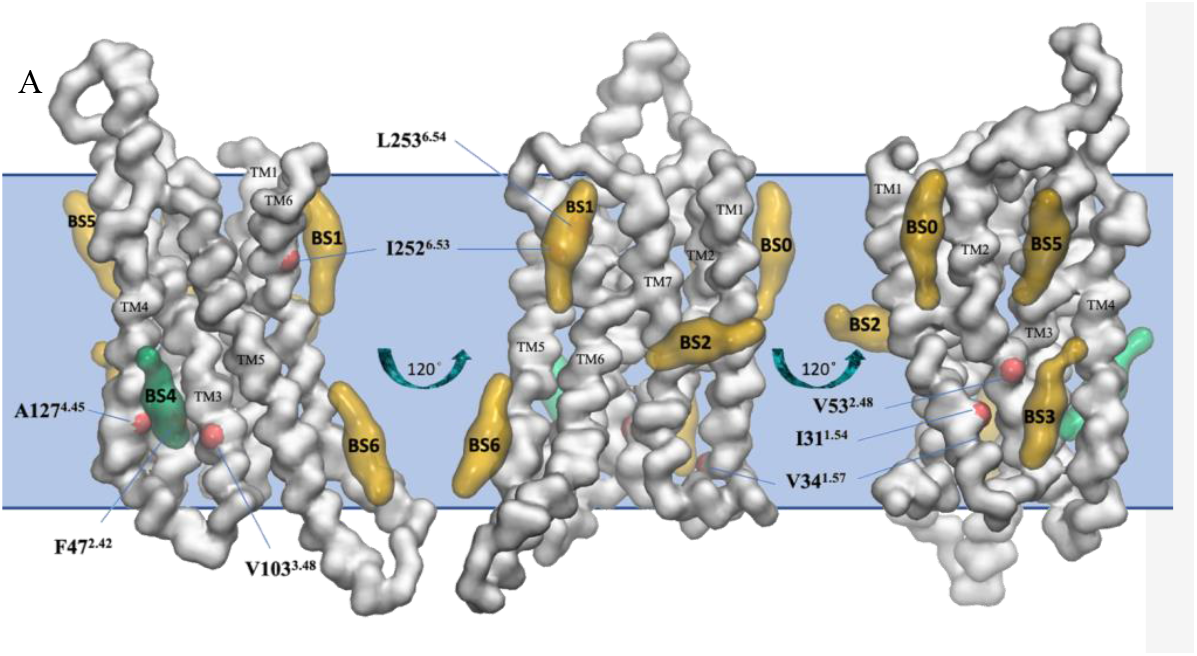
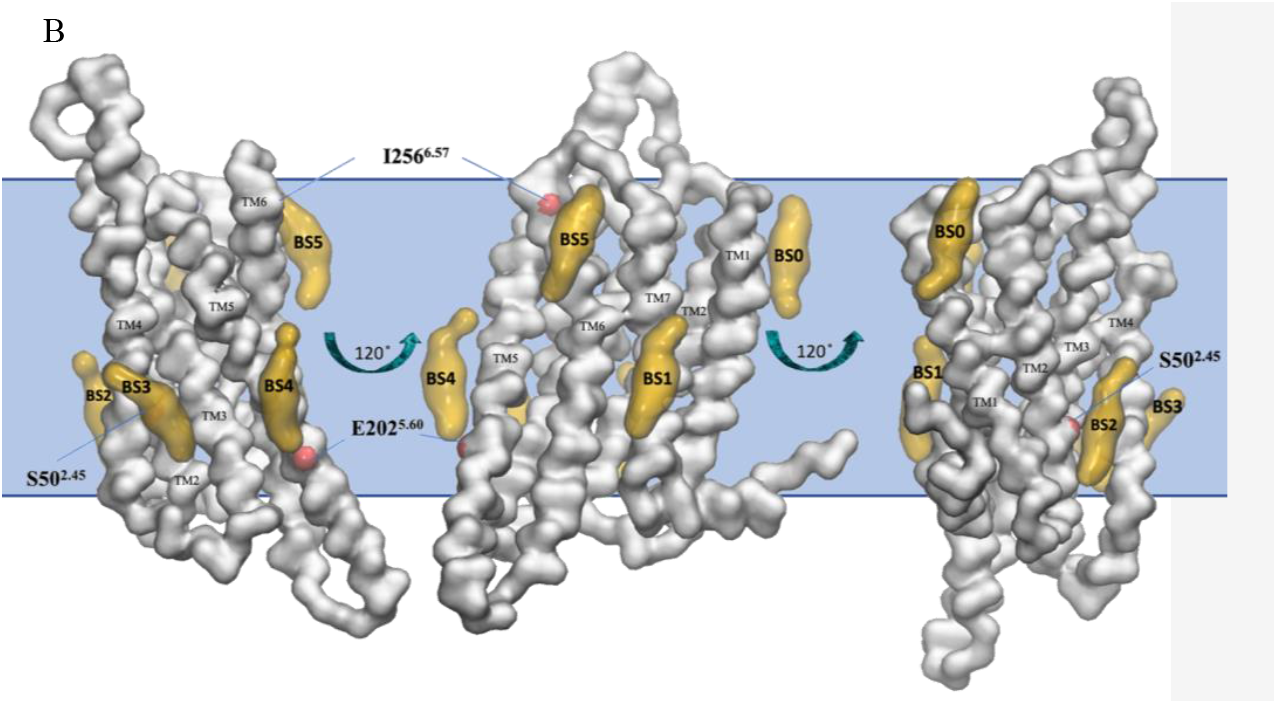
(A) Binding poses of cholesterol in the seven distinct binding sites (BS0-BS6) for the active state of A_1_R in a POPC - 20% cholesterol membrane. The approximate position of the membrane is shown in blue. (B) Binding poses of cholesterol in the six distinct binding sites (BS0-BS5) for the inactive state of A_1_R in POPC - 20% cholesterol membrane. Binding poses were identified after the analysis of the last 8 μs of the 10 μs-CG MD simulations (three repeats) with the Martini force field.^91–93^ The receptor is shown in white surface and representative cholesterol binding poses are shown in green surface (residence time in these binding sites is more than 1 μs) or yellow surface (residence time in these binding sites is less than 1 μs). Residues that belong to the identified binding sites with more than 1 μs cholesterol residence time are shown with a red surface.

### Binding sites analysis for inactive A_2A_R in plasma mimetic membrane

Again, we grouped together some of these binding sites by visual inspection, leading to 13 (BS0–BS12) distinct cholesterol binding sites for the inactive state (Figure 3B), spanning also the whole TMD.

In the upper leaf BS0 lies on TM1 with Y9^1.30^ (1.12 μs), BS2 between TM1 and TM2 with L14^1.40^ (1.2 μs), A17^1.43^ (1.09 μs), V57^1.55^ (1.1 μs), L58^1.56^ (2.23 μs), A59^1.57^ (1.28 μs) and BS1 lies on TM2 with G56^2.54^ (1.39 μs). In the lower TMD region, BS5 lies on TM4 with L141^4.62^ (1.13 μs) and BS9 is located between TM5 and TM6 in the intracellular leaf and includes residues L187^5.48^ (1.81 μs), L190^6.51^ (1.77 μs), L191^5.52^ (2.8 μs), M193^5.54^ (2.46 μs), L194^5.55^ (2.17 μs), S234^6.39^ (1.24 μs), V239^6.41^ (3.24 μs), G240^6.42^ (3.2 μs), A243^6.45^ (3.1 μs), L244^6.46^ (1.73 μs). BS12 is between the extracellular part of TM6 and TM7 with residues P248^6.50^ (4.03 μs), I251^6.53^ (1.39 μs), P266^7.33^ (1.37 μs), L269^7.34^ (1.37 μs), M270^7.35^ (1.4 μs), L272^7.37^ (1.02 μs) and A273^7.38^ (1.37 μs). Both BS9 and BS12 have been observed in crystal structures of A_2A_R in the antagonist-bound, inactive form (e.g. PDB ID 4EIY ^10^ or 5IU4 ^27^) and may be cholesterol positions critical for the stabilization of the inactive form of A_2A_R. An additional important finding is that for the inactive state of A_2A_R in both POPC – 20% cholesterol membrane and plasma mimetic membrane, cholesterol binds in the cleft between TM6 and TM7 in the extracellular leaf (BS5 in Figure 2B, and BS12 in Figure 3B) and in the inner leaf in TM6 (BS4 in Figure 2B and BS9 in Figure 3B) which are binding sites previously identified through computational analysis. ^61^ Since our CG MD simulations for A_2A_R predicted the previously resolved experimentally ^10,27^ and computationally ^61,63^ binding sites of cholesterol to A_2A_R.

### A_1_R in POPC - cholesterol membrane

Having established the utility of our simulations using the A_2A_R, we extended our calculations to the A_1_R. For the active state of A_1_R in a POPC ‒ 20% cholesterol membrane (Figure 4A), we observed five regions with cholesterol residence between 1-2 μs (centered at residues ca. 32, 50, 102, 125 and 252). A rough estimation of the location of these sites is shown in the density maps of cholesterol (Figure S4). For the inactive A_1_R state (Figure 4B), the binding regions of cholesterol are similar but the binding frequency is slightly reduced, as is shown in Figures 4B and S5A-C (right part).

### Binding sites analysis for active A_1_R in a POPC ‒ cholesterol membrane

Binding site analysis using PyLipID revealed 10 cholesterol binding sites for the active A_1_R, BS0′-BS9′ (Figure S5 left part) with residence times of cholesterol in each of these sites shown in Table S4 and Figure S5 left part. As before, we grouped these binding sites by visual inspection, which resulted in 7 distinct binding sites, BS0-BS6 (Figure 5A) and then identified residues belonging to these seven binding sites and having more than 1 μs cholesterol residence time.

The first binding site BS0 is formed in the upper part between TM1 and TM2 (Figure 5A). The second binding site BS1 is formed between TM6 and TM7 where cholesterol has frequent contacts with I252^6.53^ and L253^6.54^ with residence time for these residues 1.6 μs and 1.3 μs, respectively (Table 1). BS2 is formed between TM7 and TM1 and, since it is perpendicular to the membrane normal, can represent a transition spot for flip-flop movement of cholesterol molecules. BS3 forms a cavity for cholesterol in the intracellular leaf of the membrane between TM1, TM2 and TM4; in this binding site cholesterol has frequent contacts with I31^1.54^, V34^1.57^ and V53^2.48^ with residence time 1.6 μs, 1.7 μs and 1.3 μs, respectively (Table 1). BS4 is spanning a cavity for cholesterol formed between TM3 and TM4 and in this spot cholesterol additionally interact with TM2. Cholesterol has frequent contacts in BS4 with A127^4.45^(1.6 μs), F47^2.42^ (1.7 μs) and V103^3.48^ (1.33 μs) (Table 1). It is worth noting that V103^3.48^ is part of a CRAC motif included by the sequence V103^3.48^-Y106^3.51^-R108^IL2^/K110^IL2^. BS5 is formed by the extracellular part of TM2 and BS6 formed by TM5 has the lowest cholesterol residence times.

### Binding sites analysis for inactive A_1_R in a POPC ‒ cholesterol membrane

For the inactive A_1_R the 9 cholesterol binding sites, BS0′-BS8′ (Figure S5 right part), were identified with residence times for each of these sites shown in Table S5 and Figures S5A-C (right part). We grouped some of them and end up with 6 distinct binding sites, BS0-BS5 (Figure 5B). BS0 lies in the area between TM1 and TM2 and BS1 is formed in the cavity between TM1 and TM7. BS2 is located between TM2 and TM4 and when cholesterol is interacting with BS2 it also makes contacts with residues at TM3. BS3 lies between TM3 and TM4 while BS4, BS5 in TM5. From these distinct six binding sites in Figure 5B we showed only three residues with more than 1 μs cholesterol residence time, namely S50^2.45^ (1.88 μs) in BS3, E202^5.60^ (1.21 μs) in BS4 and I256^6.57^ (1.48 μs) in BS5. Of note, apart from the overall decrease of cholesterol residence time bound to the inactive state of A_1_R compared to the active, we also observed a significant decrease in cholesterol’s residence time close to residue V103^3.48^ which is part of the CRAC motif calculated in active A_1_R.

### A_1_R in a POPC ‒ cholesterol ‒ PIP_2_ membrane

We then simulated the receptors in membranes containing PIP_2_ in the intracellular leaf (Tables S6, S7). Comparisons between the density maps suggested that several binding sites are common between active and inactive A_1_R conformations in POPC ‒ cholesterol and POPC ‒ cholesterol ‒ PIP_2_ membranes (Figure S6). The binding interactions pattern of cholesterol to A_1_R in this membrane are similar to the POPC ‒ 20% cholesterol system as compared from Figures 4A, C and 4B, D or Figure 5 and Figure S8 (see also Figures S5, S7 and Tables 1, S4-S8). This suggests that PIP_2_ does not appear to alter the nature of cholesterol interaction with A_1_R.

### A_1_R in plasma mimetic membrane

The CG MD simulations-derived density maps of cholesterol binding to active A_1_R conformation in plasma mimetic membrane consisting of 10 different lipid species (Methods Sections) do not differ significantly compared to A_1_R active state in POPC ‒ 20% cholesterol membrane (Figures S4, S9). The frequency analysis in Figures 4C,E and 4D,F showed that while most of the binding sites are common, in the plasma membrane we observed the “noise” reduction effect observed also in the case of active A_2A_R (Figures 2A,C). The preferred residues that cholesterol binds from the CG MD simulation are clearly identified (centered at residues ca. 64, 82, 99, 253 with cholesterol residence time ca. 6-7.5 μs and at residues ca. 268 with cholesterol residence time ca. 5 μs) (Figure 4E). For the inactive A_1_R in the plasma membrane (Figure 4F) we observed many amino acid regions to which cholesterol binds. One region centered at residue ca. 265 with cholesterol residence time of ca. 3 μs, three regions (centered at residues ca. 100, 128 and 255) with cholesterol residence time ca. 2 μs, and four regions with cholesterol residence time ca. 1 μs (centered at residue ca. 33, 50, 70, 83). The “noise” reduction was also observed for A_1_R and A_2A_R (see Figures 2B,D) in the inactive state (Figures 4B,F) but was less significant. It seems that the presence of plasma mimetic membrane and active conformation of the AR with more conformational transitions compared to the inactive resulted in the highest affinity cholesterol binding sites.

### Binding sites analysis for active A_1_R in plasma mimetic membrane

Binding site analysis of active A_1_R using PyLipID and after grouping together the 11 binding sites that lie in proximity (Figure S10 left part, with residence time of cholesterol in each of these sites shown in Table S8) we ended up with six distinct binding sites BS0-BS5 shown in Figure 6. BS0 is formed between TM2 and TM3 in the extracellular leaf including five residues (Figure 6A) with cholesterol residence time more than 1 μs, in residues L61^2.56^ (1.06 μs), P64^2.59^ (1 μs), L65^2.60^ (6.39 μs), L68^2.63^ (1.09 μs), V83^3.28^ (8 μs), with V83^3.28^ standing out as retaining the cholesterol contacts during the whole trajectory. BS1 is formed between TM1 and TM7 but interacts also with TM6. In BS1 cholesterol has almost a perpendicular orientation to TM helices. Low residence times of cholesterol in BS1 (Figure S11, left part) suggested that this site between TM1 and TM7 is a metastable binding site which was also observed in the active state of A_1_R in POPC ‒ 20% cholesterol membrane. BS2 lies in TM3 in the intracellular leaf and at this extended binding site L99^3.44^ stands out with cholesterol residence time 5.78 μs with next longer in cholesterol residence time residues A100^3.45^ (1 μs) and V119^IL2^ (1 μs). BS3 in contact with BS2 includes residues in TM1, TM2 and TM4 with L140^4.58^ having the longest residence time of 1.97 μs. BS4 occupies the upper area of A1R - close to the extracellular domain-between TM6-TM7. BS4 lies between TM6 and TM7 in the extracellular region and is formed by the residues C263^EL3^ (0.88 μs), P249^6.50^ (2.36 μs), I252^6.53^ (3.58 μs), L253^6.54^ (8 μs), I256^6.57^ (2.12 μs), P266^7.31^ (1.16 μs) and L269^7.34^ (4.67 μs). Finally, BS5 lies in TM5-TM6-TM7 and has TM5-TM6 part of the binding site exposed to the bulk membrane hydrocarbon lipids.

**Figure 6.**
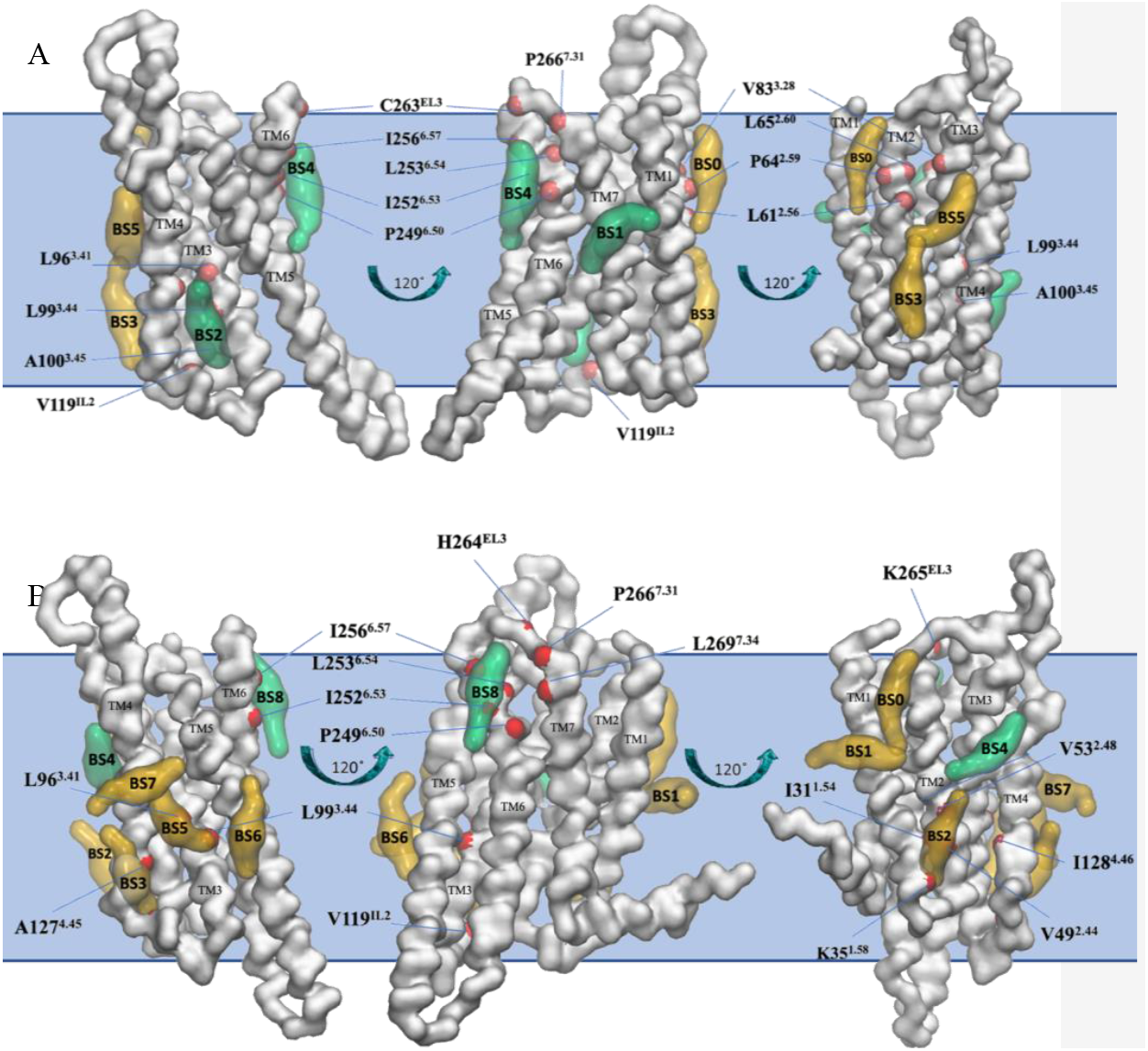
(A) Binding poses of cholesterol in the six distinct binding sites (BS0-BS5) of the active state of A_1_R in plasma mimetic membrane. (B) Binding poses of cholesterol in the nine binding sites (BS0-BS8) for the inactive state of A_1_R in plasma mimetic membrane. Binding poses were identified after the analysis of the last 8 μs of the 10 μs-CG MD simulations (three repeats) with Martini force field.^91–93^ The receptor is shown in white surface and representative cholesterol binding poses are shown in green surface (residence time in these binding sites is more than 1 μs) or yellow surface (residence time in these binding sites is less than 1 μs). Residues that belong to the identified binding sites with more than 1 μs cholesterol residence time are shown with a red surface.

### Binding sites analysis for inactive A1R in plasma mimetic membrane

The nine distinct binding sites (BS0-BS8) are shown in Figure 6B (see also Figure S10 right part, with residence time of cholesterol in each of these sites shown in Table S9) of cholesterol for the inactive state of A_1_R in a plasma mimetic membrane. BS0 and BS1 are located on the upper area of TM1-TM2. In BS2, that lies in a cavity formed by the lower part of TM1-TM2, cholesterol has contacts with residues I31^1.54^ (1.03 μs), K35^1.58^ (1.15 μs), V49^2.44^ (1.01 μs), V53^2.48^ (1.02 μs) and V119^IL2^ (1.29 μs). BS3 with important residue A127^4.45^ (1.83 μs) is located at the lower area of TM4. BS5-BS7 are positioned in the middle-lower area of TM4-TM3 and because of their orientation they may be involved in cholesterol flip-flop movement. Important residues for cholesterol binding are L96^3.41^ (2.07 μs), A127^4.45^ (1.83 μs) and I128^4.46^ (1.24 μs) in BS5. Cholesterol interacts also with residues in BS8 positioned between TM5 and TM7 which are P249^6.50^ (1.80 μs), I252^6.53^ (2.09 μs), L253^6.54^ (1.70 μs), I256^6.57^(1.59 μs), H264^EL3^ (1.96 μs), K265^EL3^ (3.33 μs), P266^7.31^(1.06 μs), L269^7.34^ (1.52 μs) (Figure 6B). Common and uncommon residues are depicted also in Figure S12.

### Potential of Mean Force calculations

To characterize further the A_1_R ‒ cholesterol binding sites in the plasma mimetic membrane we performed US/PMF calculations for characteristic binding sites. We calculated the BS8 (or BS8′, Figure S11) and BS4 (or BS4′, Figure S11) for the inactive A_1_R (Figures 7A,C) and BS4 (BS9′, Figure S11), BS1 (BS2′, Figure S11) for the active A_1_R (Figures 7B,D). The results in Figure 7 suggested that for the active state A_1_R in BS4 cholesterol is stabilized with a binding free energy minimum, of ∼ ‒ 6.9 kJ/mol (1.65 kcal/mol) while for the inactive state A_1_R in BS8 cholesterol is stabilized with a minimum, of ∼ ‒ 5 kJ/mol (1.20 kcal/mol). In contrast in the low residence binding sites BS1 and BS4 for active and inactive state respectively, cholesterol lies in unstable minima that are quite rapidly exchange with bulk concentration of cholesterol in membrane with thermal fluctuations.

**Figure 7.**
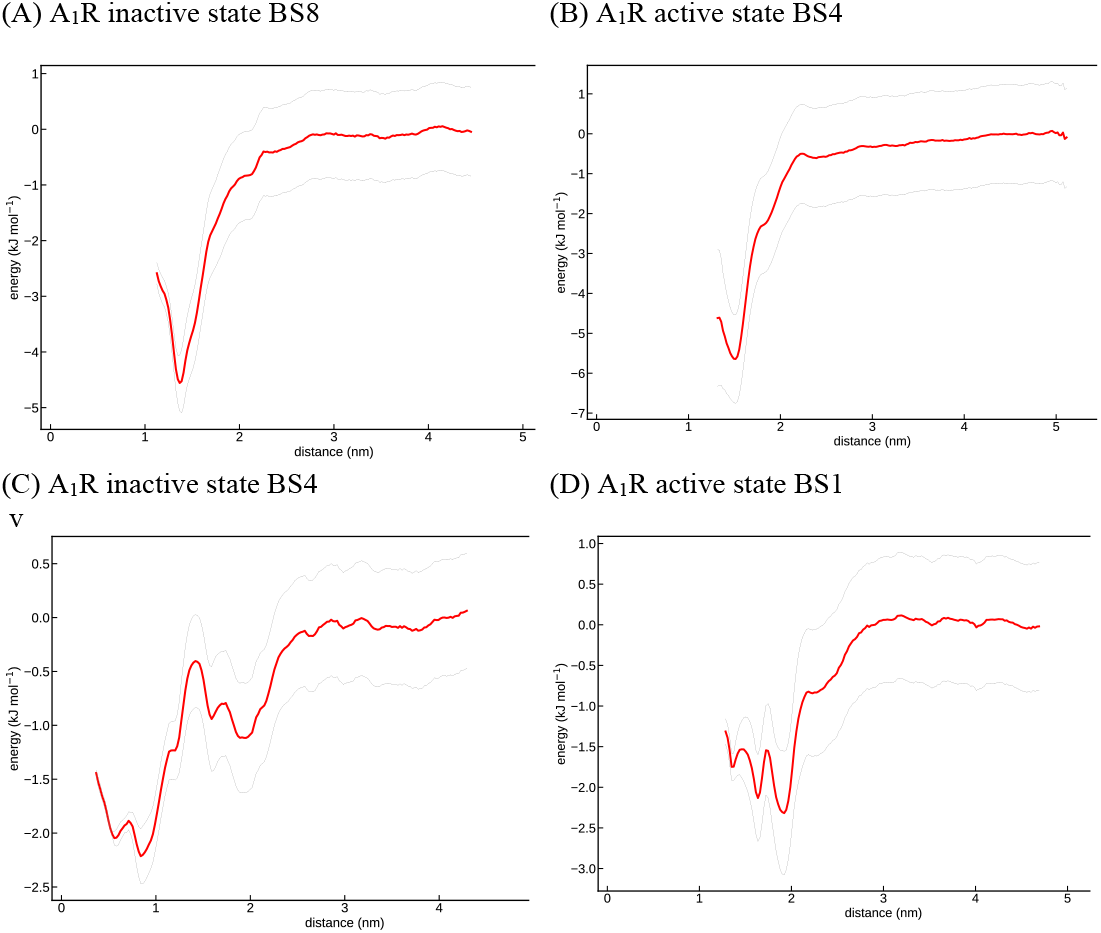
US/PMF plots with Martini force field ^91–93^ for selected binding sites of A_1_R active and inactive states in plasma mimetic membrane. (A) A_1_R inactive state BS8 (or BS8′ in Figure S11) (B) A_1_R active state BS4 (BS9′in Figure S11) (C) A_1_R inactive state BS4 (or BS4 in Figure S11) (D) A_1_R active state BS1 (BS2′ in Figure S11).

## Discussion

Plasma membranes concentrate 80–90% of the total cell cholesterol content, that represents 20– 25% of total lipids in the plasma membrane and 34% of the total lipid content in mammalian plasma membranes ^111,112^ and its high content affects GPCR structure and function. ^113,114^ Membrane cholesterol has been reported to have a modulatory role in the function of a number of GPCRs, in particular on signaling and recycling on ligand binding. ^24^ As some GPCRs are enriched in cholesterol-rich domains, these signaling complexes may also promote interactions between GPCRs. Homo- or hetero-oligomerization, that also play a role in the signaling of some GPCRs, may be influenced by membrane cholesterol content and cholesterol mediated binding sites. ^115^

The 25% of the cholesterol total body content is found in the central nervous system and cholesterol has been shown to affect serotonin 1A (5-HT1A) receptors. ^116,116^ It has been demonstrated the inhibitory function of cholesterol against α- and β-adrenergic receptors which is corroborated by the enhanced signalling of α- and β-adrenergic receptors in cardiomyocytes depleted of cellular cholesterol. ^117^ Being highly abundant in the cell membrane cholesterol may be a prime regulator of adrenergic receptors, keeping their basal activity low by stabilizing their intermediate active conformation. It is also very likely that the activity of many further GPCRs is modulated by cholesterol. For example, cholesterol was recently shown to act as a positive allosteric modulator of the cannabinoid receptor CB2, since depletion of cholesterol from the plasma membrane decreases its activity, both in the presence of agonists and in the apo state. ^118^ Furthermore, cholesterol experimentally is a known stabilizer of β_2_ adrenergic receptor (β_2_AR) ^39^ and A_2A_R ^82^ and has been suggested as allosteric modulator of cholecystokinin 1 (CCK1) receptor, ^119,47^ cannabinoid CB2 receptor, ^120^ μ-opioid Mu 1 receptor (OPRM1), ^121^ oxytocin receptor (OXTR). ^122^ Also, it has been observed that the signaling of A_2A_R, coupled to Gα_s_ is reduced with cholesterol depletion; ^82^ this indicates that cholesterol plays an important role in G_αs_ mediated cAMP accumulation, independently of ligand binding stimulation.

Cholesterol also impacts ligand binding affinity for several GPCRs, but the direction of modulation (increasing or decreasing) is not the same since positive, negative and no modulation have been reported. Thus, the binding affinity of a selective antagonist was enhanced by cholesterol depletion in the case of A_2A_R selective antagonist ^59^ and was reduced in presence of cholesterol for chemokine receptor type 5 (CCR5). ^123^ In contrast, the binding affinity of an agonist was increased in a dose-dependent manner with cholesterol concentration for the chemokine receptor CCR3 ^124^ or oxytocin receptor (OTR) ^124 125^ and was reduced by cholesterol depletion in the case of 5-HT1A receptors. ^116^

The first structural evidence for site-specific cholesterol binding in GPCRs was provided by early X-ray structures of inactive states of β_2_AR ^25^ and A_2A_R, ^10^ in which cholesterol binds in distinct TM positions forming cavities. Much progress has been made since, and now there are hundreds of experimentally resolved GPCR structures available. ^26^ Empty cavities may fulfill key functions in GPCRs such as high-affinity binding of hydrophobic ligand moieties ^126^ or directing functional motions. ^127^ It has recently shown for β_1_AR that cholesterol fills specific voids that must be compressed for β_1_AR activation acting as allosteric antagonists. ^51^

Filling spaces between the transmembrane helices modulate the dynamics and activity of many GPCRs, as demonstrated by the development of a number of positive and negative allosteric modulators, ^128^–^134^ which occupy such crevices. In contrast to the orthosteric binding sites, the surfaces of allosteric binding sites are often lined by amino acids that are distinctive between GPCR subtypes. This property gives them a high potential for the development of therapeutic drugs targeting specific GPCR subtypes. Several allosteric modulators of other GPCRs bind to the same site of a cholesterol as in β_1_AR. Although their binding site is identical, the functional output may vary. For example, such allosteric modulators increase the activity of GPR40 ^129^ and DRD1, ^131^ but down-modulate β_2_AR ^128^ and C5aR1 ^132^ or CB1. ^135^ Blocking TM5 sliding appears as the hallmark of their mechanism of action with the position of the blocked TM5 relative to the active conformation defining the direction of modulation.

Unlike A_2A_R where computational work ^58–62,63^ experimental structural ^10,27–35,36^ and functional data ^5982^ are available for cholesterol binding such data are not available for A_1_R with experimental structures available after 2017. ^14^–^17^ In A_2A_R has been suggested that the two cholesterol molecules sandwich the phenol ring of Phe255^6.57^ in close proximity of the binding pocket and stabilize the conformation of the extracellular part of TM6. ^10^

Here, we employed CG MD simulations to predict specific binding sites of cholesterol with A_1_R in the antagonist-bound inactive state and agonist-bound active state, embedded in model membranes. We used as model membrane POPC – cholesterol, POPC – cholesterol – PIP_2_ and a plasma mimetic membrane. The latter seems to filter much better cholesterol binding interactions. In Figure 8 are shown the low and high residence receptor for the two states of A_1_R.

**Figure 8.**
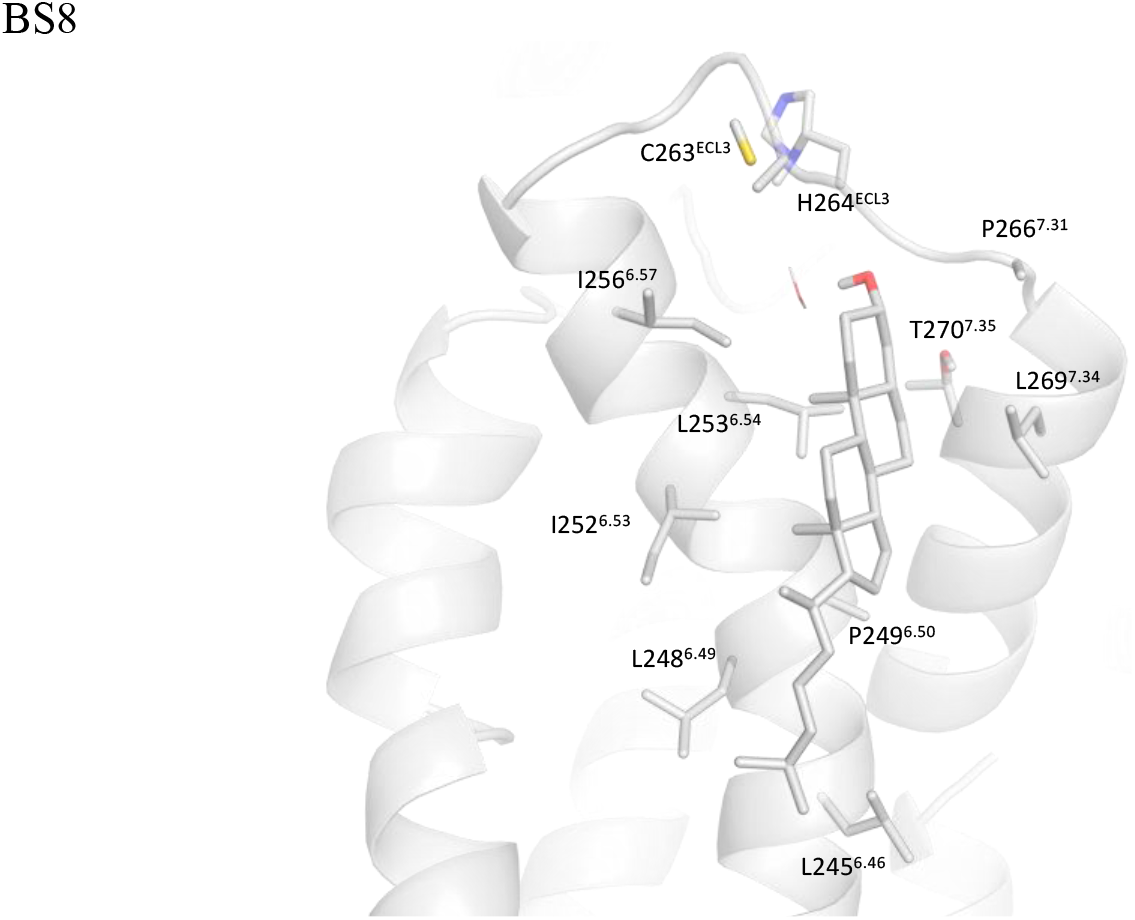
Snapshot showing disposition of cholesterol in BS8 (see Figure 6) between TM6 and TM7 in the extracellular region of the receptor with inactive A_1_R after mack-mapping to an AA model the last snapshot from 10 μs-CG MD simulations with Martini force field. ^91–93^ cholesterol can form a hydrogen bond with to T270^7.35^ and stabilized with hydrophobic interactions with I256^6.57^, L253^6.54^, L269^7.34^ and P249^6.50^, L248^6.49^, L245^6.46^. The receptor is shown in white surface; cholesterol and side chains are shown in sticks.

The analysis of binding sites to the inactive A_1_R in the plasma mimetic membrane cholesterol showed that cholesterol binds in BS8 in TM6, TM7 in the extracellular region of the receptor including residues P249^6.50^ (1.80 μs), I252^6.53^ (2.09 μs), L253^6.54^ (1.70 μs), I256^6.57^(1.59 μs), P266^7.31^(1.06 μs), L269^7.34^ (1.52 μs). This binding site has been observed to accommodate cholesterol molecules as observed in crystal structures of A_2A_R in the inactive state ^10,27,29,30,33,36,80^ and as predicted computationally in the current and previous work. ^61^ This may be a position critical for the stabilization of the inactive state of A_2A_R and A_1_R. In Figure 8 is shown an all-atom model of cholesterol binding pose resulted from back-mapping of the CG MD simulation last snapshot. The hydroxyl group of cholesterol can be hydrogen bonded to T270^7.35^ while the ring skeleton is embraced by I256^6.57^, L253^6.54^, L269^7.34^ and the alkyl chain by P249^6.50^, L248^6.49^, L245^6.46^.

In Figure 9 the atomistic model of BS0 lying on TM2/TM3 in the extracellular leaf of active A_1_R is shown after back-mapping the last snapshot of CG MD simulations. Cholesterol is stabilized by van der Waals interactions with residues L68^2.63^, P64^2.59^, I63^2.58^, A60^2.55^, V83^3.28^ while V83^3.28^ has the highest cholesterol residence time (8 μs) being in contact with cholesterol during the whole trajectory.

**Figure 9.**
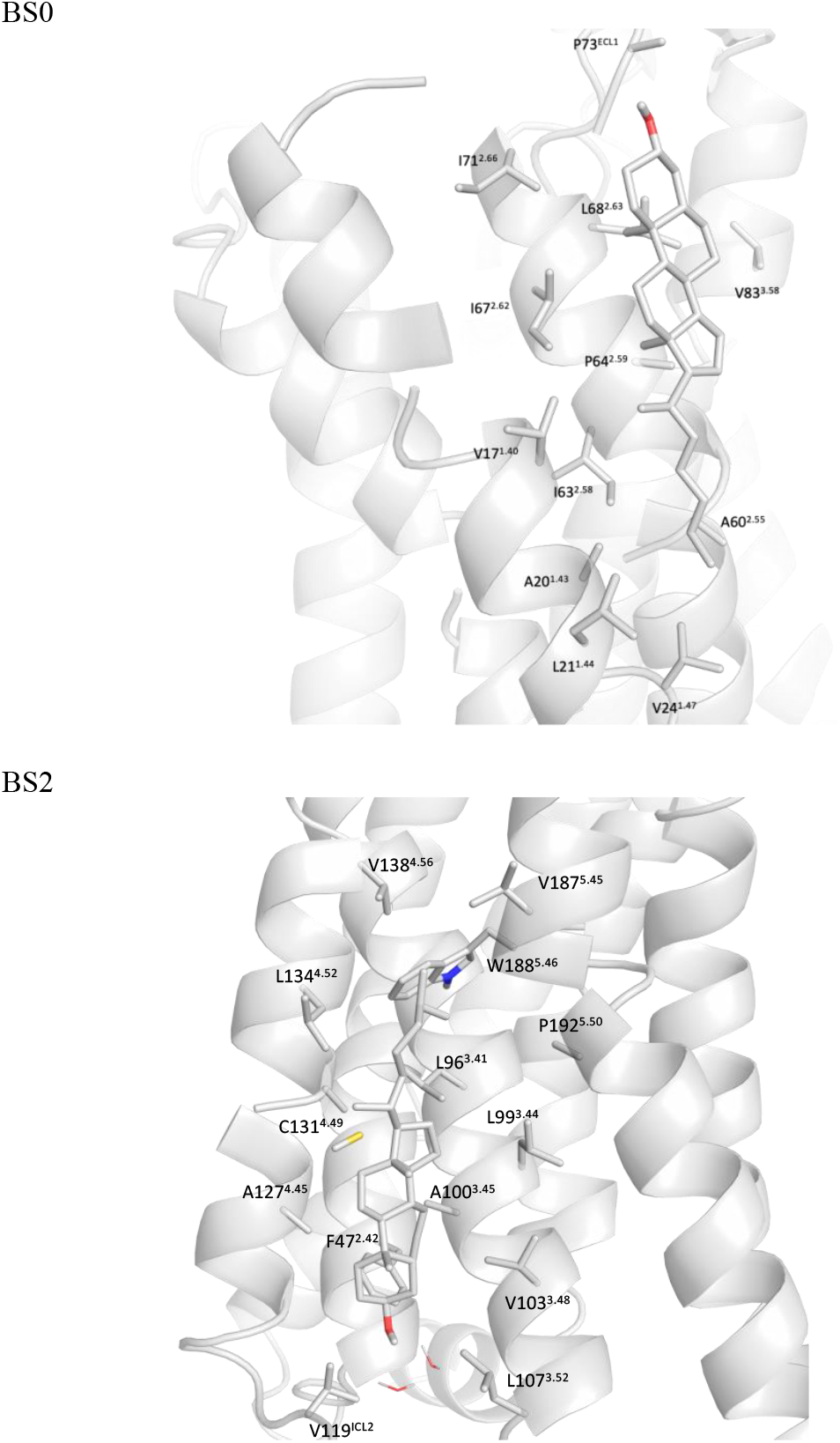
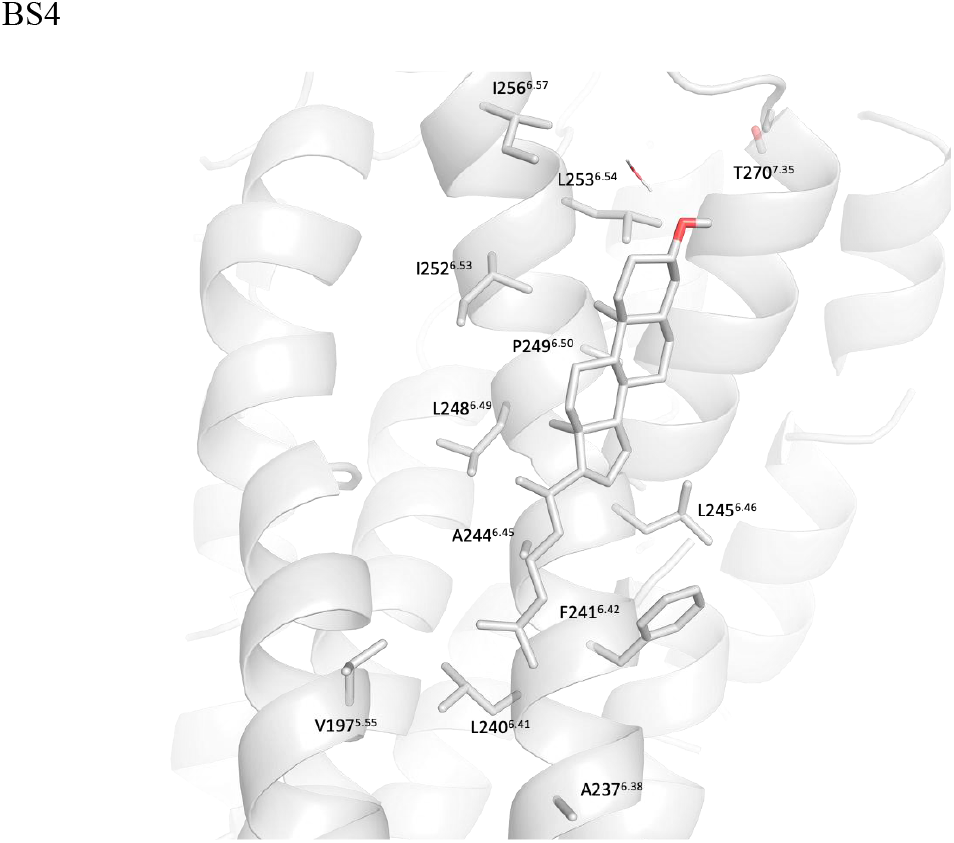
Snapshot showing disposition of cholesterol in the active A_1_R state after back-mapping to an AA model the last snapshot from 10 μs-CG MD simulations with Martini force field. ^91–93^ It is shown BS0 which lies between TM2/TM3 in the extracellular leaf and is stabilized by van der Waals interactions including residues L68^2.63^, P64^2.59^, I63^2.58^, A60^2.55^ with V83^3.28^ being in contact during the whole trajectory(8 μs); in BS2 which lies in TM3 in the intracellular region and is stabilized by van der Waals contacts with L99^3.44^ (5.78 μs), A100^3.45^and V119^IL2^; in BS4 which lies between TM6, TM7 in the extracellular region and is stabilized by van der Waals interactions with C263^EL3^ (0.88 μs), P249^6.50^ (2.36 μs), I252^6.53^ (3.58 μs), L253^6.54^ (8 μs), I256^6.57^ (2.12 μs), P266^7.31^ (1.16 μs) and L269^7.34^ (4.67 μs). The receptor is shown in white surface; cholesterol and side chains are shown in sticks.

In BS2 of the active A_1_R, cholesterol lies along TM3 in the intracellular having van der Waals interactions with V119^IL2^ (1 μs), A100^3.45^ (1 μs) and L99^3.44^ (5.78 μs). BS4 lies between TM6 and TM7 in the extracellular leaf and is stabilized by van der Waals interactions with C263^EL3^ (0.88 μs), P249^6.50^ (2.36 μs), I252^6.53^ (3.58 μs), L253^6.54^ (8 μs), I256^6.57^ (2.12 μs), P266^7.31^ (1.16 μs) and L269^7.34^ (4.67 μs). Although several of the residues in BS8 of the inactive A_1_R are common with BS4 in the active A_1_R, the residence time of cholesterol in the active state of A_1_R is considerably longer including residues I252^6.53^ (3.58 μs), L253^6.54^ (8 μs), L269^7.34^ (4.67 μs) in agreement with the observed free energy minima obtained from the US/PMF plots shown in Figure 7.

Cholesterol binding can be conditioned just on hydrophobic residue environment that stabilizes cholesterol and a geometric compatibility between cholesterol and the protein interface, which accommodates its ring structure (Figure 9). While shielding of the cholesterol hydroxyl group from the hydrophobic environment may be important (Figure 8). ^136^

In A_2A_R the highest residence time is BS0 formed in the middle region of TMD between TM1 and TM7 (Figure 5A) and including residues V12^1.38^ (1.86 μs), G13^1.39^ (1.12 μs), I16^1.42^ (1.4 μs), W268^7.33^ (8 μs), Y271^7.36^ (1.62 μs), L272^7.37^ (8 μs), A273^7.38^ (1.33 μs), L276^7.41^ (1.4 μs) and V275^7.42^ (1.50 μs).

We identified the previously detected experimentally or computationally binding sites of inactive A_2A_R. Thus, for the inactive A_2A_R we detected that cholesterol binds stably in the cleft between TM6 and TM7 in the extracellular leaf and in the inner leaf in TM5/TM6 which are binding sites previously identified with AA and CG MD simulations ^61^ and have also been previously observed in crystal structures of inactive A_2A_R. While it has been shown that cholesterol is a stabilizer of inactive A_2A_R ^82^ conformation it has been shown that depletion of cholesterol attenuates signaling of A_2A_R, coupled to G_αs_. ^82^

We calculated that in the active A_1_R cholesterol binds to a similar extracellular area between TM6 and TM7 as in the inactive A_1_R and inactive A_2A_R ^10,27^ but with much longer residence time. The cholesterol binding in this region can stabilize the active A_1_R conformation but the binding area is partially overlapped with this in the inactive A_1_R state, with cholesterol molecules antagonizing the inactive with respect to the active state.

We also observed for the active A_1_R two cholesterol binding sites in the extracellular membrane leaf in a cavity between TM2/TM3 and along TM3 in the intracellular leaf. For the active A_2A_R we predicted a high residence time cholesterol position between TM1 and TM7 in the middle region of TMD which is different from the binding sites calculated in active or inactive A_1_R and for the inactive A_2A_R.

It remains generally unclear how A_1_R-cholesterol interactions modulate ligand binding, functional state, and function, such as receptor activation and downstream signaling, in a physiologically relevant context and new experimental structures and functional experiments may help addressing these questions.

The differences in cholesterol binding sites between active and inactive states of A_1_R and for A_2A_R can be important for functional activity and orthosteric agonist or antagonist affinity (see also Table of Contents Graphic). A suggestion on the mechanism of how cholesterol modulates functional activity of β_2_AR has been suggested recently. ^51^ Cholesterol can fill spaces between the transmembrane helices which are required to be suppressed for activation. ^51^ Cholesterol displacement can suppress these voids and modulate allosterically the conformational dynamics by shifting the equilibrium to the active state. It has also been suggested that cholesterol binding to certain cavities can stabilize the active or inactive state and can shift the equilibrium accordingly. For example, the binding affinity of an agonist has been shown to increase in a dose-dependent manner with cholesterol concentration for the chemokine receptor CCR3 ^124^ or oxytocin receptor (OTR) ^124,125^ or for an antagonist against angiotensin II receptor type 1, ^137^ and has been shown to reduce by cholesterol depletion in the case of 5-HT1A receptors. ^116^ Thus, these stable, long residence time cholesterol binding sites can be used for the design of allosteric modulators which can bind through lipid pathways as has been described for CB1 receptor ^138,135^ and can be accomplished for other GPCRs, e.g. ARs.

## Supporting information

GitHub - etankol/CG_Adenosine_Receptors

## Abbreviations

AR: adenosine receptor
β_2_AR: β_2_ adrenergic receptor
cAMP: cyclic adenosine monophosphate
CCK1: cholecystokinin 1
CCM: Cholesterol Consensus Motif
CNC: Cholesterol Network Cluster
CNR1: cannabinoid CB1 receptor
CG: coarse grained
CRAC: Cholesterol Recognition Amino Acid Consensus
cryo-EM: cryogenic electron microscopy
DOPC: 1,2-dioleoyl-*sn*-glycero-3-phosphocholine
DOPE: 1,2-dioleoyl-*sn*-glycero-3-phosphoethanolamine
DOPS: 1,2-dioleoyl-sn-glycero-3-phosphoserine
GM3: monosialodihexosylganglioside
GPCR: G protein-coupled receptor
MD: molecular dynamics
NECA: 5’-N-ethylcarboxamidoadenosine
OPRM1: μ-opioid Mu 1 receptor
OXTR: oxytocin receptor
OPM: orientation of proteins in membranes
PDB: Protein Data Bank
PIP_2_: phosphatidylinositol-bisphosphate
PME: Particle Mesh Ewald
POPC: 1-palmitoyl-2-oleoyl-sn-glycero-3-phosphocholine
POPE: 1-palmitoyl-2-oleoyl-*sn*-glycero-3-phosphoethanolamine
POPS: 1-palmitoyl-2-oleoyl-*sn*-glycero-3 phosphoserine
PC: phosphatidylcholine
PE: phosphatidylethanolamine
PI: phosphatidylinositol
PPM: positioning of proteins in membranes
PS: phosphatidylserine
RMSD: Root Mean Square Deviation
SPH: sphingomyelin
TM: transmembrane
TMD: transmembrane domain

## Associated Content

### Supporting Information

Supplementary materials include 9 Tables and 11 Figures

#### Data and Software Availability

Information about simulation systems setup, parameterization etc. are provided in the Methods section. The following link is provided to access output frames of all the CG MD simulations: GitHub - etankol/CG_Adenosine_Receptors

## Acknowledgements

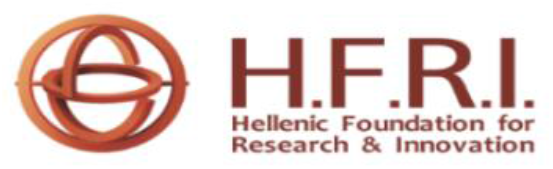

This research work represents part of the PhD thesis of ET and was supported by the Hellenic Foundation for Research and Innovation (HFRI) under the HFRI PhD Fellowship grant (Fellowship Number: 1619). This work was supported by computational time granted from the Greek Research & Technology Network (GRNET) in the National HPC facility ARIS (pr001007). We thank also Chiesi Hellas for the supporting of this research (SERG grant No 10354). We thank Margarita Stampelou for calculations performed for A_2A_R.

## Authors Contribution

AK conceived the project; AK, ET designed the research; ET performed the simulations and analyzed the results and RAC contributed to the improvement of the data appearance; AK interpreted the data; AK, ET wrote the manuscript and RAC criticized and edited it.

## TOC Graphic

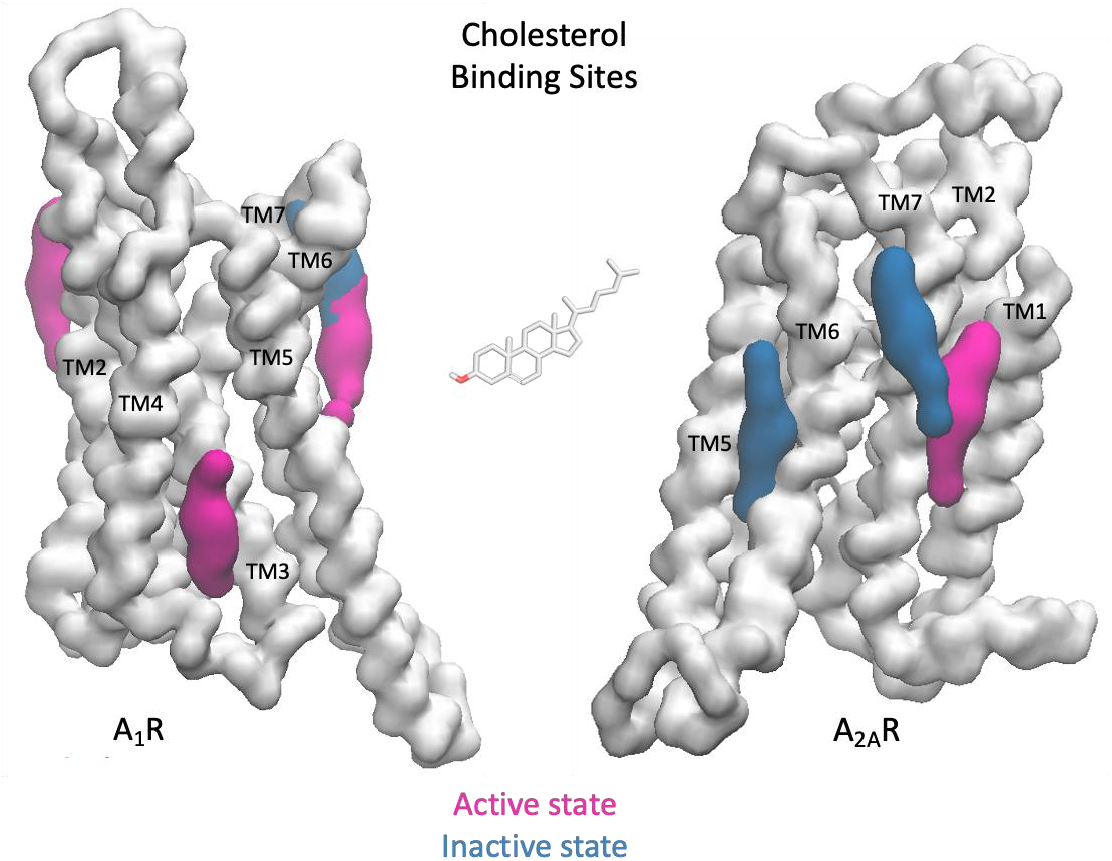

## References

1. Hauser AS, Attwood MM, Rask-Andersen M, Schiöth HB, Gloriam DE. Trends in GPCR drug discovery: New agents, targets and indications. Nat Rev Drug Discov. 2017;16(12):829–842. doi:10.1038/nrd.2017.178

2. Wacker D, Stevens RC, Roth BL. How Ligands Illuminate GPCR Molecular Pharmacology. Cell. 2017. doi:10.1016/j.cell.2017.07.009

3. Sriram K, Insel PA. G protein-coupled receptors as targets for approved drugs: How many targets and how many drugs? In: Molecular Pharmacology. ; 2018. doi:10.1124/mol.117.111062

4. Fredholm BB, Ijzerman AP, Jacobson KA, Linden J, Muller CE, Müller CE. International Union of Basic and Clinical Pharmacology. LXXXI. Nomenclature and Classification of Adenosine Receptors — An Update. Pharmacol Rev. 2011;63(1):1–34. doi:10.1124/pr.110.003285.1

5. Borea PA, Gessi S, Merighi S, Vincenzi F, Varani K. Pharmacology of Adenosine Receptors: The State of the Art. Physiol Rev. 2018;98(3):1591–1625. doi:10.1152/physrev.00049.2017

6. Lebon G, Warne T, Edwards PC, et al. Agonist-bound adenosine A2A receptor structures reveal common features of GPCR activation. Nature. 2011;474(7352):521–525. doi:10.1038/nature10136

7. García-Nafría J, Lee Y, Bai X, Carpenter B, Tate CG. Cryo-EM structure of the adenosine A2Areceptor coupled to an engineered heterotrimeric G protein. Elife. 2018;7:e35946:1-19. doi:10.7554/eLife.35946.001

8. Jaakola V-P, Griffith MT, Hanson MA, et al. The 2.6 Angstrom Crystal Structure of a Human A2A Adenosine Receptor Bound to an Antagonist. Science (80-). 2008;322(5905):1211–1217. doi:10.1126/science.1164772

9. Doré AS, Robertson N, Errey JC, et al. Structure of the adenosine A 2A receptor in complex with ZM241385 and the xanthines XAC and caffeine. Structure. 2011;19(9):1283–1293. doi:10.1016/j.str.2011.06.014

10. Liu W, Chun E, Thompson AA, et al. Structural Basis for Allosteric Regulation of GPCRs by Sodium Ions. Science (80-). 2012;337(6091):232–236. doi:10.1126/science.1219218

11. Sun B, Bachhawat P, Chu ML-H, et al. Crystal structure of the adenosine A 2A receptor bound to an antagonist reveals a potential allosteric pocket. Proc Natl Acad Sci. 2017;114(8):2066–2071. doi:10.1073/pnas.1621423114

12. Amelia T, Van Veldhoven JPD, Falsini M, et al. Crystal Structure and Subsequent Ligand Design of a Nonriboside Partial Agonist Bound to the Adenosine A2AReceptor. J Med Chem. 2021;64(7):3827–3842. doi:10.1021/acs.jmedchem.0c01856

13. Carpenter B, Nehmé R, Warne T, Leslie AGW, Tate CG. Structure of the adenosine A2A receptor bound to an engineered G protein. Nature. 2016;536(7614):104–107. doi:10.1038/nature18966

14. Glukhova A, Thal DM, Nguyen AT, et al. Structure of the Adenosine A1 Receptor Reveals the Basis for Subtype Selectivity. Cell. 2017;168(5):867-877.e13. doi:10.1016/j.cell.2017.01.042

15. Cheng RKY, Segala E, Robertson N, et al. Structures of Human A1 and A2A Adenosine Receptors with Xanthines Reveal Determinants of Selectivity. Structure. 2017;25(8):1275-1285.e4. doi:10.1016/j.str.2017.06.012

16. Draper-Joyce CJ, Khoshouei M, Thal DM, et al. Structure of the adenosine-bound human adenosine A1 receptor-Gi complex. Nature. 2018;558(7711):559–565. doi:10.1038/s41586-018-0236-6

17. Draper-Joyce CJ, Bhola R, Wang J, et al. Positive allosteric mechanisms of adenosine A1 receptor-mediated analgesia. Nature. 2021. doi:10.1038/s41586-021-03897-2

18. Lebon G, Warne T, Edwards PC, et al. Agonist-bound adenosine A2Areceptor structures reveal common features of GPCR activation. Nature. 2011;474(7352):521–525. doi:10.1038/nature10136

19. Dror RO, Pan AC, Arlow DH, et al. Pathway and mechanism of drug binding to G-protein-coupled receptors. Proc Natl Acad Sci U S A. 2011;108(32):13118–13123. doi:10.1073/pnas.1104614108

20. Luo J, Yang H, Song BL. Mechanisms and regulation of cholesterol homeostasis. Nat Rev Mol Cell Biol. 2020. doi:10.1038/s41580-019-0190-7

21. Duncan AL, Song W, Sansom MSP. Lipid-dependent regulation of ion channels and g protein-coupled receptors: Insights from structures and simulations. Annu Rev Pharmacol Toxicol. 2020;60:31–50. doi:10.1146/annurev-pharmtox-010919-023411

22. Genheden S, Essex JW, Lee AG. G protein coupled receptor interactions with cholesterol deep in the membrane. Biochim Biophys Acta - Biomembr. 2017. doi:10.1016/j.bbamem.2016.12.001

23. Gimpl G. Interaction of G protein coupled receptors and cholesterol. Chem Phys Lipids. 2016. doi:10.1016/j.chemphyslip.2016.04.006

24. Fantini J, Barrantes FJ. How cholesterol interacts with membrane proteins: an exploration of cholesterol-binding sites including CRAC, CARC, and tilted domains. Front Physiol. 2013;4:31. doi:10.3389/fphys.2013.00031

25. Cherezov V, Rosenbaum DM, Hanson MA, et al. High-resolution crystal structure of an engineered human beta2-adrenergic G protein-coupled receptor. Science (80-). 2007;318(5854):1258–1265. doi:10.1126/science.1150577

26. García-Nafría J, Tate CG. Cryo-Electron Microscopy: Moving Beyond X-Ray Crystal Structures for Drug Receptors and Drug Development. Annu Rev Pharmacol Toxicol. 2020;60(1):51–71. doi:10.1146/annurev-pharmtox-010919-023545

27. Segala E, Guo D, K. Y. Cheng R, et al. Controlling the Dissociation of Ligands from the Adenosine A2A Receptor through Modulation of Salt Bridge Strength. J Med Chem. 2016;59(13):6470–6479. doi:10.1021/acs.jmedchem.6b00653

28. Melnikov I, Polovinkin V, Kovalev K, et al. Fast iodide-SAD phasing for high-throughput membrane protein structure determination. Sci Adv. 2017;3(5):e1602952. doi:10.1126/sciadv.1602952

29. Batyuk A, Galli L, Ishchenko A, et al. Native phasing of x-ray free-electron laser data for a G protein–coupled receptor. Sci Adv. 2016. doi:10.1126/sciadv.1600292

30. Martin-Garcia JM, Conrad CE, Nelson G, et al. Serial millisecond crystallography of membrane and soluble protein microcrystals using synchrotron radiation. IUCrJ. 2017. doi:10.1107/S205225251700570X

31. Weinert T, Olieric N, Cheng R, et al. Serial millisecond crystallography for routine room-temperature structure determination at synchrotrons. Nat Commun. 2017. doi:10.1038/s41467-017-00630-4

32. Cheng RKY, Segala E, Robertson N, et al. Structures of Human A1and A2AAdenosine Receptors with Xanthines Reveal Determinants of Selectivity. Structure. 2017;25(8):1275–1285. doi:10.1016/j.str.2017.06.012

33. Broecker J, Morizumi T, Ou W-L, et al. High-throughput in situ X-ray screening of and data collection from protein crystals at room temperature and under cryogenic conditions. Nat Protoc. 2018;13(2):260–292. doi:10.1038/nprot.2017.135

34. Eddy MT, Lee MY, Gao ZG, et al. Allosteric Coupling of Drug Binding and Intracellular Signaling in the A2A Adenosine Receptor. Cell. 2018;172(1-2):68-80.e12. doi:10.1016/j.cell.2017.12.004

35. Rucktooa P, Cheng RKY, Segala E, et al. Towards high throughput GPCR crystallography: In Meso soaking of Adenosine A2A Receptor crystals. Sci Rep. 2018. doi:10.1038/s41598-017-18570-w

36. Martynowycz MW, Shiriaeva A, Ge X, et al. MicroED structure of the human adenosine receptor determined from a single nanocrystal in LCP. Proc Natl Acad Sci U S A. 2021. doi:10.1073/pnas.2106041118

37. Gater DL, Saurel O, Iordanov I, Liu W, Cherezov V, Milon A. Two classes of cholesterol binding sites for the β2AR revealed by thermostability and NMR. Biophys J. 2014. doi:10.1016/j.bpj.2014.10.011

38. Hanson MA, Cherezov V, Roth CB, et al. A specific cholesterol binding site is established by the 2.8 Å. Structure. 2009;16(6):897–905. doi:10.1016/j.str.2008.05.001.A

39. Hanson MA, Cherezov V, Griffith MT, et al. A Specific Cholesterol Binding Site Is Established by the 2.8 Å Structure of the Human β2-Adrenergic Receptor. Structure. 2008. doi:10.1016/j.str.2008.05.001

40. Ballesteros JA, Weinstein H. Receptor Molecular Biology. Vol 25.; 1995. doi:10.1016/S1043-9471(05)80049-7

41. Fantini J, Di Scala C, Baier CJ, Barrantes FJ. Molecular mechanisms of protein-cholesterol interactions in plasma membranes: Functional distinction between topological (tilted) and consensus (CARC/CRAC) domains. Chem Phys Lipids. 2016. doi:10.1016/j.chemphyslip.2016.02.009

42. Kiriakidi S, Kolocouris A, Liapakis G, Ikram S, Durdagi S, Mavromoustakos T. Effects of Cholesterol on GPCR Function: Insights from Computational and Experimental Studies. In: Advances in Experimental Medicine and Biology. Vol 1135.; 2019:89-103. doi:10.1007/978-3-030-14265-0_5

43. Sarkar P, Chattopadhyay A. Cholesterol interaction motifs in G protein-coupled receptors: Slippery hot spots? Wiley Interdiscip Rev Syst Biol Med. 2020. doi:10.1002/wsbm.1481

44. Jafurulla M, Tiwari S, Chattopadhyay A. Identification of cholesterol recognition amino acid consensus (CRAC) motif in G-protein coupled receptors. Biochem Biophys Res Commun. 2011. doi:10.1016/j.bbrc.2010.12.031

45. Di Scala C, Baier CJ, Evans LS, Williamson PTF, Fantini J, Barrantes FJ. Relevance of CARC and CRAC Cholesterol-Recognition Motifs in the Nicotinic Acetylcholine Receptor and Other Membrane-Bound Receptors. In: Current Topics in Membranes. Vol 80.; 2017:3-23. doi:10.1016/bs.ctm.2017.05.001

46. Baier CJ, Fantini J, Barrantes FJ. Disclosure of cholesterol recognition motifs in transmembrane domains of the human nicotinic acetylcholine receptor. Sci Rep. 2011. doi:10.1038/srep00069

47. Geiger J, Sexton R, Al-Sahouri Z, et al. Evidence that specific interactions play a role in the cholesterol sensitivity of G protein-coupled receptors. Biochim Biophys Acta - Biomembr. 2021;1863(9):183557. doi:10.1016/j.bbamem.2021.183557

48. Prasanna X, Chattopadhyay A, Sengupta D. Role of lipid-mediated effects in β2-adrenergic receptor dimerization. Adv Exp Med Biol. 2015. doi:10.1007/978-3-319-11280-0_16

49. Taghon GJ, Rowe JB, Kapolka NJ, Isom DG. Predictable cholesterol binding sites in GPCRs lack consensus motifs. Structure. 2021;29(5):499-506.e3. doi:10.1016/j.str.2021.01.004

50. Corradi V, Mendez-Villuendas E, Ingólfsson HI, et al. Lipid-Protein Interactions Are Unique Fingerprints for Membrane Proteins. ACS Cent Sci. 2018;4(6):709–717. doi:10.1021/acscentsci.8b00143

51. Abiko LA, Dias Teixeira R, Engilberge S, et al. Filling of a water-free void explains the allosteric regulation of the β1-adrenergic receptor by cholesterol. Nat Chem. August 2022. doi:10.1038/s41557-022-01009-9

52. Hanson MA, Cherezov V, Roth CB, et al. A specific cholesterol binding site is established by the 2.8 Å structure of the human β2 -adrenergic receptor in an alternate crystal form. Structure. 2009.

53. Warne T, Moukhametzianov R, Baker JG, et al. The structural basis for agonist and partial agonist action on a β1-adrenergic receptor. Nature. 2011. doi:10.1038/nature09746

54. Cang X, Yang L, Yang J, et al. Cholesterol-β1AR interaction versus cholesterol-β2AR interaction. Proteins Struct Funct Bioinforma. 2014. doi:10.1002/prot.24456

55. Enkavi G, Javanainen M, Kulig W, Róg T, Vattulainen I. Multiscale Simulations of Biological Membranes: The Challenge To Understand Biological Phenomena in a Living Substance. Chem Rev. 2019;119(9):5607–5774. doi:10.1021/acs.chemrev.8b00538

56. Simunovic M, Šarić A, Henderson JM, Lee KYC, Voth GA. Long-Range Organization of Membrane-Curving Proteins. ACS Cent Sci. 2017. doi:10.1021/acscentsci.7b00392

57. Hedger G, Sansom MSP. Lipid interaction sites on channels, transporters and receptors: Recent insights from molecular dynamics simulations. Biochim Biophys Acta Biomembr. 2016;1858(10):2390–2400. doi:10.1016/j.bbamem.2016.02.037

58. Lovera S, Cuzzolin A, Kelm S, De Fabritiis G, Sands ZA. Reconstruction of apo A2A receptor activation pathways reveal ligand-competent intermediates and state-dependent cholesterol hotspots. Sci Rep. 2019. doi:10.1038/s41598-019-50752-6

59. Guixà-González R, Albasanz JL, Rodriguez-Espigares I, et al. Membrane cholesterol access into a G-protein-coupled receptor. Nat Commun. 2017. doi:10.1038/ncomms14505

60. Lyman E, Higgs C, Kim B, et al. A Role for a Specific Cholesterol Interaction in Stabilizing the Apo Configuration of the Human A2A Adenosine Receptor. Structure. 2009. doi:10.1016/j.str.2009.10.010

61. Rouviere E, Arnarez C, Yang L, Lyman E. Identification of Two New Cholesterol Interaction Sites on the A2A Adenosine Receptor. Biophys J. 2017. doi:10.1016/j.bpj.2017.09.027

62. Yang L, Lyman E. Local Enrichment of Unsaturated Chains around the A2A Adenosine Receptor. Biochemistry. 2019. doi:10.1021/acs.biochem.9b00607

63. Song W, Yen H-Y, Robinson C V., Sansom MSP. State-dependent Lipid Interactions with the A2a Receptor Revealed by MD Simulations Using In Vivo-Mimetic Membranes. Structure. 2019;27(2):392-403.e3. doi:10.1016/j.str.2018.10.024

64. Lee JY, Lyman E. Predictions for cholesterol interaction sites on the A2A adenosine receptor. J Am Chem Soc. 2012. doi:10.1021/ja307532d

65. Manna M, Niemelä M, Tynkkynen J, et al. Mechanism of allosteric regulation of β2-adrenergic receptor by cholesterol. Elife. 2016;5:1–21. doi:10.7554/elife.18432

66. Schenone S, Brullo C, Musumeci F, Bruno O, Botta M. A1 receptors ligands: past, present and future trends. Curr Top Med Chem. 2010;10(9):878-901. http://www.ncbi.nlm.nih.gov/pubmed/20370661.

67. Shah RH, Frishman WH. Adenosine1 receptor antagonism: a new therapeutic approach for the treatment of decompensated heart failure. Cardiol Rev. 2009;17(3):125–131. doi:10.1097/CRD.0b013e31819f1a98

68. Wilson CN, Nadeem A, Spina D, Brown R, Page CP, Mustafa SJ. Adenosine receptors and asthma. In: Handbook of Experimental Pharmacology. Vol 193.; 2009:329-362. doi:10.1007/978-3-540-89615-9_11

69. Maemoto T, Tada M, Mihara T, et al. Pharmacological Characterization of FR194921, a New Potent, Selective, and Orally Active Antagonist for Central Adenosine A1 Receptors. J Pharmacol Sci. 2004;96(1):42–52. doi:10.1254/jphs.FP0040359

70. Jacobson KA, Tosh DK, Jain S, Gao Z-G. Historical and Current Adenosine Receptor Agonists in Preclinical and Clinical Development. Front Cell Neurosci. 2019;13(March):1-17. doi:10.3389/fncel.2019.00124

71. Jacobson KA, Civan MM. Ocular Purine Receptors as Drug Targets in the Eye. J Ocul Pharmacol Ther. 2016;32(8):534–547. doi:10.1089/jop.2016.0090

72. Mason PK, DiMarco JP. New pharmacological agents for arrhythmias. Circ Arrhythmia Electrophysiol. 2009. doi:10.1161/CIRCEP.109.884429

73. Greene SJ, Sabbah HN, Butler J, et al. Partial adenosine A1 receptor agonism: a potential new therapeutic strategy for heart failure. Heart Fail Rev. 2016;21(1):95–102. doi:10.1007/s10741-015-9522-7

74. Ingólfsson HI, Arnarez C, Periole X, Marrink SJ. Computational “microscopy” of cellular membranes. J Cell Sci. 2016;129(2):257–268. doi:10.1242/jcs.176040

75. Ingólfsson HI, Melo MN, van Eerden FJ, et al. Lipid Organization of the Plasma Membrane. J Am Chem Soc. 2014;136(41):14554–14559. doi:10.1021/ja507832e

76. Koldsø H, Sansom MSP. Organization and Dynamics of Receptor Proteins in a Plasma Membrane. J Am Chem Soc. 2015;137(46):14694–14704. doi:10.1021/jacs.5b08048

77. Harayama T, Riezman H. Understanding the diversity of membrane lipid composition. Nat Rev Mol Cell Biol. 2018. doi:10.1038/nrm.2017.138

78. Marrink SJ, Corradi V, Souza PCT, Ingólfsson HI, Tieleman DP, Sansom MSP. Computational Modeling of Realistic Cell Membranes. Chem Rev. 2019. doi:10.1021/acs.chemrev.8b00460

79. Ansell TB, Curran L, Horrell MR, et al. Relative Affinities of Protein–Cholesterol Interactions from Equilibrium Molecular Dynamics Simulations. J Chem Theory Comput. 2021;17(10):6548–6558. doi:10.1021/acs.jctc.1c00547

80. Lee M-Y, Geiger J, Ishchenko A, et al. Harnessing the power of an X-ray laser for serial crystallography of membrane proteins crystallized in lipidic cubic phase. IUCrJ. 2020;7(6):976–984. doi:10.1107/S2052252520012701

81. Ihara K, Hato M, Nakane T, et al. Isoprenoid-chained lipid EROCOC17+4: a new matrix for membrane protein crystallization and a crystal delivery medium in serial femtosecond crystallography. Sci Rep. 2020;10(1). doi:10.1038/s41598-020-76277-x

82. McGraw C, Yang L, Levental I, Lyman E, Robinson AS. Membrane cholesterol depletion reduces downstream signaling activity of the adenosine A 2A receptor. Biochim Biophys Acta - Biomembr. 2019;1861(4):760–767. doi:10.1016/j.bbamem.2019.01.001

83. Payandeh J, Volgraf M. Ligand binding at the protein–lipid interface: strategic considerations for drug design. Nat Rev Drug Discov. 2021. doi:10.1038/s41573-021-00240-2

84. Szlenk CT, Gc JB, Natesan S. Does the Lipid Bilayer Orchestrate Access and Binding of Ligands to Transmembrane Orthosteric/Allosteric Sites of G Protein-Coupled Receptors? Mol Pharmacol. 2019;96(5):527–541. doi:10.1124/mol.118.115113

85. Jaakola VP, Griffith MT, Hanson MA, et al. The 2.6 angstrom crystal structure of a human A2Aadenosine receptor bound to an antagonist. Science (80-). 2008;322(5905):1211–1217. doi:10.1126/science.1164772

86. Waterhouse A, Bertoni M, Bienert S, et al. {SWISS-MODEL}: homology modelling of protein structures and complexes. Nucleic Acids Res. 2018;46(W1):W296--W303.

87. Lomize MA, Pogozheva ID, Joo H, Mosberg HI, Lomize AL. {OPM} database and {PPM} web server: resources for positioning of proteins in membranes. Nucleic Acids Res. 2012;40(Database issue):D370--6.

88. Kaminski GA, Friesner RA, Tirado-Rives J, Jorgensen WL. Evaluation and reparametrization of the OPLS-AA force field for proteins via comparison with accurate quantum chemical calculations on peptides. J Phys Chem B. 2001;105(28):6474–6487. doi:10.1021/jp003919d

89. Touw WG, Baakman C, Black J, et al. A series of PDB-related databanks for everyday needs. Nucleic Acids Res. 2015;43(D1):D364–D368. doi:10.1093/nar/gku1028

90. Poh SC, Chew SE. Dictionary of Protein Secondary Structure: Pattern Recognition of Hydrogen-Bonded and Geometrical Features. Biopolymers. 1983;22:2577–2637.

91. Marrink SJ, Risselada HJ, Yefimov S, Tieleman DP, De Vries AH. The MARTINI force field: Coarse grained model for biomolecular simulations. J Phys Chem B. 2007;111(27):7812–7824. doi:10.1021/jp071097f

92. Monticelli L, Kandasamy SK, Periole X, Larson RG, Tieleman DP, Marrink SJ. The MARTINI coarse-grained force field: Extension to proteins. J Chem Theory Comput. 2008;4(5):819–834. doi:10.1021/ct700324x

93. De Jong DH, Singh G, Bennett WFD, et al. Improved parameters for the martini coarse-grained protein force field. J Chem Theory Comput. 2013;9(1):687–697. doi:10.1021/ct300646g

94. Wassenaar TA, Ingólfsson HI, Böckmann RA, Tieleman DP, Marrink SJ. Computational lipidomics with insane: A versatile tool for generating custom membranes for molecular simulations. J Chem Theory Comput. 2015;11(5):2144–2155. doi:10.1021/acs.jctc.5b00209

95. Berendsen HJC, van der Spoel D, van Drunen R. GROMACS: A message-passing parallel molecular dynamics implementation. Comput Phys Commun. 1995;91(1-3):43–56. doi:10.1016/0010-4655(95)00042-E

96. Pronk S, Páll S, Schulz R, et al. GROMACS 4.5: a high-throughput and highly parallel open source molecular simulation toolkit. Bioinformatics. 2013;29(7):845–854. doi:10.1093/bioinformatics/btt055

97. Darden T, York D, Pedersen L. Particle mesh Ewald: An Nlog(N) method for Ewald sums in large systems. J Chem Phys. 1993;98(12):10089–10092. doi:10.1063/1.464397

98. Bussi G, Donadio D, Parrinello M. Canonical sampling through velocity rescaling. J Chem Phys. 2007;126(1):014101. doi:10.1063/1.2408420

99. Berendsen HJC, Postma JPM, Van Gunsteren WF, Dinola A, Haak JR. Molecular dynamics with coupling to an external bath. J Chem Phys. 1984;81(8):3684–3690. doi:10.1063/1.448118

100. Martoňák R, Laio A, Parrinello M. Predicting Crystal Structures: The Parrinello-Rahman Method Revisited. Phys Rev Lett. 2003;90(7):4. doi:10.1103/PhysRevLett.90.075503

101. Humphrey W, Dalke A, Schulten K. VMD: Visual Molecular Dynamics. J Mol Graph. 1996;14(1):33–38. doi:10.1016/0263-7855(96)00018-5

102. García AE, Stiller L. Computation of the mean residence time of water in the hydration shells of biomolecules. J Comput Chem. 1993;14(11):1396–1406. doi:10.1002/jcc.540141116

103. Duncan AL, Corey RA, Sansom MSP. Defining how multiple lipid species interact with inward rectifier potassium (Kir2) channels. Proc Natl Acad Sci U S A. 2020;117(14):7803–7813. doi:10.1073/pnas.1918387117

104. Barbera N, Ayee MAA, Akpa BS, Levitan I. Molecular Dynamics Simulations of Kir2.2 Interactions with an Ensemble of Cholesterol Molecules. Biophys J. 2018;115(7):1264–1280. doi:10.1016/j.bpj.2018.07.041

105. Hedger G, Koldsø H, Chavent M, Siebold C, Rohatgi R, Sansom MSP. Cholesterol Interaction Sites on the Transmembrane Domain of the Hedgehog Signal Transducer and Class F G Protein-Coupled Receptor Smoothened. Struct Des. 2019;27:549-559.e2. doi:10.1016/j.str.2018.11.003

106. Song W, Corey RA, Ansell TB, et al. PyLipID: A Python Package for Analysis of Protein–Lipid Interactions from Molecular Dynamics Simulations. J Chem Theory Comput. 2022;0(0):acs.jctc.1c00708. doi:10.1021/acs.jctc.1c00708

107. Song W, Corey RA, Ansell TB, et al. Wanling Song, Robin A. Corey, T. Bertie Ansell, C. Keith Cassidy, Michael R. Horrell, Anna L. Duncan, Phillip J. Stansfeld, and Mark S. P. Sansom. 2021.

108. Isralewitz B, Gao M, Schulten K. Steered molecular dynamics and mechanical functions of proteins. Curr Opin Struct Biol. 2001. doi:10.1016/S0959-440X(00)00194-9

109. Van Meer G, de Kroon Aipmpm. Lipid map of the mammalian cell. J Cell Sci. 2011;124(1):5–8. doi:10.1242/jcs.071233

110. Coskun Ü, Simons K. Cell membranes: The lipid perspective. Structure. 2011;19(11):1543–1548. doi:10.1016/j.str.2011.10.010

111. Escribá P V., Busquets X, Inokuchi JI, et al. Membrane lipid therapy: Modulation of the cell membrane composition and structure as a molecular base for drug discovery and new disease treatment. Prog Lipid Res. 2015. doi:10.1016/j.plipres.2015.04.003

112. Casares D, Escribá P V., Rosselló CA. Membrane lipid composition: Effect on membrane and organelle structure, function and compartmentalization and therapeutic avenues. Int J Mol Sci. 2019. doi:10.3390/ijms20092167

113. Chini B, Parenti M. G-protein-coupled receptors, cholesterol and palmitoylation: Facts about fats. J Mol Endocrinol. 2009. doi:10.1677/JME-08-0114

114. Oates J, Watts A. Uncovering the intimate relationship between lipids, cholesterol and GPCR activation. Curr Opin Struct Biol. 2011. doi:10.1016/j.sbi.2011.09.007

115. Gahbauer S, Böckmann RA. Membrane-mediated oligomerization of G protein coupled receptors and its implications for GPCR function. Front Physiol. 2016. doi:10.3389/fphys.2016.00494

116. Pucadyil TJ, Chattopadhyay A. Cholesterol modulates ligand binding and G-protein coupling to serotonin1A receptors from bovine hippocampus. Biochim Biophys Acta - Biomembr. 2004. doi:10.1016/j.bbamem.2004.03.010

117. Paila YD, Jindal E, Goswami SK, Chattopadhyay A. Cholesterol depletion enhances adrenergic signaling in cardiac myocytes. Biochim Biophys Acta - Biomembr. 2011. doi:10.1016/j.bbamem.2010.09.006

118. Yeliseev A, Iyer MR, Joseph TT, et al. Cholesterol as a modulator of cannabinoid receptor CB2 signaling. Sci Rep. 2021. doi:10.1038/s41598-021-83245-6

119. Potter RM, Harikumar KG, Wu SV, Miller LJ. Differential sensitivity of types 1 and 2 cholecystokinin receptors to membrane cholesterol. J Lipid Res. 2012. doi:10.1194/jlr.M020065

120. Oddi S, Dainese E, Fezza F, et al. Functional characterization of putative cholesterol binding sequence (CRAC) in human type-1 cannabinoid receptor. J Neurochem. 2011. doi:10.1111/j.1471-4159.2010.07041.x

121. Qiu Y, Wang Y, Law PY, Chen HZ, Loh HH. Cholesterol regulates μ-opioid receptor-induced β-arrestin 2 translocation to membrane lipid rafts. Mol Pharmacol. 2011. doi:10.1124/mol.110.070870

122. Muth S, Fries A, Gimpl G. Cholesterol-induced conformational changes in the oxytocin receptor. Biochem J. 2011;437(3):541–553. doi:10.1042/BJ20101795

123. Calmet P, Cullin C, Cortès S, et al. Cholesterol impacts chemokine CCR5 receptor ligand-binding activity. FEBS J. 2020. doi:10.1111/febs.15145

124. van Aalst E, Wylie BJ. Cholesterol Is a Dose-Dependent Positive Allosteric Modulator of CCR3 Ligand Affinity and G Protein Coupling. Front Mol Biosci. 2021;8. doi:10.3389/fmolb.2021.724603

125. Gimpl G, Klein U, Reilander H, Fahrenholz F. Expression of the Human Oxytocin Receptor in Baculovirus-infected Insect Cells: High-Affinity Binding Is Induced by a Cholesterol-Cyclodextrin Complex. Biochemistry. 1995. doi:10.1021/bi00042a010

126. Krimmer SG, Cramer J, Schiebel J, Heine A, Klebe G. How Nothing Boosts Affinity: Hydrophobic Ligand Binding to the Virtually Vacated S1′ Pocket of Thermolysin. J Am Chem Soc. 2017. doi:10.1021/jacs.7b05028

127. Abiko LA, Grahl A, Grzesiek S. High Pressure Shifts the β1-Adrenergic Receptor to the Active Conformation in the Absence of G Protein. J Am Chem Soc. 2019. doi:10.1021/jacs.9b06042

128. Liu X, Kaindl J, Korczynska M, et al. An allosteric modulator binds to a conformational hub in the β2 adrenergic receptor. Nat Chem Biol. 2020. doi:10.1038/s41589-020-0549-2

129. Lu J, Byrne N, Wang J, et al. Structural basis for the cooperative allosteric activation of the free fatty acid receptor GPR40. Nat Struct Mol Biol. 2017. doi:10.1038/nsmb.3417

130. Zhang D, Gao ZG, Zhang K, et al. Two disparate ligand-binding sites in the human P2Y1 receptor. Nature. 2015. doi:10.1038/nature14287

131. Xiao P, Yan W, Gou L, et al. Ligand recognition and allosteric regulation of DRD1-Gs signaling complexes. Cell. 2021. doi:10.1016/j.cell.2021.01.028

132. Robertson N, Rappas M, Doré AS, et al. Structure of the complement C5a receptor bound to the extra-helical antagonist NDT9513727. Nature. 2018. doi:10.1038/nature25025

133. Cheng RKY, Fiez-Vandal C, Schlenker O, et al. Structural insight into allosteric modulation of protease-activated receptor 2. Nature. 2017. doi:10.1038/nature22309

134. Thal DM, Glukhova A, Sexton PM, Christopoulos A. Structural insights into G-protein-coupled receptor allostery. Nature. 2018. doi:10.1038/s41586-018-0259-z

135. Obi P, Natesan S. Membrane Lipids Are an Integral Part of Transmembrane Allosteric Sites in GPCRs: A Case Study of Cannabinoid CB1 Receptor Bound to a Negative Allosteric Modulator, ORG27569, and Analogs. J Med Chem. 2022;0(0). doi:10.1021/acs.jmedchem.2c00946

136. Sejdiu BI, Tieleman DP. Lipid-Protein Interactions Are a Unique Property and Defining Feature of G Protein-Coupled Receptors. Biophysj. 2020;118(8):1887–1900. doi:10.1016/j.bpj.2020.03.008

137. Kiriakidi S, Chatzigiannis C, Papaemmanouil C, Tzakos AG, Mavromoustakos T. Exploring the role of the membrane bilayer in the recognition of candesartan by its GPCR AT1 receptor. Biochim Biophys Acta - Biomembr. 2020. doi:10.1016/j.bbamem.2019.183142

138. Shao Z, Yan W, Chapman K, et al. Structure of an allosteric modulator bound to the CB1 cannabinoid receptor. Nat Chem Biol. 2019. doi:10.1038/s41589-019-0387-2

