## Supplementary material for "Comparative Study of State-Dependent Cholesterol Binding Sites in Adenosine A_2A_ and A_1_ Receptors Using Coarse-Grained Molecular Dynamics Simulations in Biologically Relevant Membranes": GitHub - etankol/CG_Adenosine_Receptors

Active and inactive conformation of A_2A_R in POPC-CHOL membrane

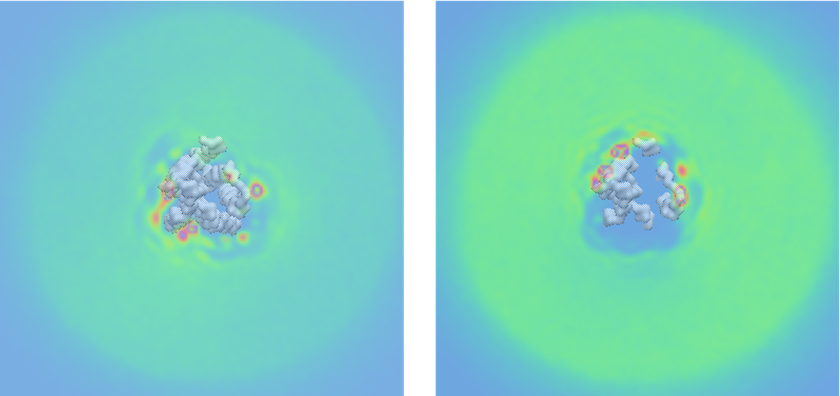
**A	C**

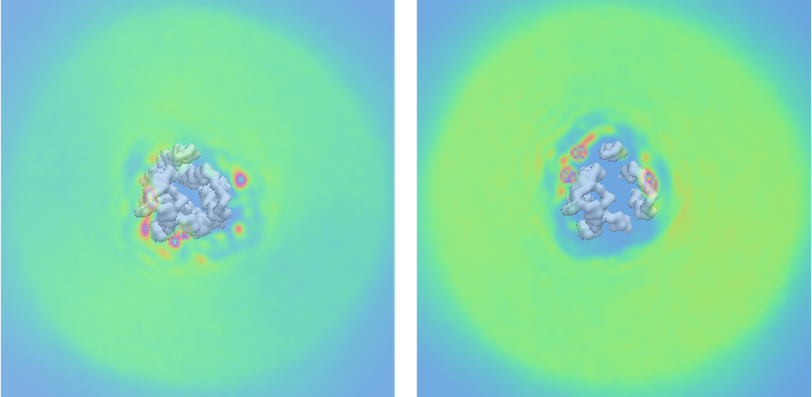

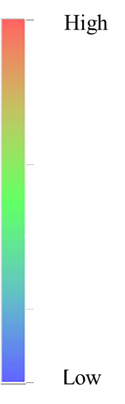

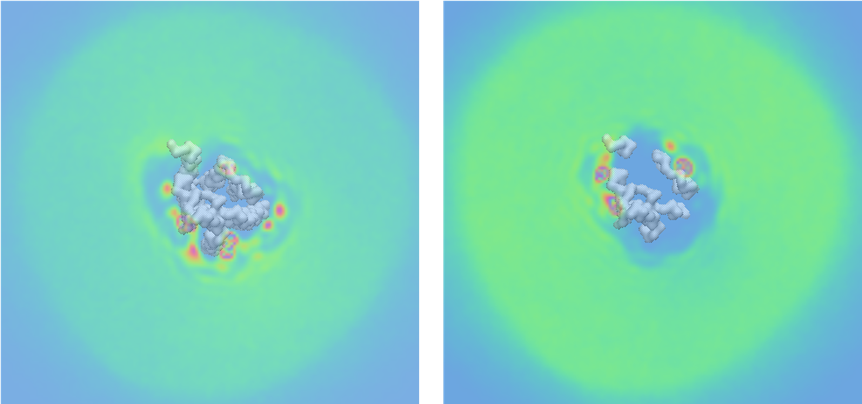
**B	D**

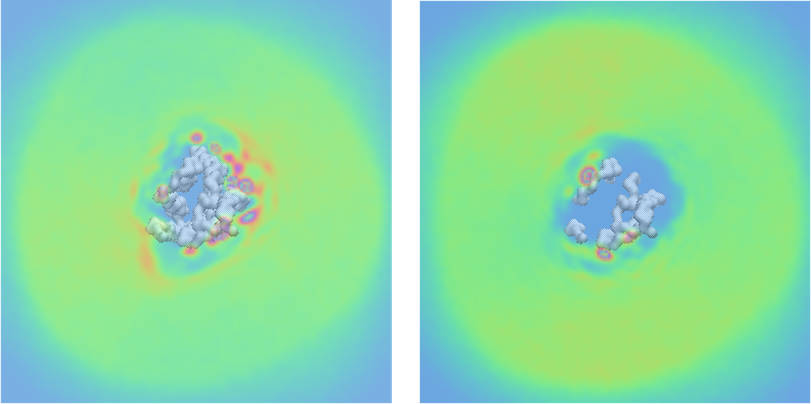

**Supplementary Figure 1.** CHOL density maps averaged for three repeats using the last 8 μs of the 10 μs-CG MD simulations of each simulated system with Martini force field ^90–92^ for the analysis. (A) CHOL density maps of the active state of A_2A_R in membrane composed by POPC – 20% CHOL. (B) CHOL density maps of the inactive state of A_2A_R in membrane composed by POPC – 20% CHOL (C) CHOL density maps of the active state of A_2A_R in plasma mimetic membrane. (D) CHOL density maps of the inactive state of A_2A_R in plasma mimetic membrane

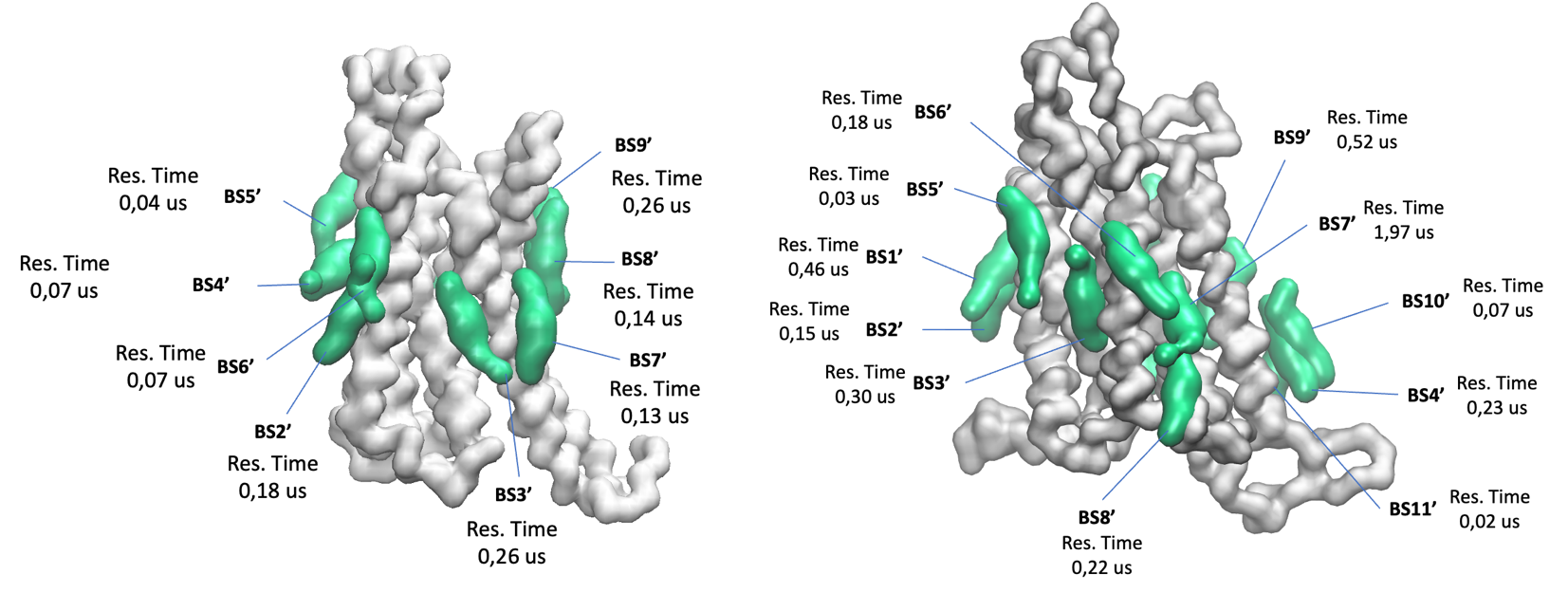

**A**

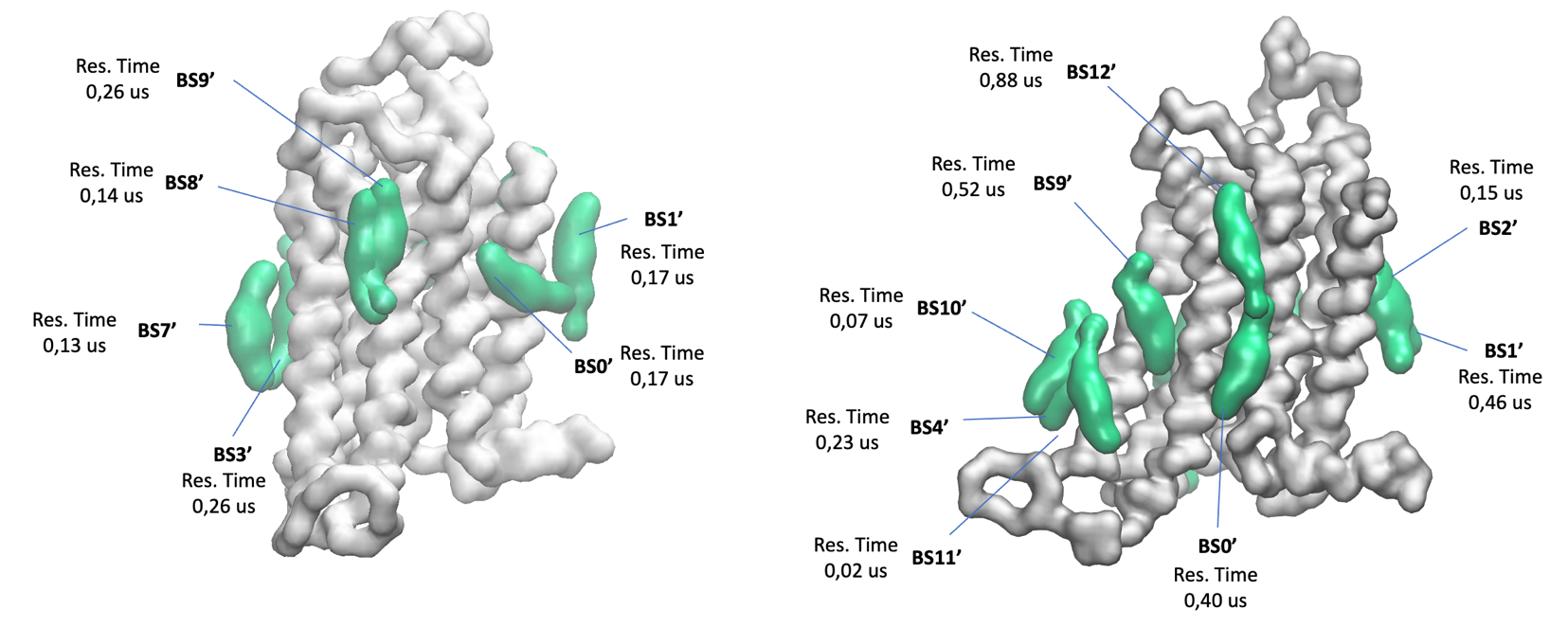
**B**

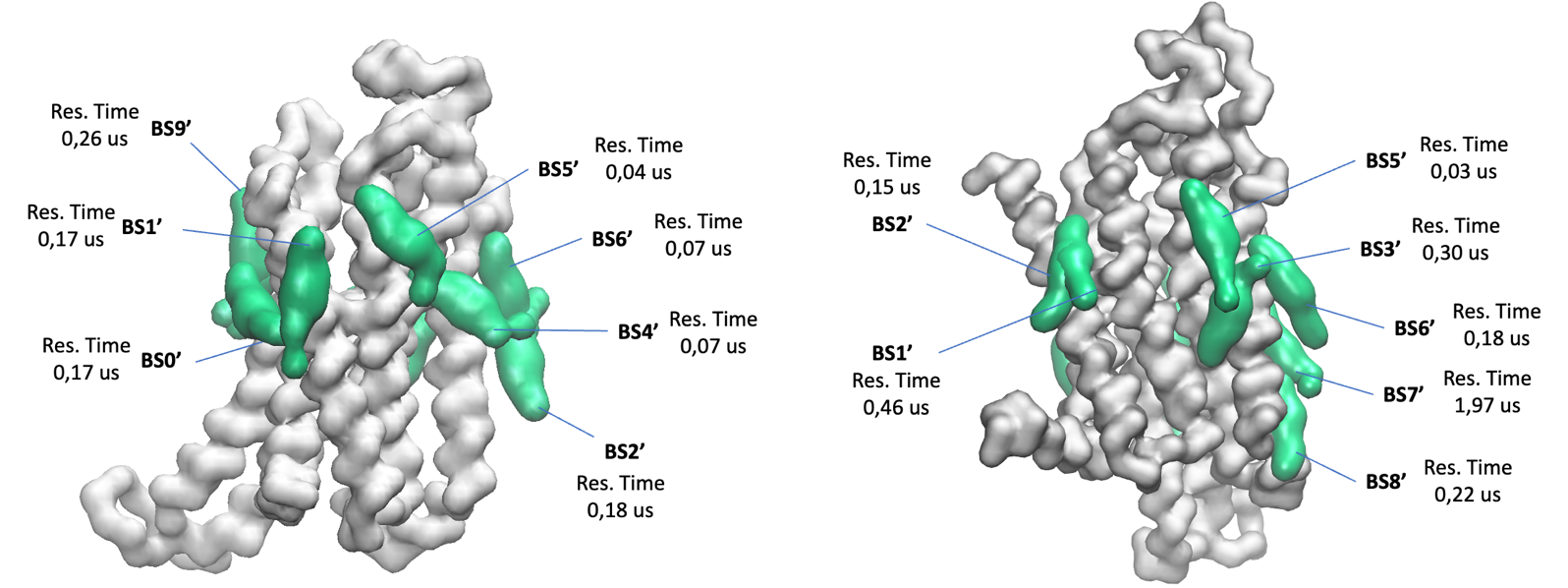
**C**

**Supplementary Figure 2.** Comparison of CHOL binding sites and their residence time between active (BS0′-BS9′) and inactive state (BS0′-BS12′) of A_2A_R in POPC–20 % CHOL membrane using the last 8 μs of the 10 μs-CG MD simulations of each simulated system with Martini force field ^90–92^ for the analysis. Different sides of the receptor are shown in A-C with the active state of the receptor shown in the left side and the inactive the in right side. Receptor is shown in white surface and CHOL representative poses for each binding site in green surface.

A

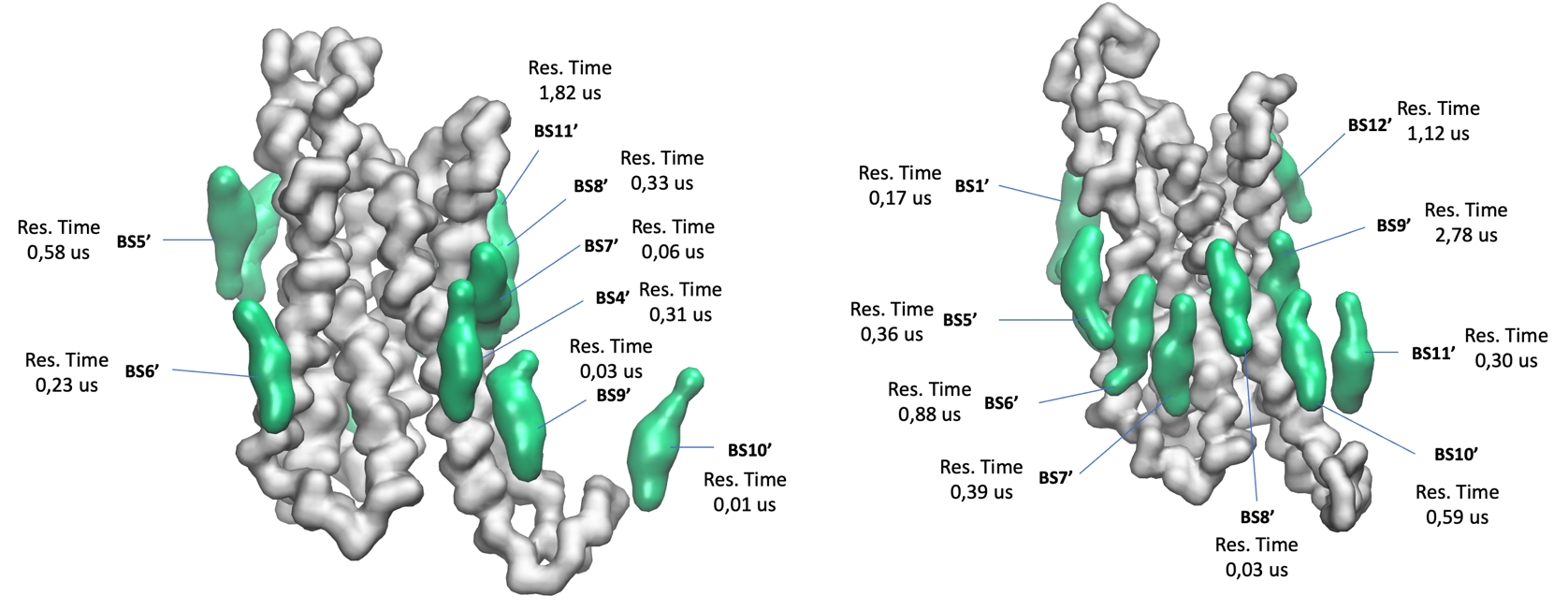

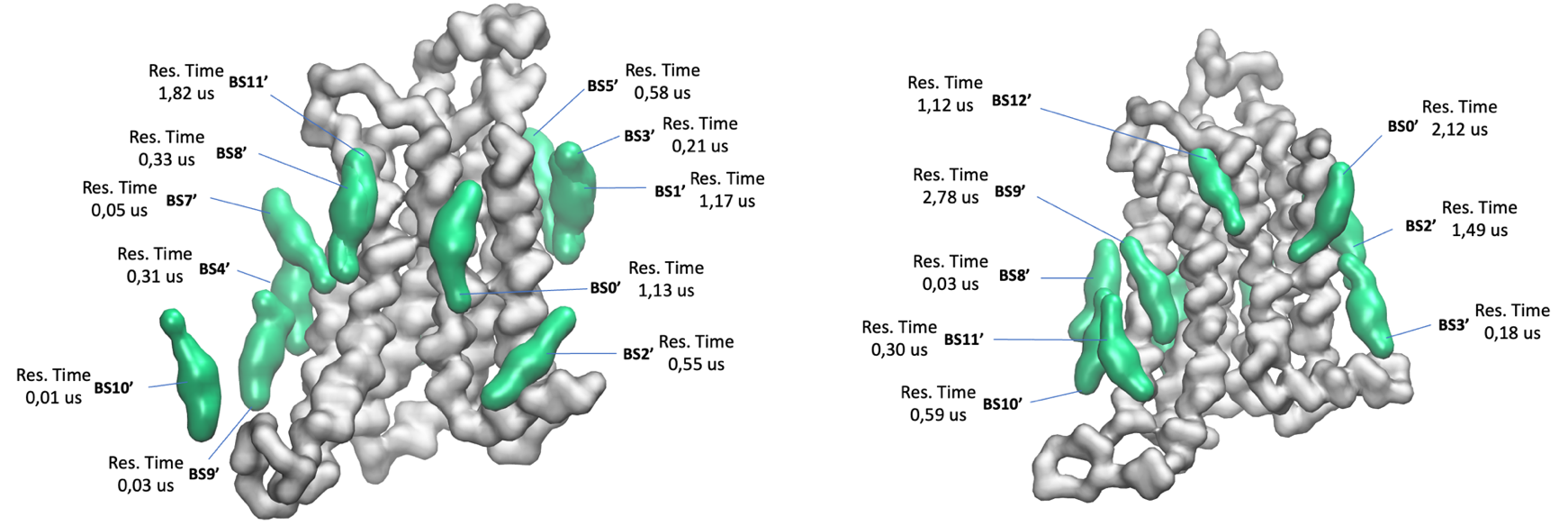
B

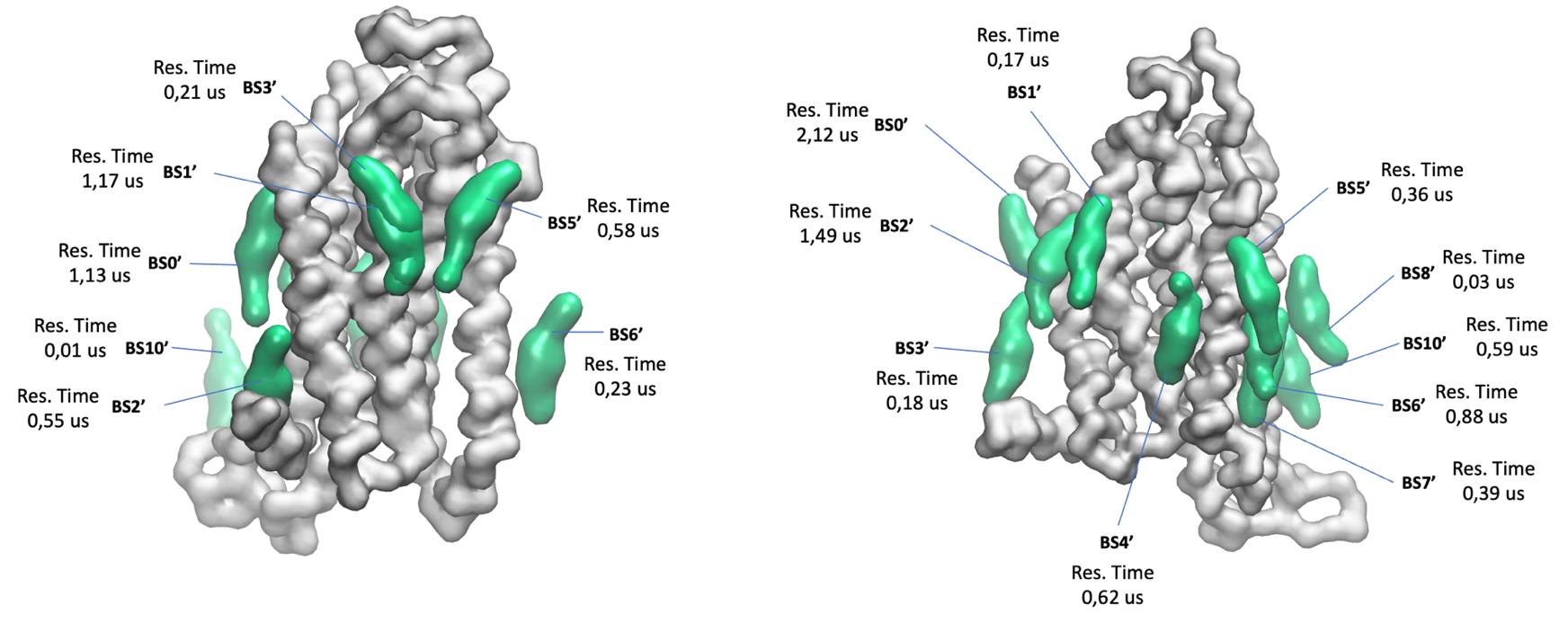
C

**Supplementary Figure 3.** Comparison of CHOL binding sites and their residence time between active (BS0′-BS11′) and inactive state (BS0′-BS12′) of A_2A_R in plasma mimetic membrane using the last 8 μs of the 10 μs-CG MD simulations of each simulated system with Martini force field ^90–92^ for the analysis. Different sides of the receptor are shown in A-C with the active state of the receptor shown in the left side and the inactive the in right side. Receptor is shown in white surface and CHOL representative poses for each binding site in green surface.

Active and inactive conformation of A_1_R in POPC-CHOL membrane

**A**

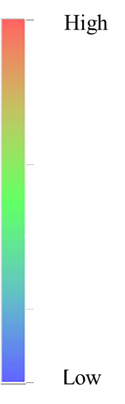

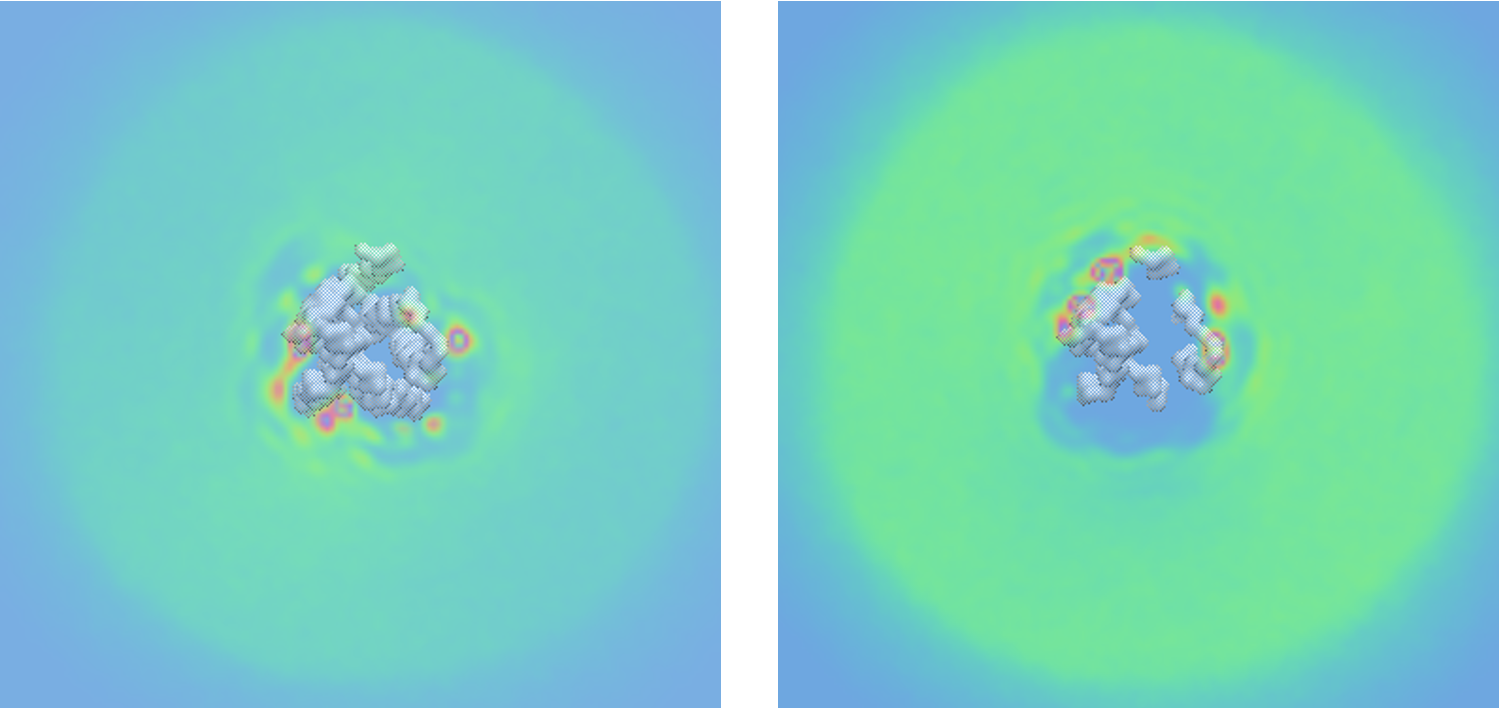

**Β**

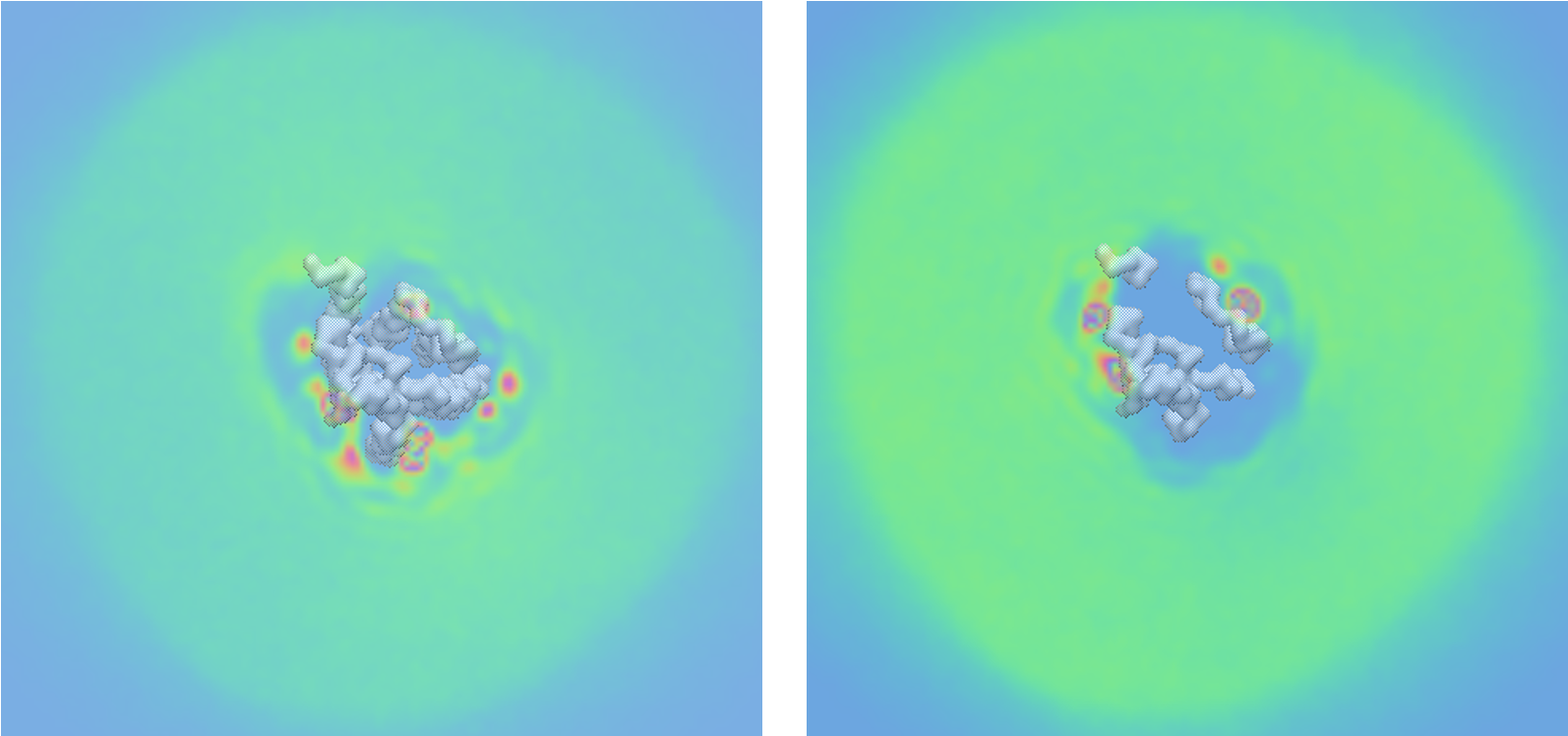

**Supplementary Figure 4.** CHOL density maps averaged for three repeats using the last 8 μs of the 10 μs-CG MD simulations of each simulated system with Martini force field ^90–92^ for the analysis. (A) CHOL density maps of the active state of A_1_R in membrane composed by POPC ‒ 20% CHOL. (B) CHOL density maps of the inactive state of A_1_R in membrane composed by POPC ‒ 20% CHOL.

Inactive conformation of A_1_R in POPC‒CHOL membrane

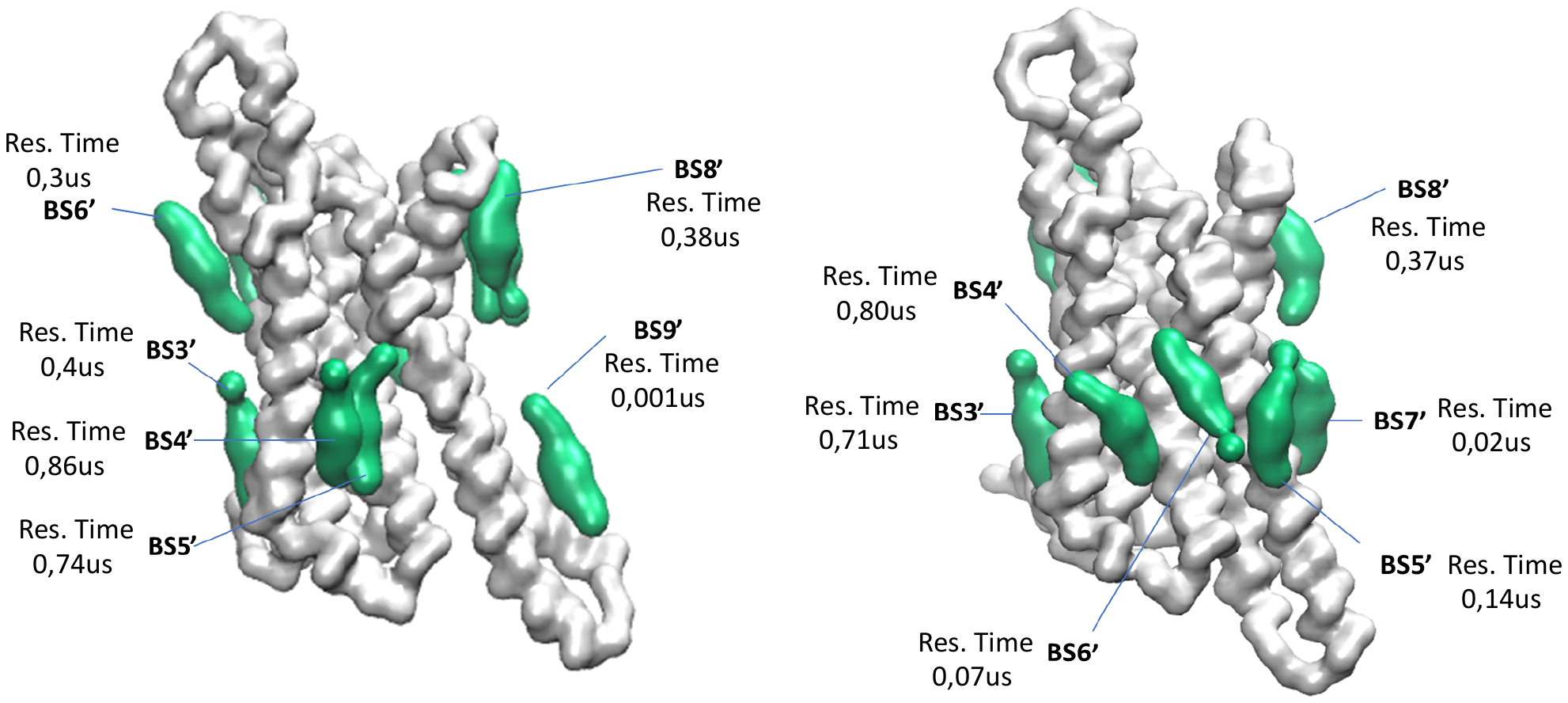
**A**

**B**

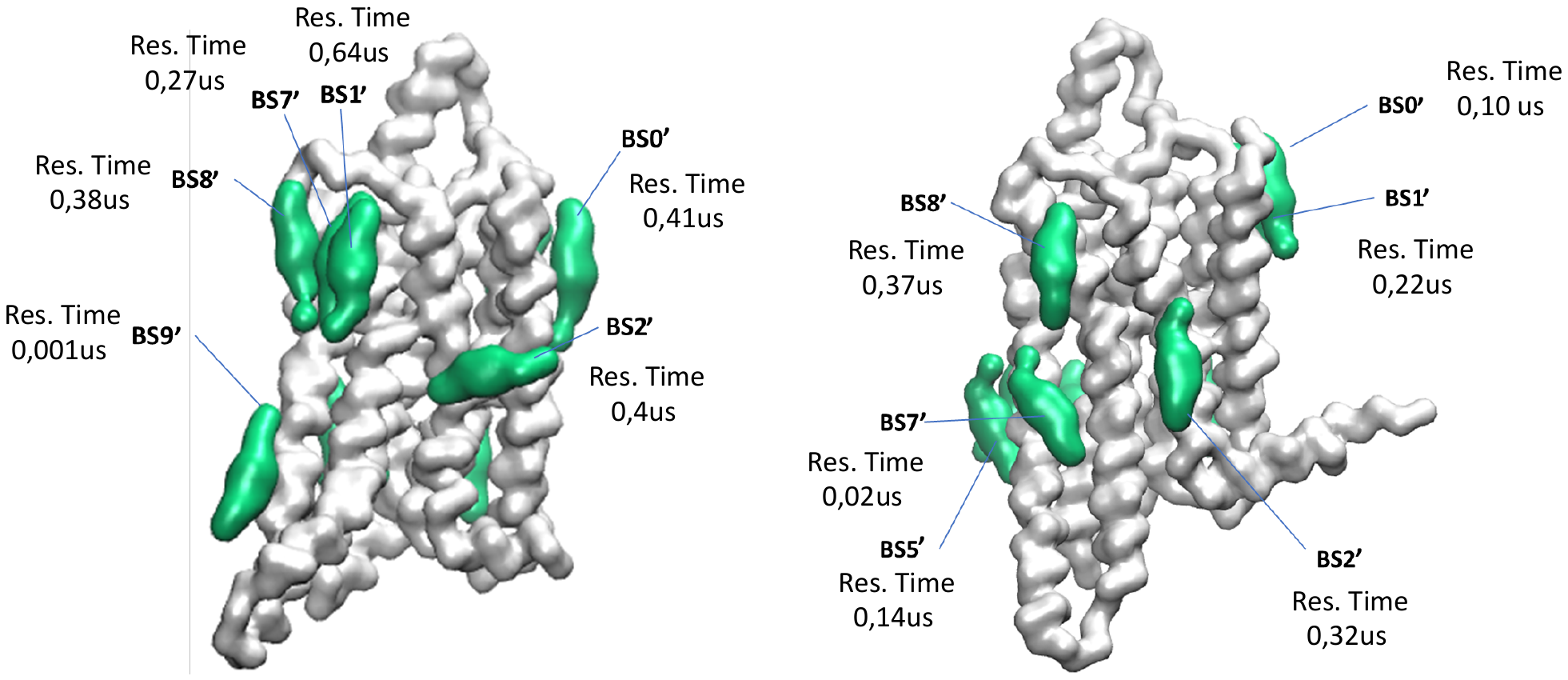

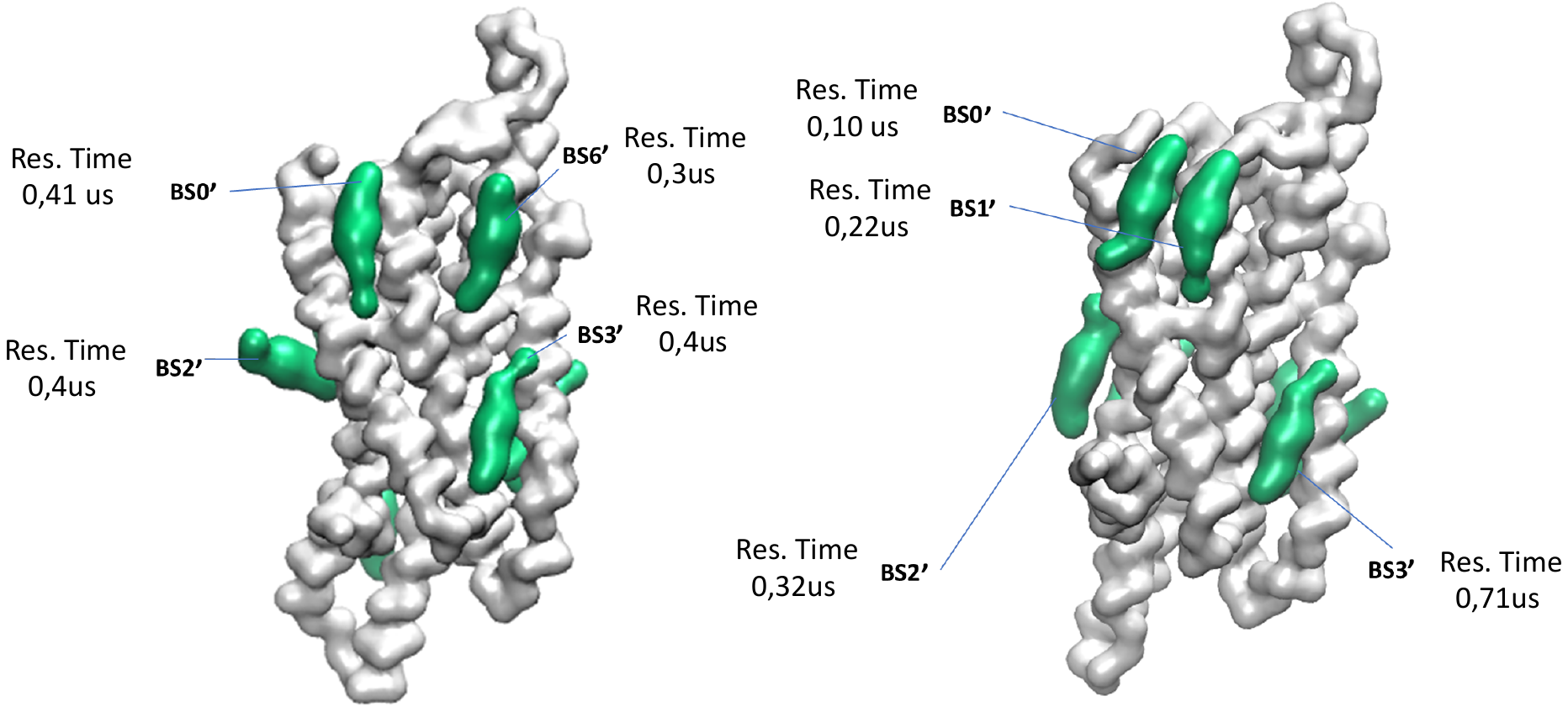
**C**

**Supplementary Figure 5.** Comparison of CHOL binding sites between active (BS0′-BS9′) and inactive state (BS0′-BS8′) of A_1_R in POPC‒20 % CHOL membrane using the last 8 μs of the 10 μs-CG MD simulations of each simulated system with Martini force field ^90–92^ for the analysis. Different sides of the receptor are shown in A, B and C with the active state of the receptor shown in the left side and the inactive the in right side. Receptor is shown in white surface and CHOL representative poses for each binding site in green surface.

Active and inactive conformation of A_1_R in POPC‒CHOL‒PIP_2_ membrane

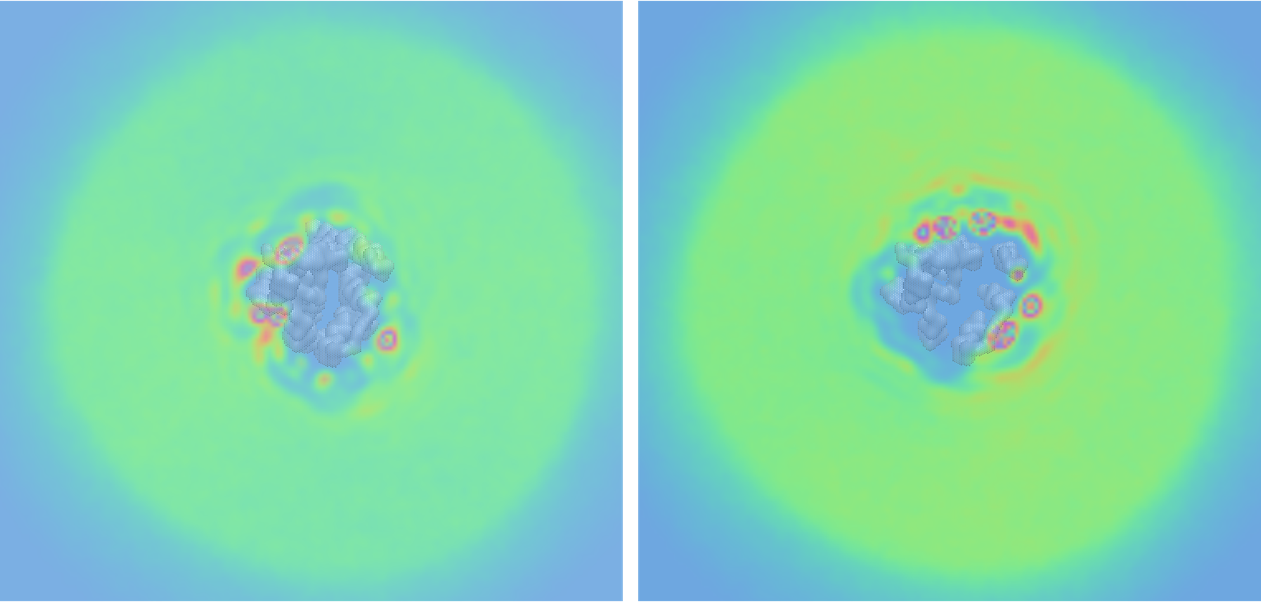
**A**

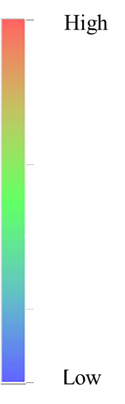

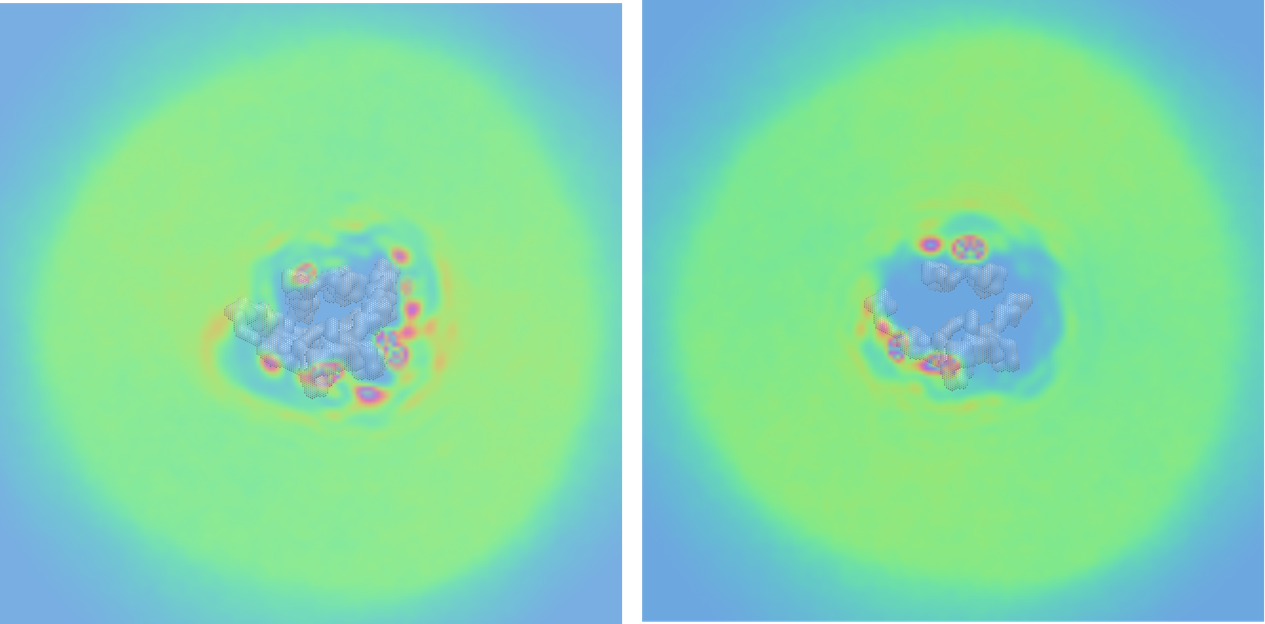
**B**

**Supplementary Figure 6.** CHOL density maps averaged for three repeats using the last 8 μs of the 10 μs-CG MD simulations of each simulated system with Martini force field ^90–92^ for the analysis.^90–92^ (A) CHOL density maps of the active state of A_1_R in membrane composed by POPC, CHOL and 10% PIP_2_ . (B) CHOL density maps of the inactive state of A_1_R in membrane composed by POPC, CHOL and 10% PIP_2_.

Active conformation of A_1_R in POPC‒CHOL‒PIP_2_ membrane

The analysis of the CHOL contact frequencies to active A_1_R resulted in 9 binding sites, BS0′-BS8′, which are shown in Figure S7 (left part) and their residence time shown in both Figure S7 (left part) and Table S6. In Figure S8A, are shown the 8 distinct binding sites, BS0-BS7 to active A_1_R, we ended up, after the visual inspection of the initially 9 binding sites identified by PyLipID ^105^ and, are shown the residues with more than 1 μs residence time of CHOL in these binding sites too.

BS0 is located in the upper part of the receptor between TM1-TM2. BS1 is perpendicular to the membrane’s plane as BS2 in the POPC-CHOL membrane does. BS2 is a continuation of BS0 and occupies the middle area of TM1-TM2. BS3 lies in the cavity formed by the lower part of TM2-TM3. BS4, which is the only binding site with CHOL residence time more than 1us (4.76μs), lies in the cavity formed by TM2-TM3 and TM4. Next to BS4 is BS5 which lies between TM3-TM5, BS6 lies between TM3-TM4 and BS7 occupies the space of the upper part of TM5-TM6. Residues with high CHOL residence time for the active conformation of A_1_R in the membrane composed of POPC-CHOL-PIP_2_ are F47^2.42^ (1.16 μs), V53^2.48^ (1.76 μs), L96^3.41^ (1.36 μs), L99^3.44^ (1.42 μs), A100^3.45^ (1.28 μs), V103^3.48^ (1.58 μs), A124^4.42^ (1.47 μs), A127^4.45^ (1.86 μs), C131^4.49^ (1.33 μs), Y271^7.36^ (1.83 μs).

Inactive conformation of A_1_R in POPC‒CHOL‒PIP_2_ membrane

In Figure S8B are shown the 10 identified distinct binding sites BS0-BS9, while Figure S7 shows the initial 11 binding sites, BS0′-BS10′ (identified by PyLipID) with residence time of CHOL in each of these sites shown also in Table S7. In Figure 7B are shown the residues with more than 1 μs CHOL residence time, which are I31^1.54^ (1.59 μs), V34^1.57^ (1.10 μs), K35^1.58^ (2.10 μs), C46^2.41^ (1.69 μs), V49^2.44^ (2.12 μs), S50^2.45^ (1.32 μs), V53^2.48^ (1.89 μs), A123^4.42^ (1.31 μs), V287^7.52^ (1.51 μs).

**A**

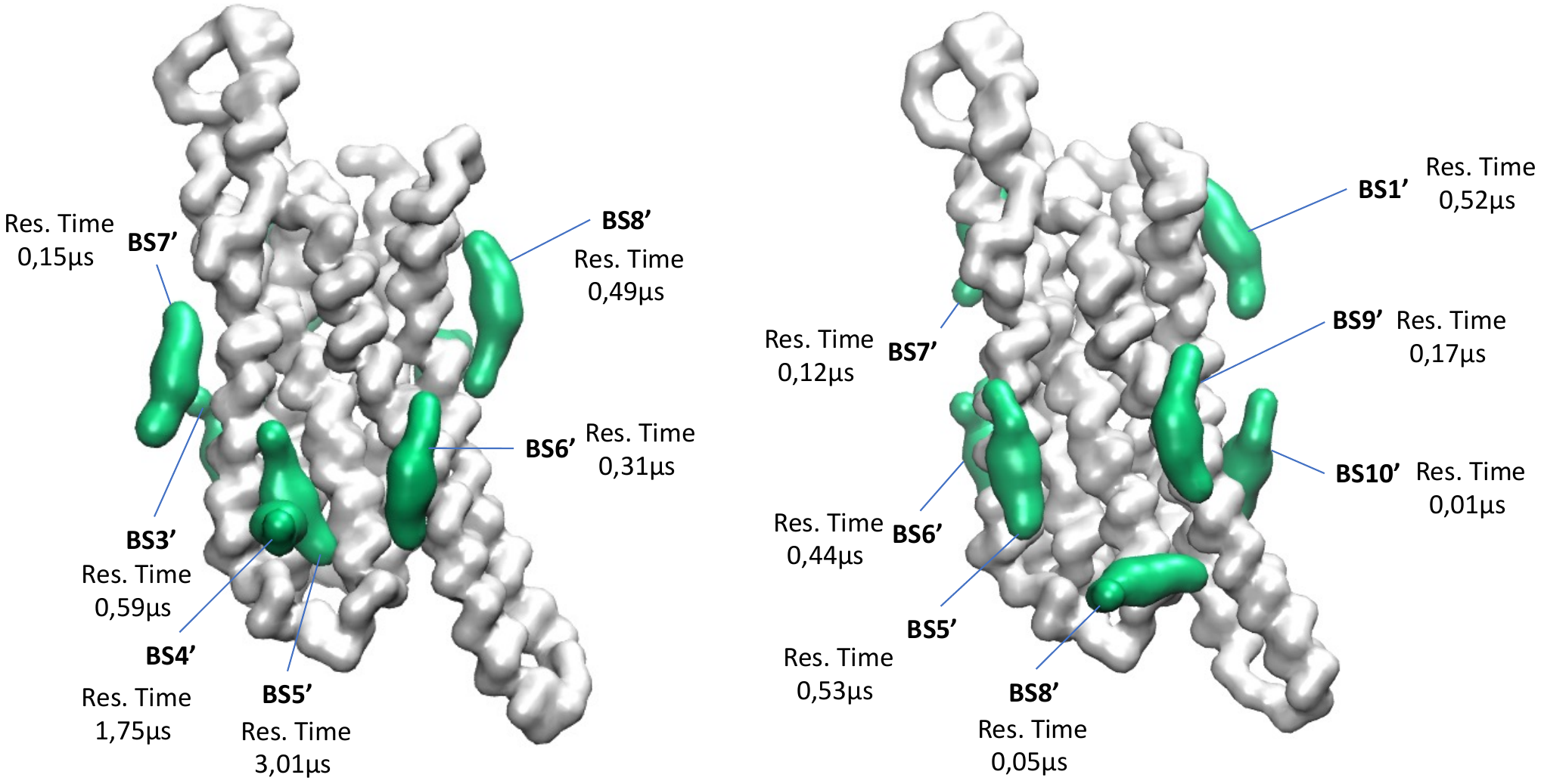

**B**

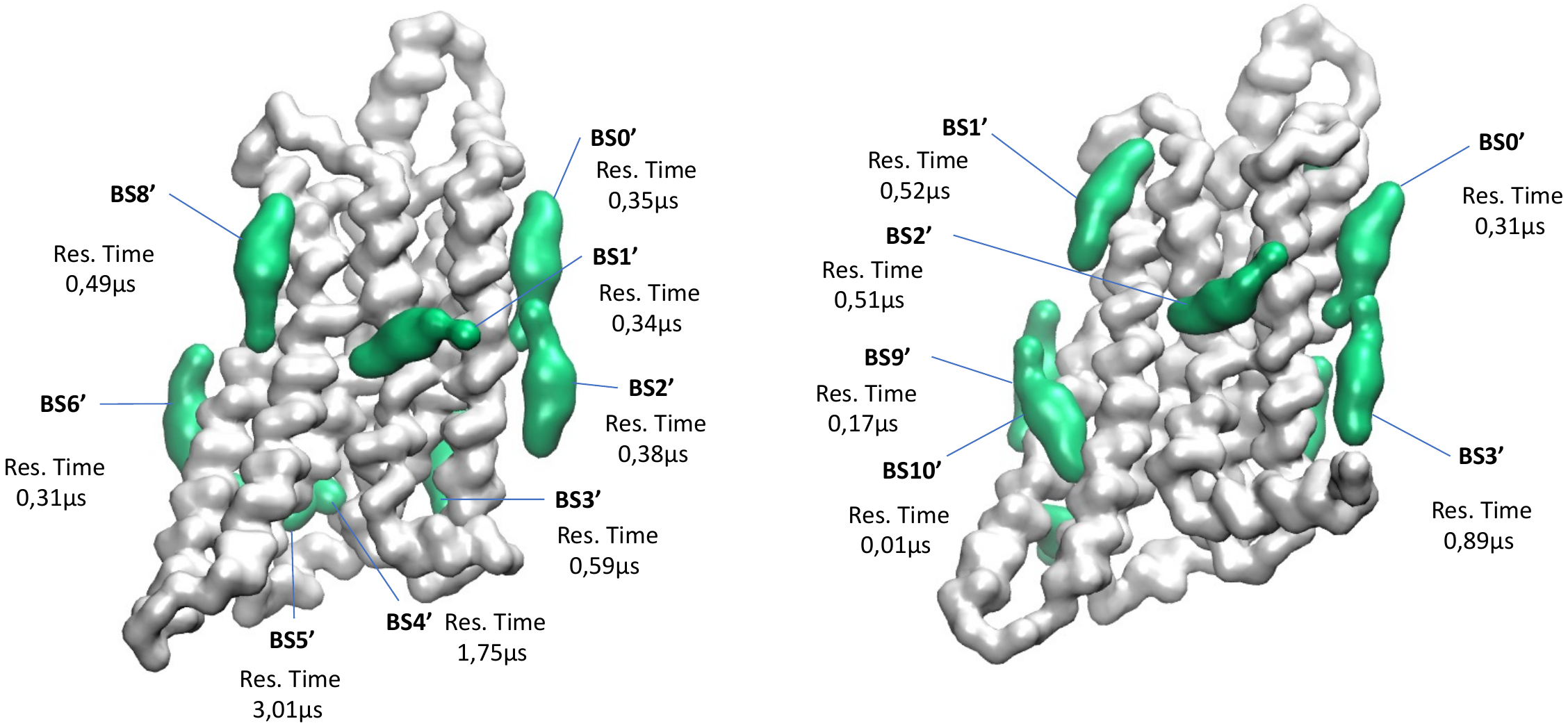

**C**

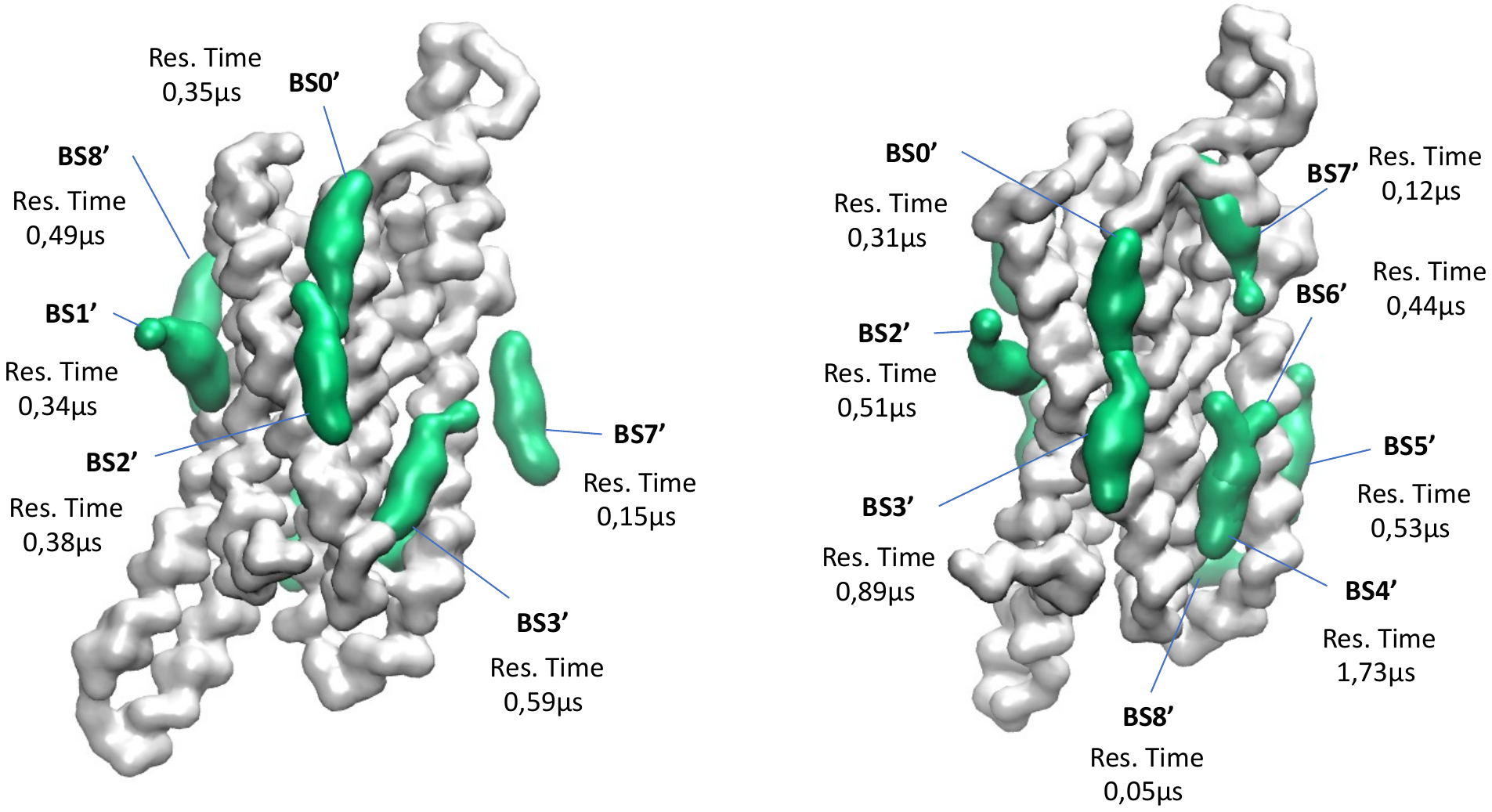

**Supplementary Figure 7.** Comparison of CHOL binding sites between active (BS0′-BS8′) and inactive state (BS0′-BS10′) of A_1_R in POPC‒20%CHOL‒5%PIP_2_ membrane using the last 8 μs of the 10 μs-CG MD simulations with Martini force field ^90–92^ for the analysis of each simulated system calculated by PyLipID. Different sides of the receptor are shown in A, B and C with the active state of the receptor shown in the left side and the inactive the in right side. Receptor is shown in white surface and CHOL representative poses for each binding site in green surface.

**A**

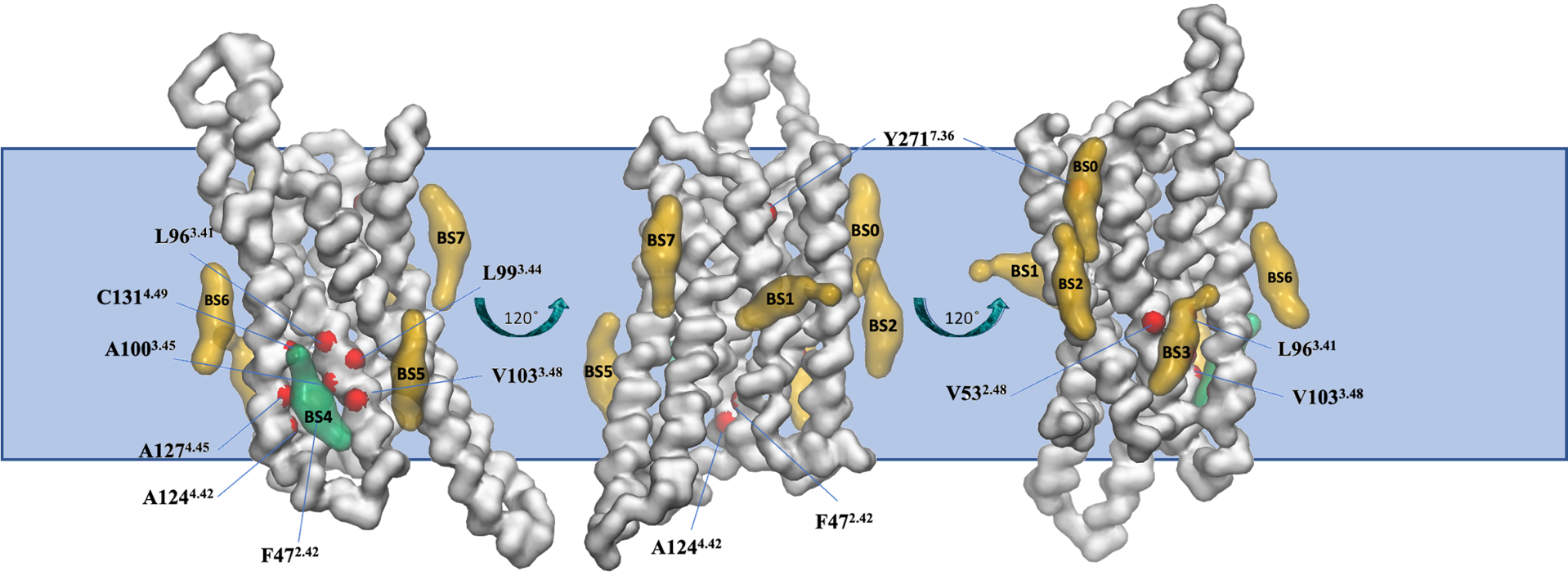

**B**

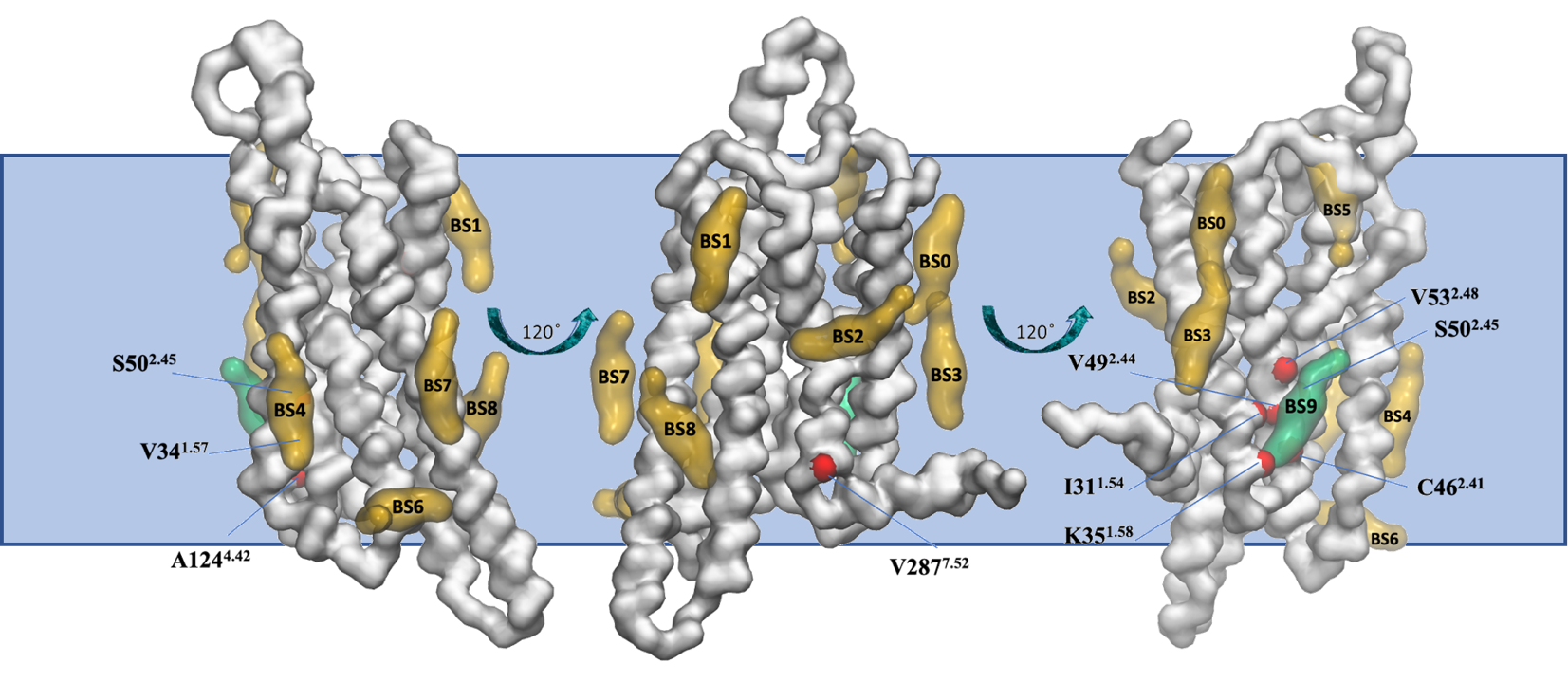

**Supplementary Figure 8.** (A) Binding poses of CHOL in the eight binding sites (BS0-BS7) of the active state of A_1_R in POPC-20% CHOL-5% PIP_2_ membrane. (B) Binding poses of CHOL in the ten binding sites (BS0-BS9) of the inactive state of A_1_R in POPC-20% CHOL-5% PIP_2_ membrane. Binding poses were identified after the analysis of the last 8 μs of the 10 μs-CG MD simulations (three repeats) with Martini force field. Receptor is shown in white surface and CHOL representative binding poses in each binding site are shown in green surface when CHOL residence time in the binding site is more than 1 μs or yellow surface when the residence time of CHOL in the binding site is less than 1 μs. Residues that belong to the identified binding sites with more than 1 μs CHOL residence time οf CHOL are only shown and represented with red surface.

Active and inactive conformation of A_1_R in plasma mimetic membrane

**A**

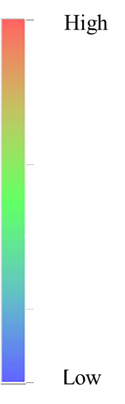

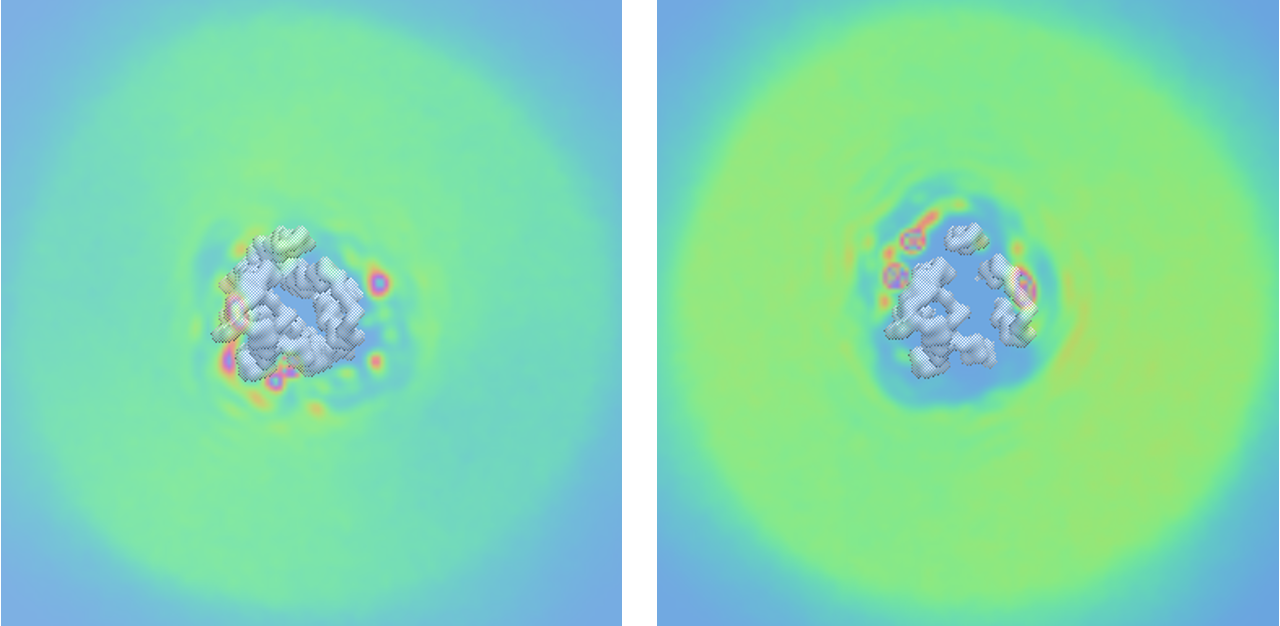

**B**

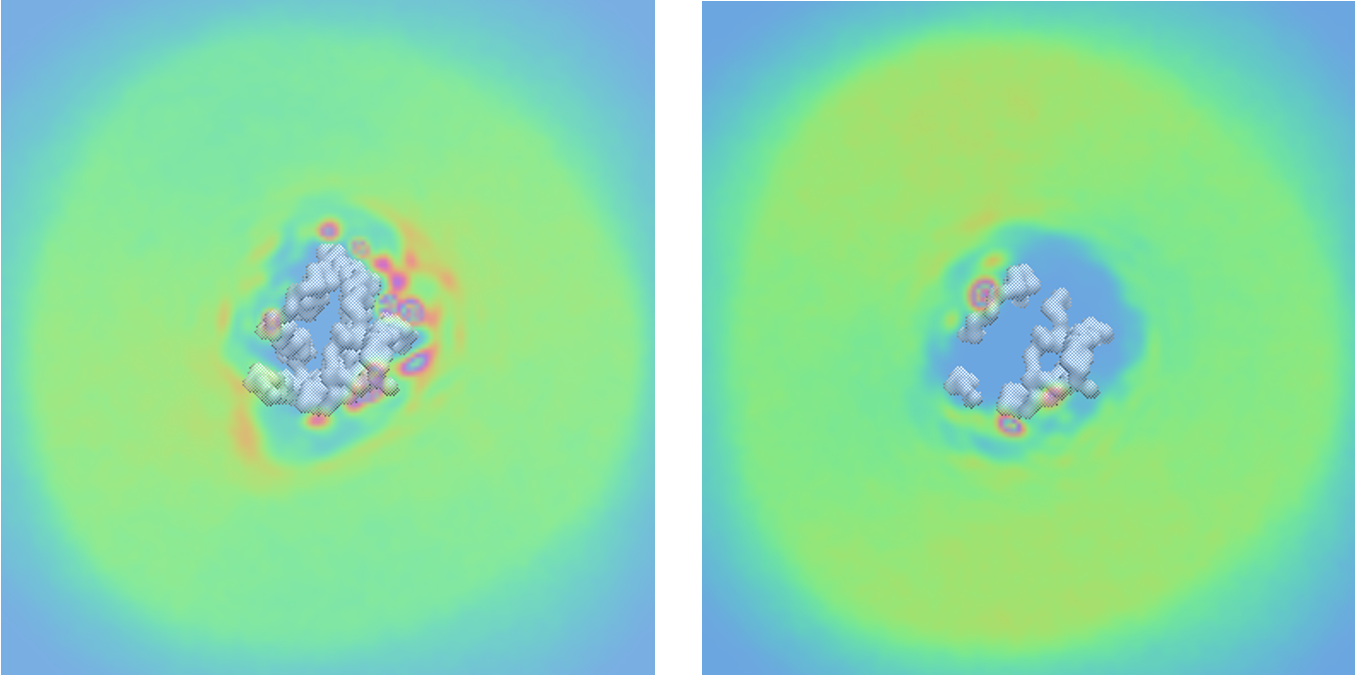

**Supplementary Figure 9.** CHOL density maps averaged for three repeats using ^90–92^ for the analysis the last 8 μs of the 10 μs-CG MD simulations of each simulated system with Martini force field. ^90–92^ (A) CHOL density maps of the active state of A_1_R in plasma mimetic membrane. (B) CHOL density maps of the inactive state of A_1_R in plasma mimetic membrane.

Active conformation of A_1_R in plasma mimetic membrane

The 9 binding sites (BS0’-BS8’) calculated by PyLipID for the inactive state of A_1_R in the plasma membrane, after the visual inspection and grouping together binding sites that lie in proximity, i.e., BS1′, BS2′, BS7′ (BS1) or BS4′, BS5′ (BS2) and BS8′-BS10′ (BS4), ended up 6 distinct binding sites BS0-BS5 shown in Figure 4 which shows residues with residence time > 1μs.

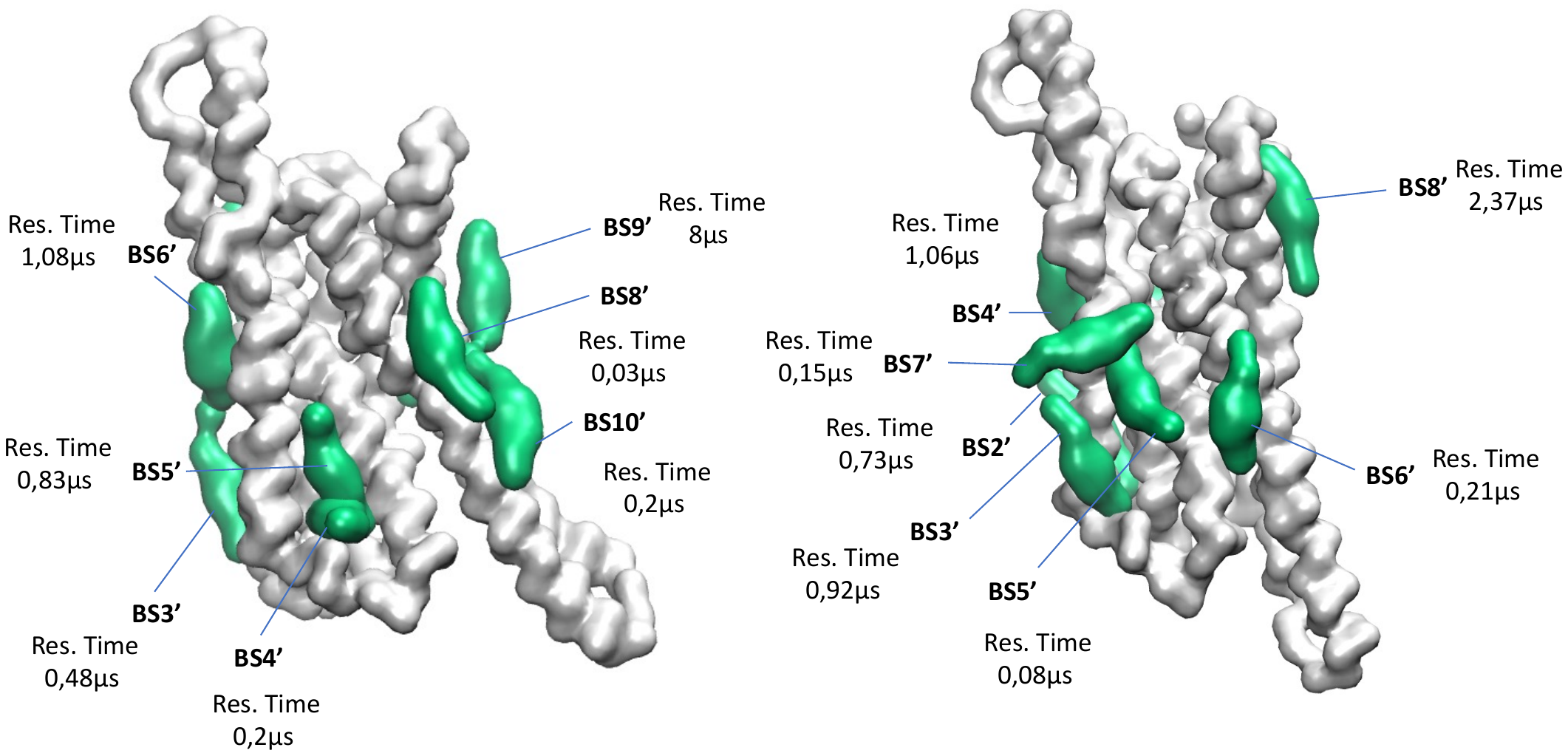

**Α**

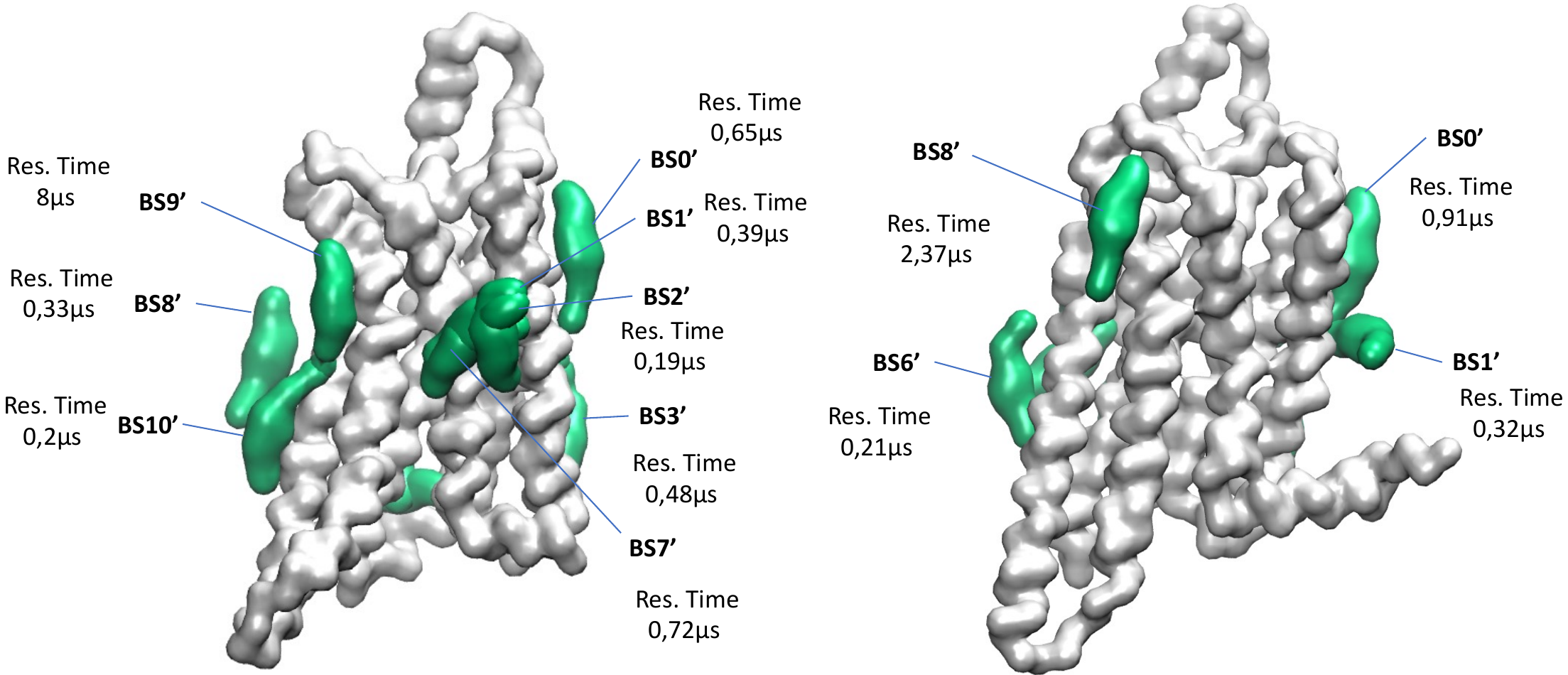

**Β**

**C**

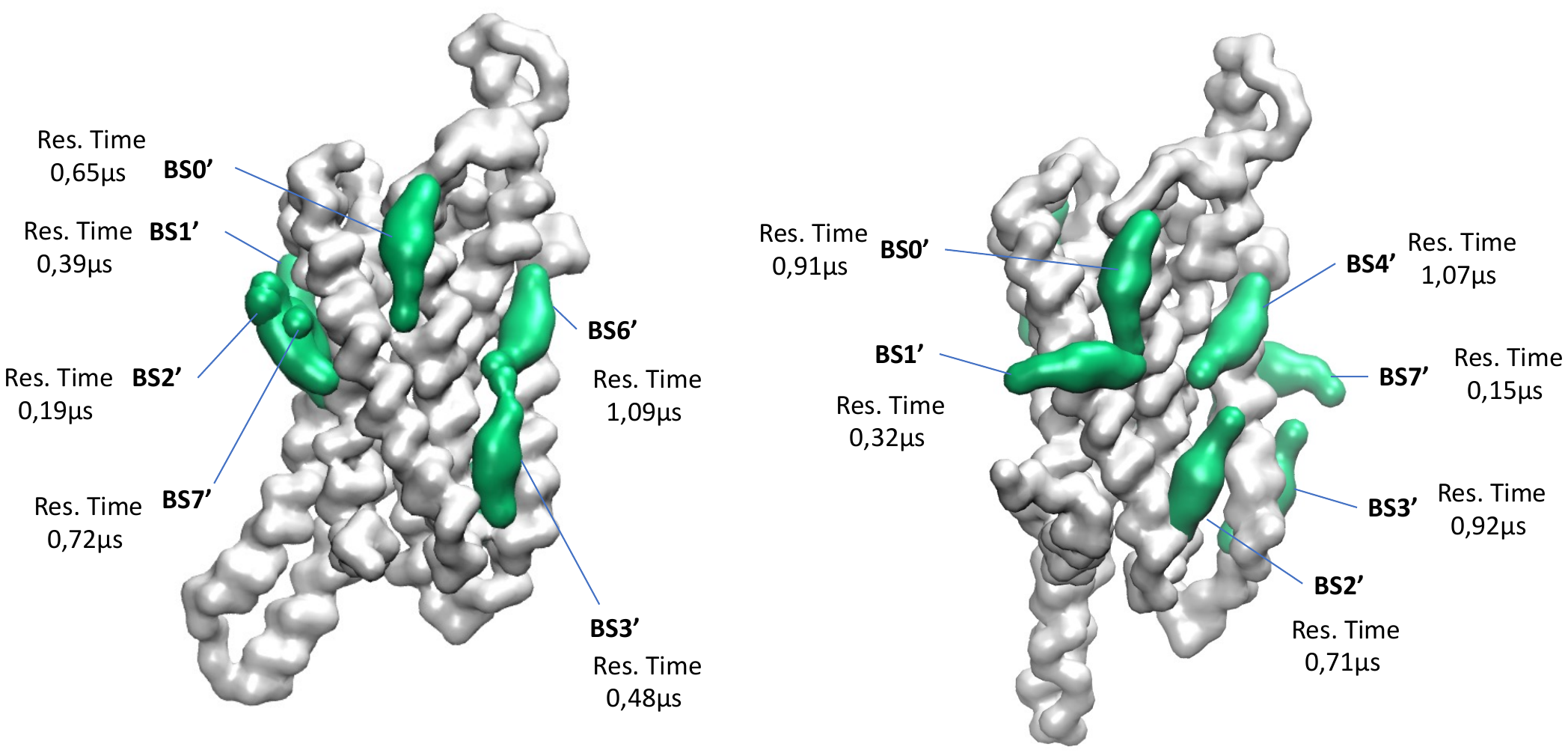

**Supplementary Figure 10.** Comparison of CHOL binding sites between active (BS0′-BS10′, left side) and inactive state (BS0′-BS8′, right side) of A_1_R in plasma mimetic membrane using for analysis the last 8 μs of the 10 μs-CG MD simulation (three repeats) with Martini force field. ^90–92^ Different sides of the receptor are shown in A, B and C with the active state of the receptor shown in the left side and the inactive the in right side. Receptor is shown in white surface and CHOL representative pose in green surface.

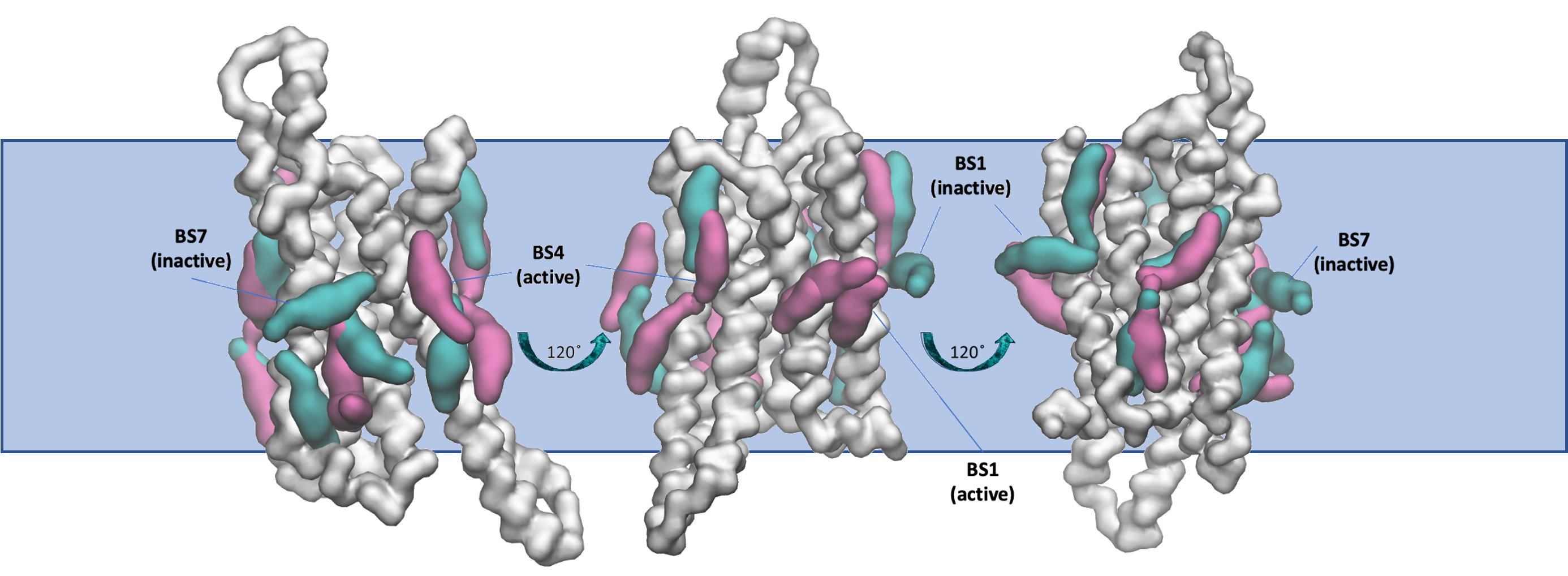

**Supplementary Figure 11.** Comparison of binding poses of CHOL between the active and inactive state of A_1_R in plasma mimetic membrane after the analysis of the last 8 μs of the 10 μs-CG MD simulation (three repeats) with the Martini force field. ^90–92^ The receptor is shown in white surface. CHOL representative poses with more than 1 μs CHOL residence time are shown in magenta surface (active state), or in cyan surface (inactive state).

**Supplementary Table 1.** Lipid composition of the plasma mimetic membrane.

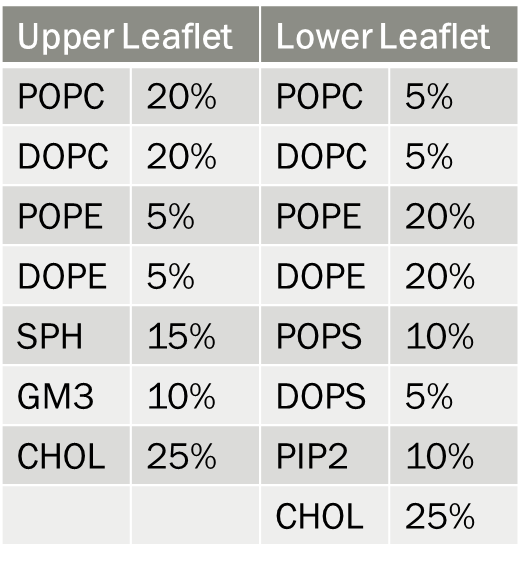

**Supplementary Table 2.** Residence time for the binding sites BS0′-BS9′ of CHOL in the active (left) and for BS0′-BS12′ of CHOL in the inactive (right) A_2A_R conformation in POPC – 20% CHOL membrane averaged for three repeats with PyLipID, ^102,104^ using for analysis the last 8 μs from the 10 μs-CG MD simulation with Martini force field. ^90–92^

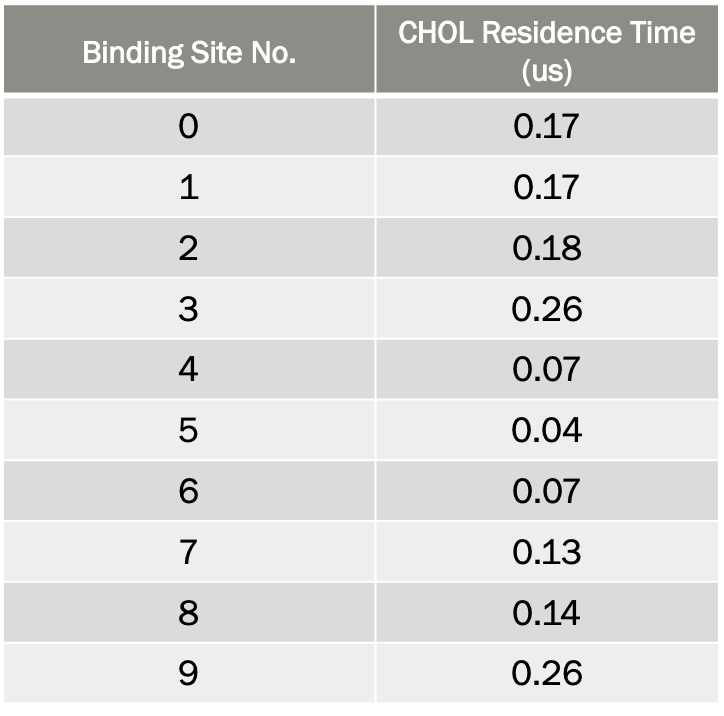

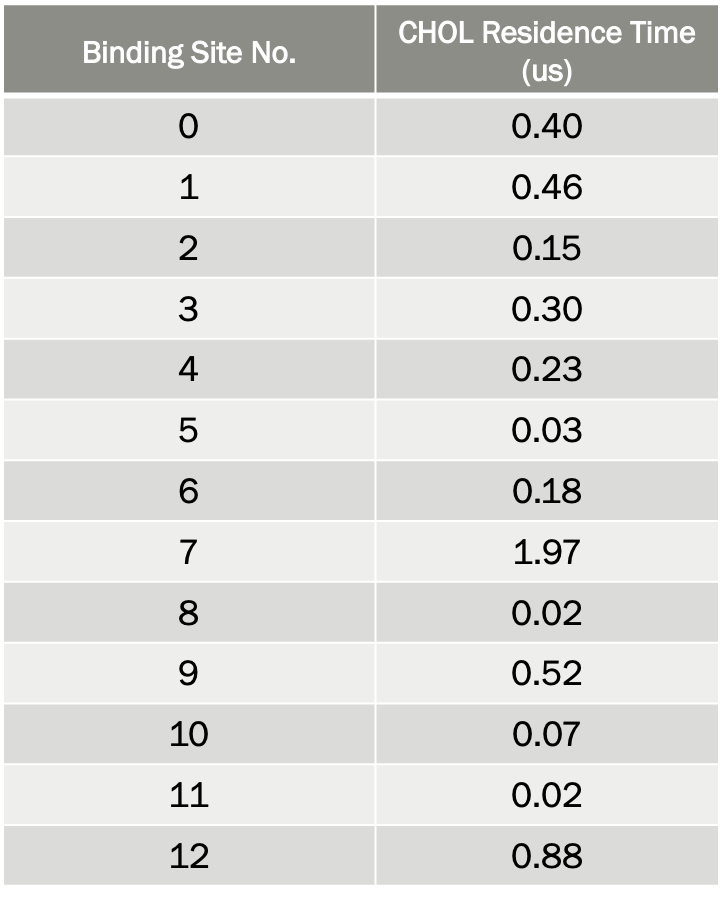

**Supplementary Table 3.** Residence time for the binding sites BS0′-BS10′ of CHOL in the active (left) and for BS0′-BS10′ of CHOL in the inactive (right) A_2A_R conformation in plasma mimetic membrane membrane averaged for three repeats with PyLipID, ^102,104^ using for analysis the last 8 μs from the 10 μs-CG MD simulation with Martini force field. ^90–92^

| 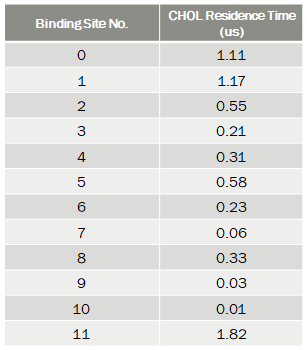 | 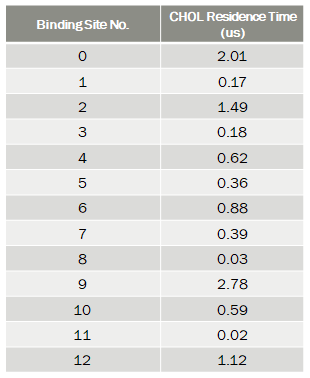 |
| --- | --- |

**Supplementary Table 4.** Residence time for the binding sites BS0′-BS9′ of CHOL in the active A_1_R conformation in POPC ‒ 20% CHOL membrane averaged for three repeats with PyLipID ^102,104^, using for analysis the last 8 μs from the 10 μs-CG MD simulation with Martini force field. ^90–92^

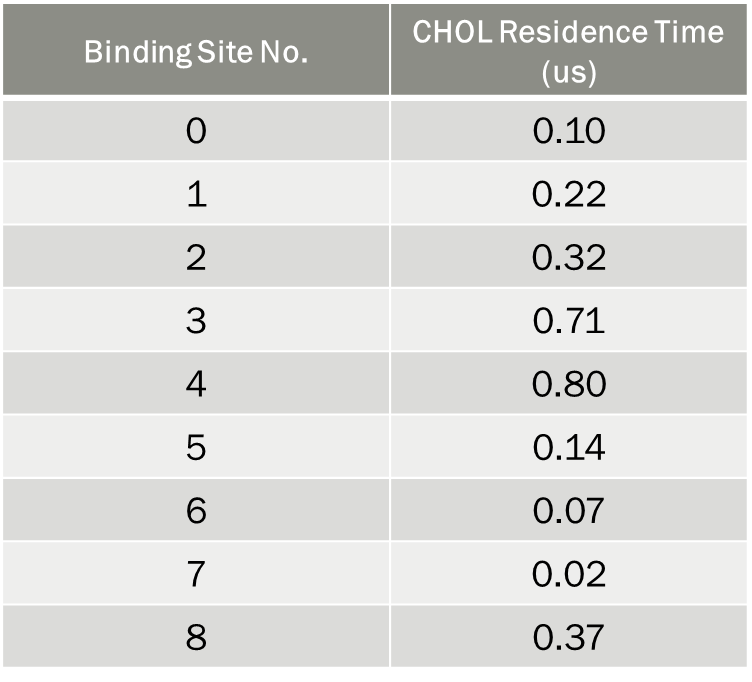

**Supplementary Table 5.** Residence time for the binding sites of CHOL in the inactive A_1_R conformation in POPC ‒ 20% CHOL membrane identified using the last 8 μs of the 10 μs-CG MD simulations of each simulated system with Martini force field ^90–92^ for the analysis with PyLipID. ^102,104^

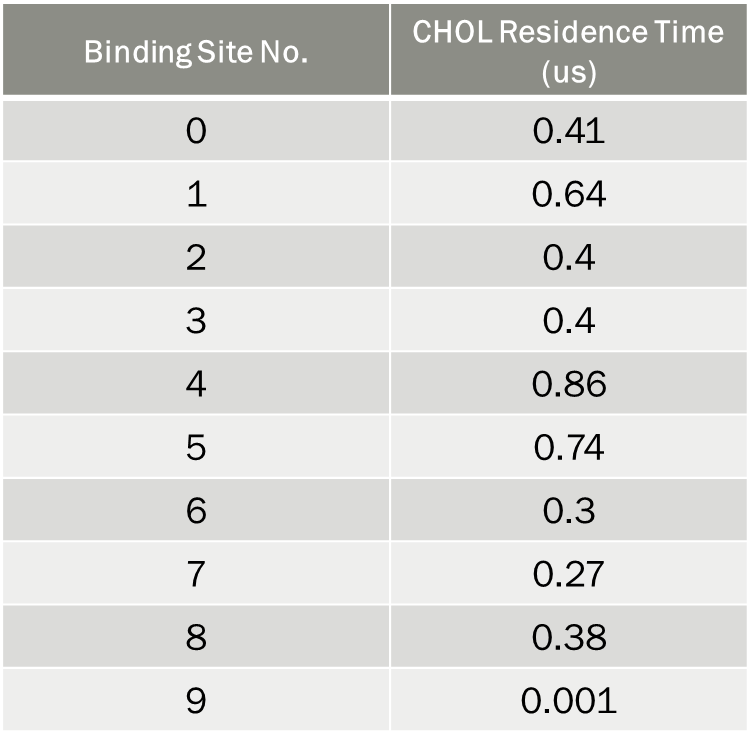

**Supplementary Table 6.** Residence time for the binding sites BS0′-BS8′ of CHOL in the active A_1_R conformation in the membrane composed of POPC-CHOL and 5% PIP_2_ identified using the last 8 μs of the 10 μs-CG MD simulations of each simulated system with Martini force field ^90–92^ for the analysis with PyLipID. ^102,104^

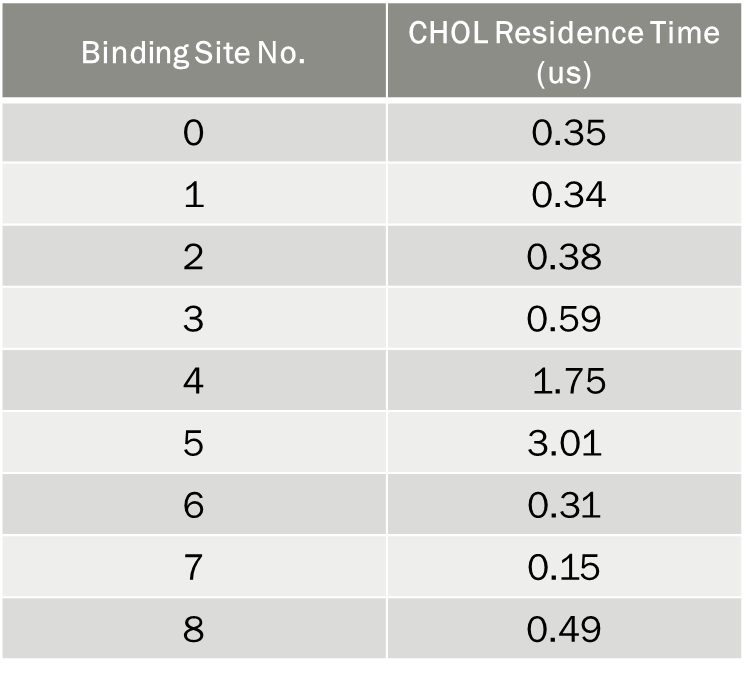

**Supplementary Table 7.** Residence time for the binding sites of CHOL in the inactive A_1_R conformation in the membrane composed of POPC-CHOL and 5% PIP_2_ identified using the last 8 μs of the 10 μs-CG MD simulations of each simulated system with Martini force field ^90–92^ for the analysis with PyLipID. ^102,104^

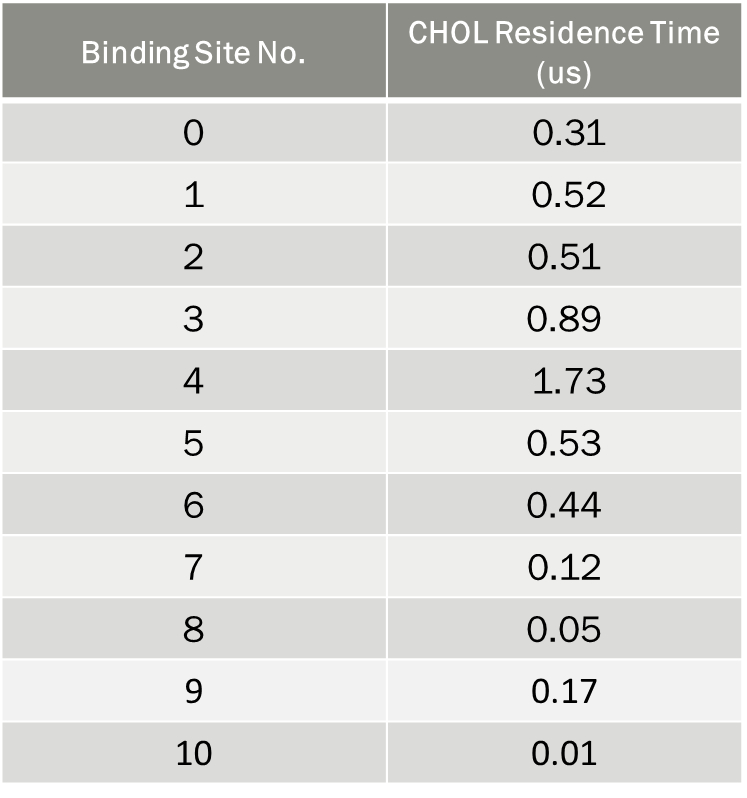

**Supplementary Table 8.** Residence time for the binding sites BS0′-BS10′ of CHOL in the active A_1_R conformation in plasma mimetic membrane identified using the last 8 μs of the 10 μs-CG MD simulations of each simulated system with Martini force field ^90–92^ for the analysis with PyLipID. ^102,104^

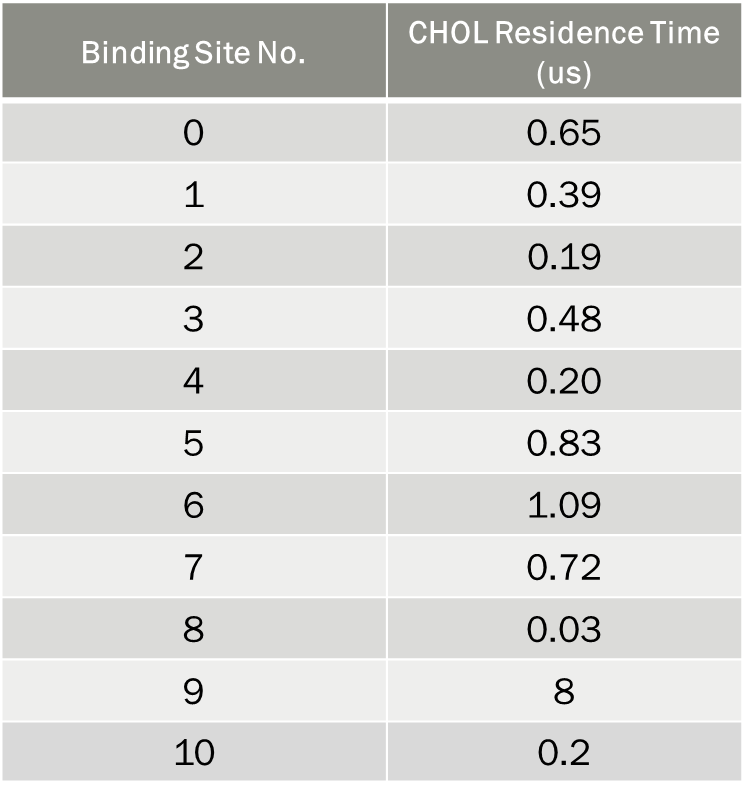

**Supplementary Table 9.** Residence time for the binding sites of CHOL in the inactive state A_1_R in plasma mimetic membrane identified using the last 8 μs of the 10 μs-CG MD simulations of each simulated system with Martini force field ^90–92^ for the analysis with PyLipID. ^102,104^

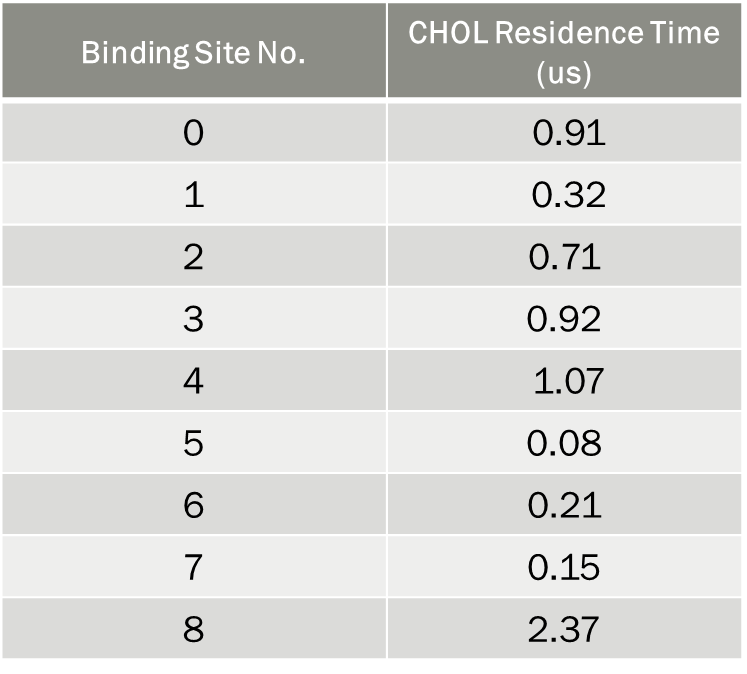
